# Quantum Chemistry in a Pocket: A Multifaceted Tool to Link Structure and Activity

**DOI:** 10.1101/2024.04.22.590615

**Authors:** Filipe Menezes, Valeria Napolitano, Tony Fröhlich, Sarah Rioton, Julian D. Janna Olmos, Grzegorz Dubin, Grzegorz M. Popowicz

**Affiliations:** Helmholtz Munich, Molecular Targets and Therapeutics Center, Institute of Structural Biology, Ingolstaedter Landstr. 1, 85764 Neuherberg, Germany; Malopolska Centre of Biotechnology, Jagiellonian University, Gronostajowa 7a, 30-387 Krakow, Poland

## Abstract

We introduce In-Pocket Analysis, a simple and efficient protein-ligand complex structure optimization algorithm. It provides structural biology and structure-based drug discovery with a much-needed tool for data curation and quantification analysis. In-pocket analysis removes unphysical tension from experimental structures, yielding high-quality atomic models. This includes deformed bonds, structural clashes, proton refinement, and solvent envelope. The algorithm is compatible with quantum methods or force fields, delivering precise calculations of binding energies and QSAR. Its applications include refinement of experimental structures, calculation of inter- and intramolecular energetics, scoring of docked poses, scaffold decoration/hopping, or building of explainable QSAR.

For validation and benchmarking we used high-resolution crystal structures of the N-terminal domain of PEX14 with small-molecule inhibitors. PEX14 is a target candidate against Trypanosomiasis and Leishmaniasis. The application of our algorithm allowed for an explanation of unexpected binding poses, rationalization of binding energies, and affinities as well as precise quantification of solvent-mediated interactions.

**TABLE OF CONTENTS:** 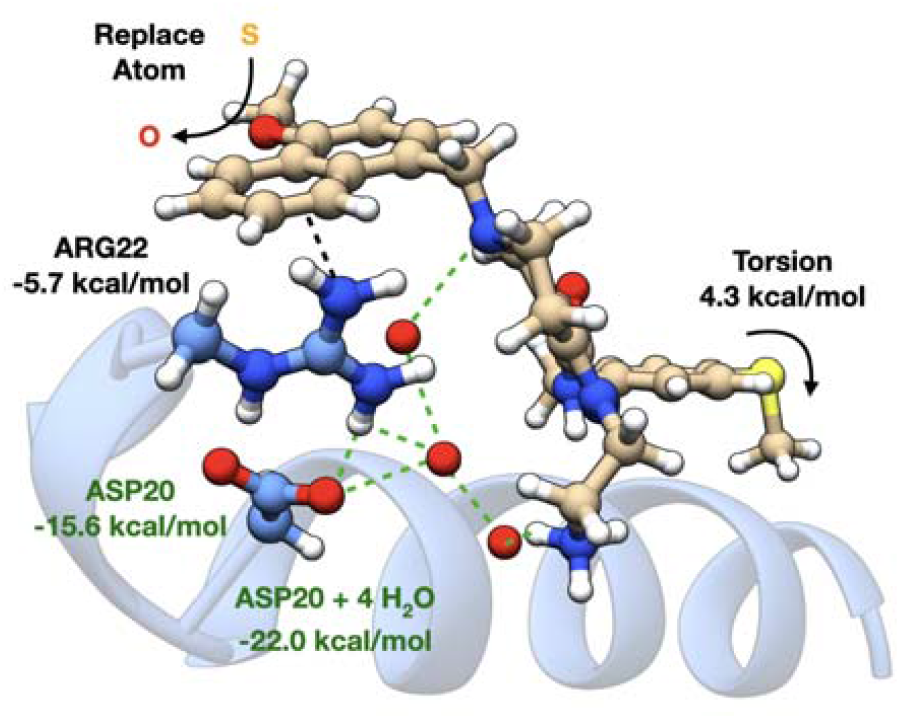

## INTRODUCTION

The existence of stable complexes between ligands and proteins is essential for medical and life sciences, especially in medicinal chemistry, chemical biology, and biochemistry.^1^ Understanding how a ligand affects and interacts with a protein requires determining the structure of the respective complex, *i.e*., identifying the relative coordinates of all atoms in the system. This enables analysis of interactions, followed by fine-tuning of the ligand for optimal binding, constituting the basis of structure-based drug discovery (SBDD).

Building atomic models of protein-ligand complexes is achieved indirectly from experimental data since atomic coordinates are not directly observable. For instance, structures determined by X-ray crystallography are obtained from diffraction patterns.^2-9^ The quality of this model is strongly impacted by the crystal’s degree of order: low-order lattices lead to poor diffraction patterns, which result in low-resolution structures. This is mitigated through geometrical constraints of free molecules, which bias the structure towards ideal fragment/residue geometries. Since they may not be representative of the bound system, errors and unphysical strain are introduced in the structural model. Furthermore, electronic densities are not always unambiguously interpreted. Even in high-resolution structures, the orientation of imidazole rings in histidine residues may only be retrieved indirectly by weighting different hydrogen bond (H-Bond) networks. The orientation of asparagine or glutamine residues requires similar considerations. Isotropic B factors, rather than anisotropic, or human bias contribute even further as sources of error. Particularly sensitive is the situation for hydrogen atoms.^10-12^ For structures of insufficient resolution (>1.2 Å), the electronic density cannot reflect the presence of these atoms, and models are built without their explicit coordinates. These considerations affect the reconstructed structure of the protein and ligand similarly. The resulting proton-less structures may be corrected by rule-based algorithms that determine atomic hybridizations, possibly also *pK*_*a*_s, and estimate how many protons should be added until the octets are completed (for a few examples refer to the literature^13-18^). This task is well standardized for protein residues, where structural variability and the effects of chemical environments are somewhat limited. Still, low resolution on protein structures leads to inaccuracies in the actual positions of the protons, not to mention the ambiguities in assigning protonation states to internal residues.^19,20^ For the small molecule ligand the situation is less straightforward and countless other factors may come together to yield even chemically inconsistent structures.^21^ The propagation of errors due to unphysical strain and structural inconsistencies has an impact at many levels. Deformation energies are overestimated.^22^ This leads to an unrealistic over-stabilization of the ligand by the protein and calculated interaction energies (enthalpies) are incorrect. It also renders electrostatics wrong since atomic charges are determined by structure. It is therefore important to have the best possible structures for the most faithful representations of the protein-ligand complexes, and this can only be accomplished by optimization to remove unphysical biasses.^23,24^

All components, including the solvent, are participating in the binding event. Consequently, they all influence the landscape of the Potential Energy Surface (PES). It is not uncommon for proteins to favor higher energy conformers of the free ligand.^25^ Therefore, obtaining the real complex conformation demands consideration of all these effects. While the inclusion of the full protein is manageable at the molecular mechanical (MM) level, the computational costs become prohibitive in quantum mechanical (QM) treatments. At semi-empirical levels of theory, the full inclusion of the protein in the geometry optimization leads to time and resource-intensive calculations. If higher-level quantum chemistry is required due to the ligand’s electronic structure (*ab initio* methods), including just a few residues may have catastrophic effects on calculation times. It is therefore desirable to minimize the number of protein atoms entering the calculation for optimal efficiency, while minimally sacrificing accuracy. Unfortunately, from the geometry optimization point of view, removing non-pocket residues strongly impairs the stability of the resulting complex, not to mention the nature of the interaction itself.

The concept of using QM to improve the quality of protein-ligand structures is not new to SBDD. Merz and coworkers defined a QM/MM modification of the crystallographic structure refinement protocol.^10^ This allowed using high-level quantum chemistry to refine experimental structures, though with the limitations of the QM/MM method.^26^ A different procedure was proposed by Perola and Charifson,^25^ and later by Merz and by Stewart,^27-29^ who biased the PES with harmonic potentials towards the experimental geometry. However, the choice of parameters for harmonic potentials is not straightforward, which rendered the technique cumbersome. Further, the increased risk of contaminating the structures with inaccuracies of the underlying methods was large.^25,29^ Biasing is also widely employed in the field of molecular dynamics. Dating back to the work of Torrie and Valleau in umbrella sampling^30^ this concept fully flourished with Laio and Parrinello when metadynamics was introduced.^31^ Metadynamics constructs a bias potential based on Collective Variables (CVs). Grimme introduced Cartesian RMSDs as CVs for QM-based metadynamics,^32^ which was used for designing several algorithms like the CREST conformer search^33,34^ and the single-point Hessian.^35,36^ However, the use of CVs in biasing for experimental structural refinement or in computational SAR studies remains unexplored.

A seemingly unrelated problem is that of ligand docking to a protein.^37^ Docking protocols aim at predicting the binding pose of a ligand in the protein’s pocket using a scoring function, a generalization of the concept of (Gibbs free) energy. Several docking programs are popular in drug discovery and differ conceptually in how they treat the system.^38-52^ In Schrödinger’s Glide^45^ for instance, most constraints are filters, and consequently they do not affect directly nor guide the optimization/determination of the binding pose: constraints are used predominantly as guidelines to disregard certain binding poses from further processing. Beyond ligand-only flexibility, the InducedFit protocol^53-55^ of Schrodinger introduces protein flexibility. This, however, takes the form of a series of subsequent docking and optimization steps. The residues considered for the flexible docking are selected based on distance criteria (concerning the ligand) or are hand-picked by the user. The inaccuracies and simplifications used limit the accuracy for the benefit of speed.

Here, we propose mitigating these problems by extending the use of Cartesian RMSD CVs to emulate the constraining effects of a deliberately omitted part of a large molecular complex (Figure 1a). For example, the omitted part can be the whole receptor, enabling ligand-specific properties to be explored by high-level quantum chemistry. Or it may be higher shells of residues around the ligand. This is accomplished with an RMSD-based Gaussian potential that modifies the molecular PES to lock the experimental conformation while relaxing tension accumulated in bonds and angles. The methodology, called in-pocket analysis (IPA), is validated for several applications in drug discovery: *i)* removing unphysical strain (distortion) from experimental structures for protein-induced ligand-strain analysis; *ii)* understanding protein-ligand interactions by isolation of specific target groups; *iii)* SAR/scaffold hopping studies where a ligand-pocket complex is held adaptatively immobile and a constraint-free fragment of the ligand is modified; *iv)* structure refinement for binding energy or affinity calculation; *v)* docking. More applications can easily be envisioned. Different degrees of flexibility for different molecules or parts of the same molecule are possible. While CVs have previously been used in docking, their role was, to the best of our knowledge, restricted to enhance conformational sampling,^51,52^ and not to guide the docking protocol. Much like other applications of CVs, the technique may be mixed with MM and QM methods alike.

**Figure 1.**
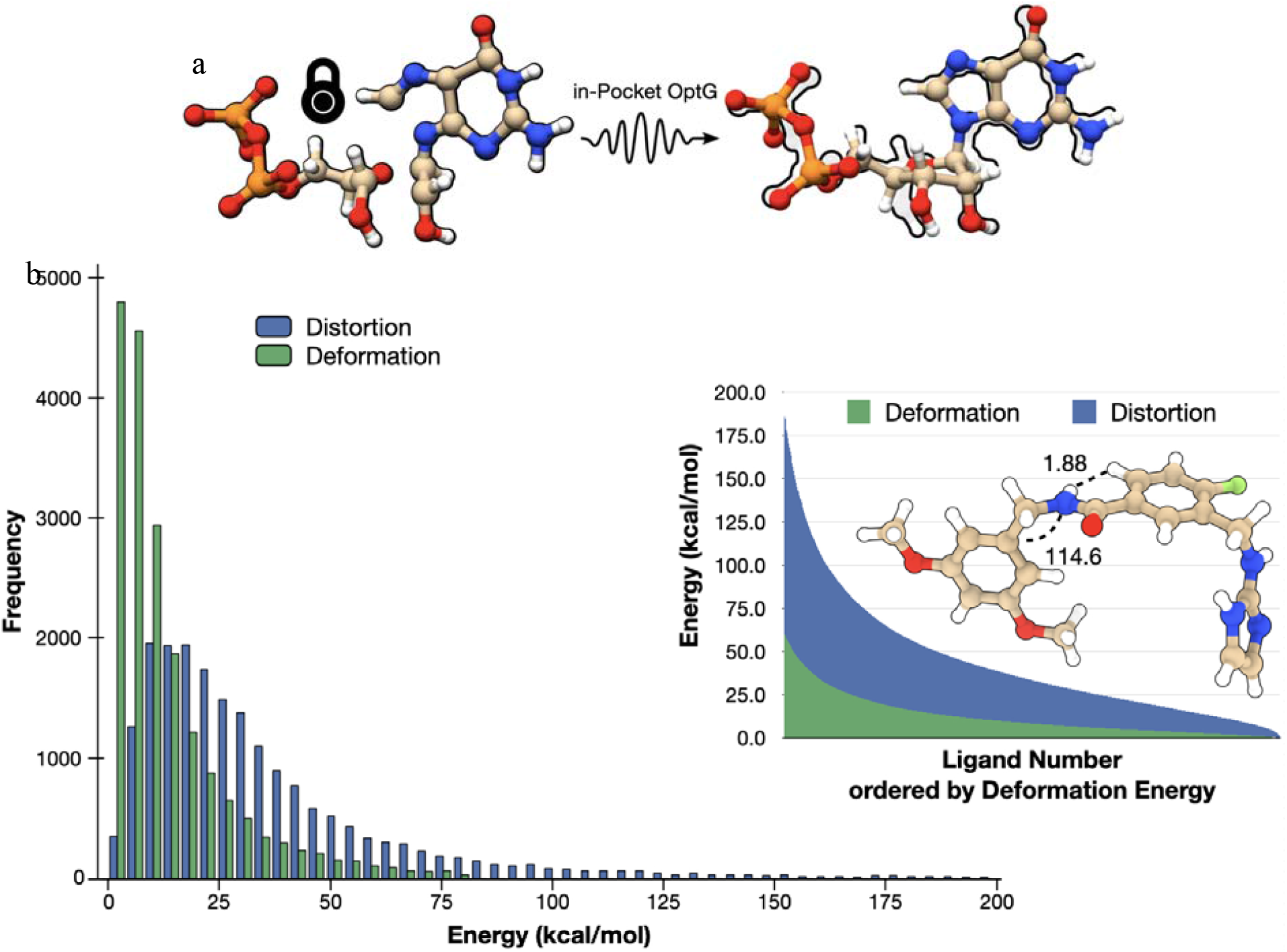
In-pocket optimization, a Collective-Variable modified unconstrained geometry optimization with applications in drug discovery. a) By applying a Gaussian bias potential to lock the conformation of a starting structure, unphysical tension on bonds and angles is effectively removed from experimental structures. b) The distributions of deformation and distortion energies for ligands in the MISATO database.^21^ Example of ligand in PDB ID 6E22, where an angle and a distance are major contributors to distortion.

Though useful, three-dimensional structures, fail to provide clear guidance in hit-to-lead optimization. This is due to a lack of reliable methods to quantitatively characterize interactions, solvent participation in binding, or crystallization artifacts, but also due to structural inaccuracies that remain unaccounted for. We distinguish unphysical distortion from protein-induced strain, deformation, or tension, which we consider chemical or biologically motivated. Therefore, in what follows, we reserve the concept of structure distortion to inaccuracies caused by the limited resolution of experimental techniques, whereas tension, deformation, or strain are interchangeably reserved for conformational changes required for binding to occur. Figure 1b shows histograms for distortion and deformation energies for the ligands in the MISATO database.^21^ Distortion energies were calculated after IPA refinement. The distribution of distortions is broad and flat. This distribution’s peak ranges distortion energies of 11.1-18.3 kcal/mol, and the long tail was truncated at 200 kcal/mol. Deformation energies peak at 3 kcal/mol and rapidly decay. An alternative view is offered in Figure 1b, where deformation and distortion are compared for the same molecules over the whole database. The latter is systematically 2-4 times larger than the former, indicating a fundamental problem in the molecular structures on which drug discovery and chemical biology are based on. Though the large values recorded stem primarily from the incorrect placement of protons, this problem reflects underlying inaccuracies in the molecular scaffolds. The structures and properties of the refined ligands are provided as a supplement database to this manuscript.

We primarily validate our method using experimental SAR and binding data for Protein-Protein Interface (PPI) inhibitors. An X-ray structure of *Trypanosoma brucei* (*Tb*) PEX14 in complex with compound **1** is the main case study (*cf*. Figure 2a). The PEX5-PEX14 interface offers a therapeutic door against *Trypanosoma* infections.^56^ Yet, this is a difficult drug target due to typical PPI features, like its largely lipophilic nature and absence of deep pockets or polar interactions. The water network around the PPI has been also recognized as critical for binding.^57^ We show how IPA can explain the SAR of PEX14 inhibitors^57^ and rationalize their mode of action.

**Figure 2.**
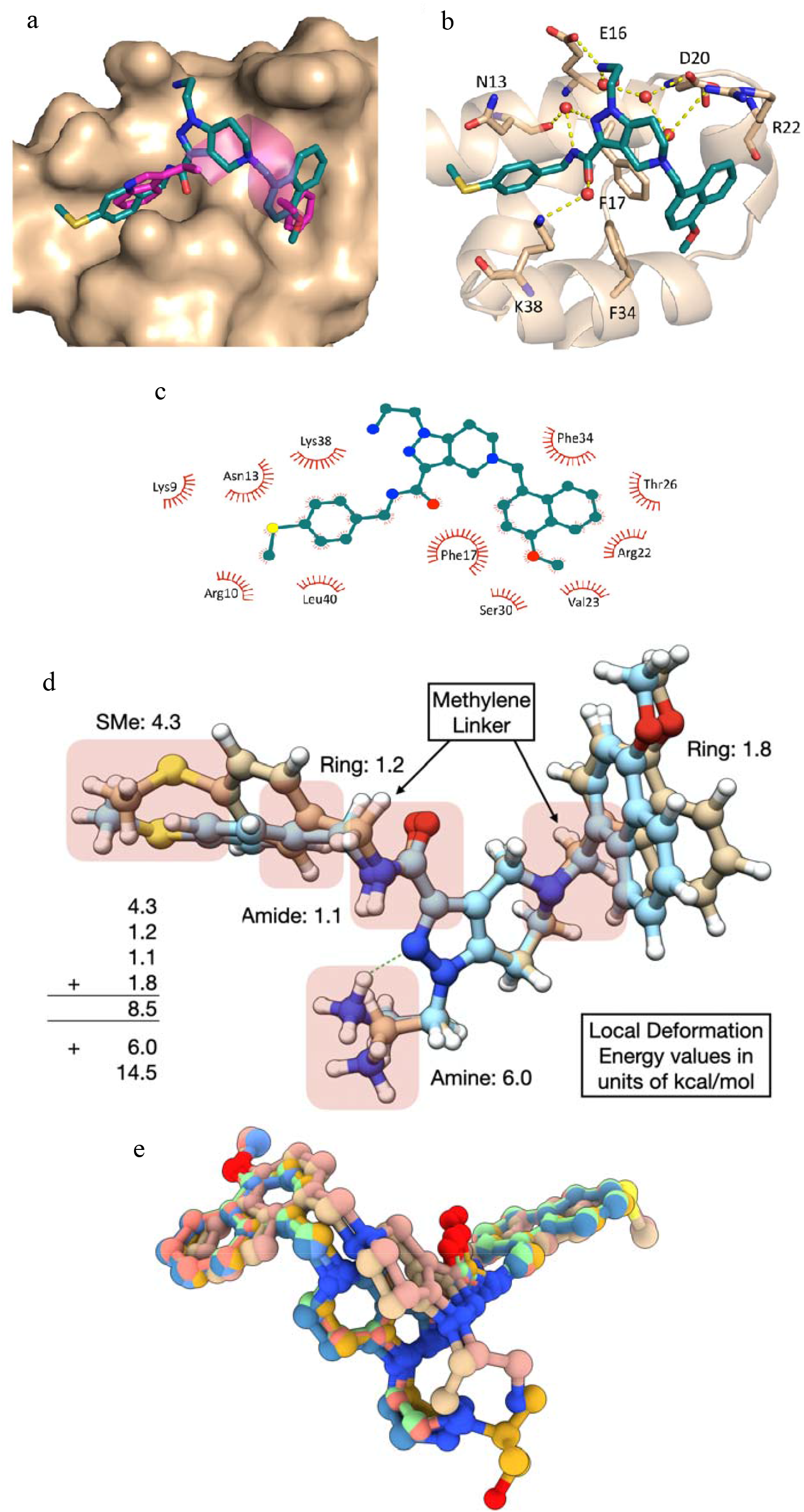
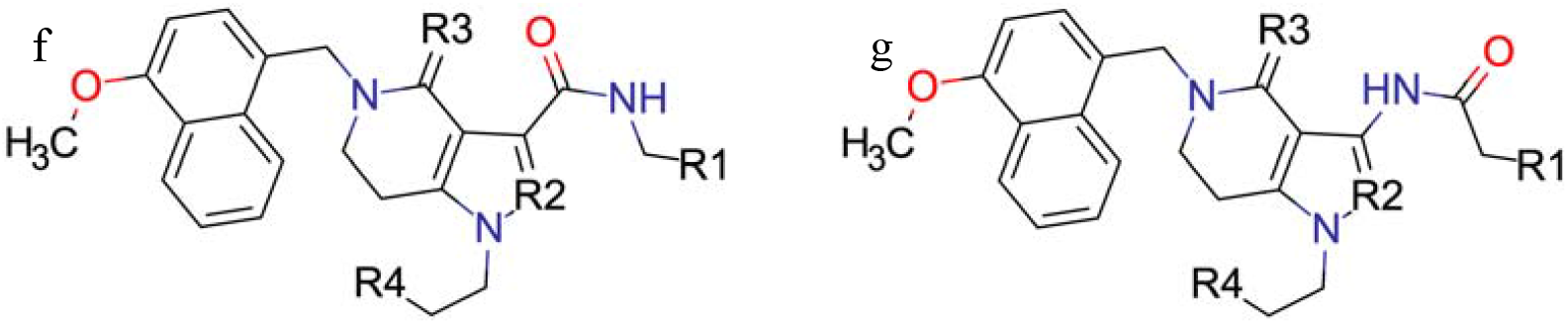
a) Surface representation of *Tb*PEX14. Compound **1** (green sticks) binds at the PEX14-PEX5 Protein-Protein interaction and mimics the binding pose of the WxxxF motif from PEX5 (magenta; PDB code: 2W84^59^). **b) Complex structure of water-mediated, polar interactions between *Tb*PEX14 and compound 1. c) 2D molecular interaction plot of the hydrophobic contact between TbPEX14 and compound 1. d) The structure of compound 1 and the respective tension analysis (in kcal/mol)**. The latter was obtained by comparing the experimental structure of the bound ligand (light brown) against the freely optimized structure (light blue). The green, dashed line represents an intramolecular H-Bond. **e) Overlay of binding poses for compound 1 (light brown and salmon), the ligand in 6SPT (gold), and the ligand in 5N8V (blue, pink, green)**. The change in conformation for the pyrazole-[4,3-c]pyridine scaffold is evident, especially due to the conservation of the position of the aromatic rings and of the primary ammonium group (in 5N8V ligands). Based on classical structure analysis there is no explanation for the observed differences. **f) General scaffold summarizing the variations of the original ligand (1) considered in this work. g) Structure of the amide with inverted orientation**. For specific structures, see Table 1.

## RESULTS

In this work, we modify the PES of a molecular system using a bias potential of the form

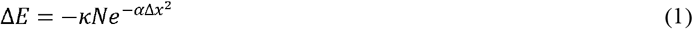

**Table 1.**
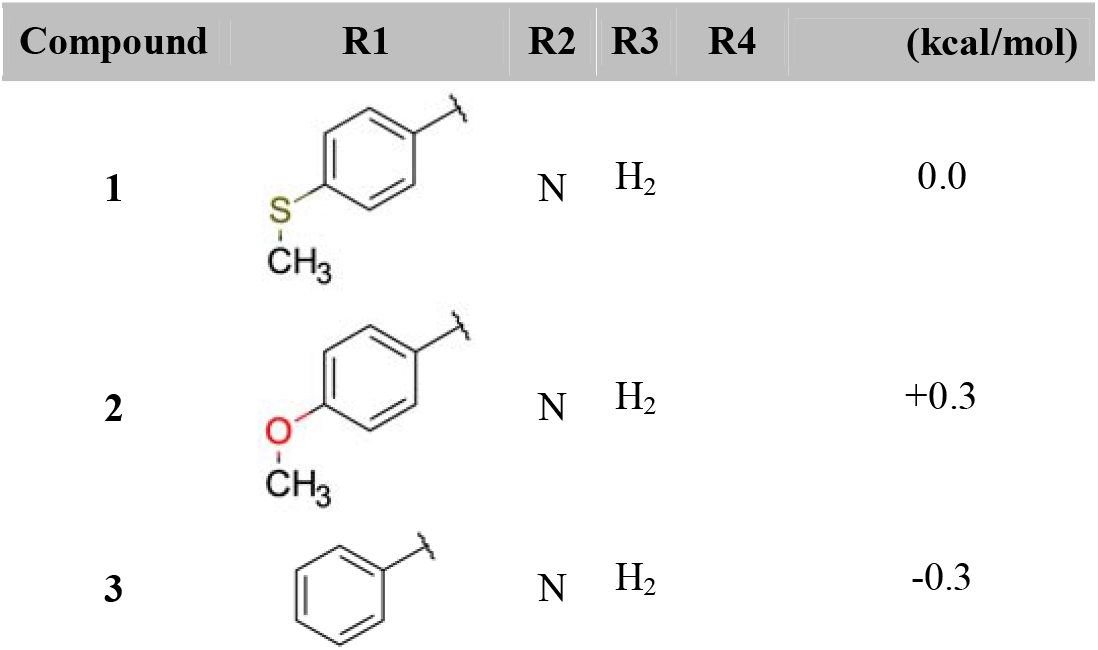
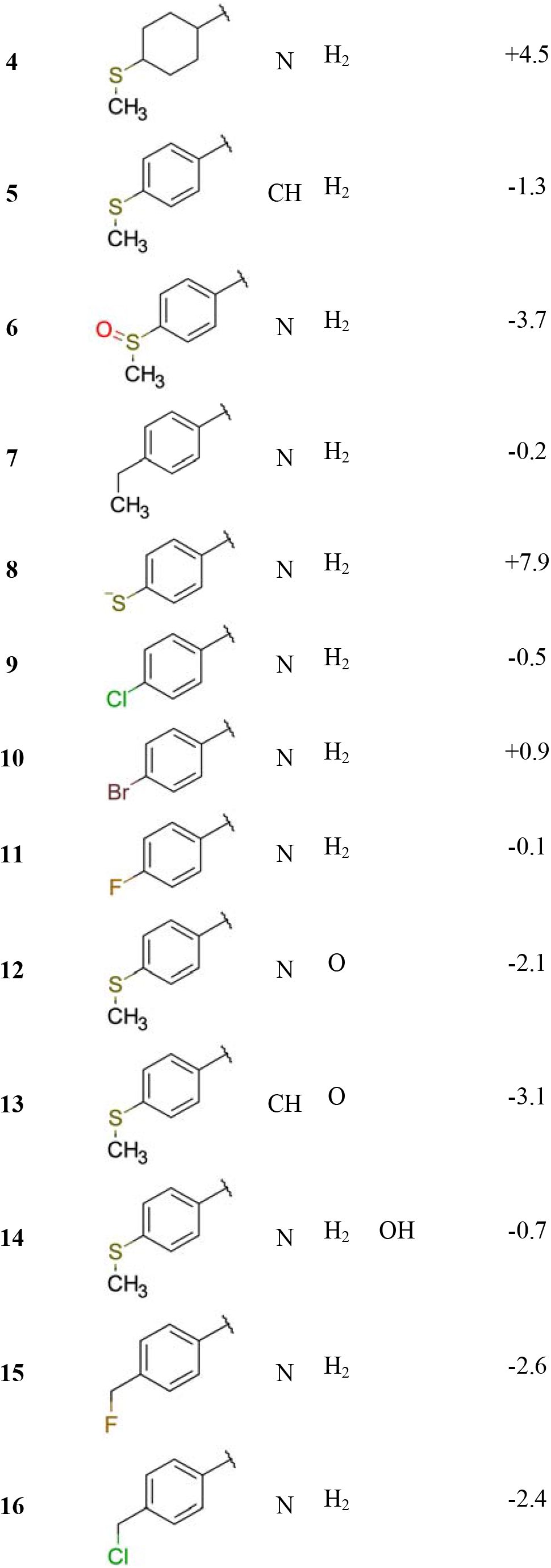
Substitution patterns and binding energies () for the several *in-silico* modifications of compound 1.

The RMSD, Δ*x*, is constructed from an experimental geometry, which is the target conformation of the molecule. *κ* and *α* are parameters in the model potential. We consider only positive *κ* and *α*, such that the bias potential Δ*E* is attractive towards the reference structure. This contrasts with the typical use of CVs, where bias potentials are repulsive. *N* is a factor that we take as the total number of atoms used for calculating the RMSD. This setup permits running constrained minimizations while exploiting the efficiency of unconstrained optimization algorithms. In the Supplementary Material (S1) such an example is provided for a Lennard-Jones function. Ideally, *κ* should be in the range of 15-30 kcal/mol, whereas *α* should take values within 0.5 and 1.0 Å^−2^ (see S2-S8). This allows a very quick and systematic optimization of structures while retaining conformation: bond distances and angles are the primary variables optimized. This is key for reducing the distortion accumulated due to the limited resolution of experimental techniques.

In-pocket optimization is available from the ULYSSES suite.^58^ The source code is available from https://gitlab.com/siriius/ulysses.git.

### A complex of PEX14 with compound 1: analysis of the structure

We co-crystallized *Tb*PEX14 with compound **1** (Figure 2a). The crystal diffracted at 1.1 Å resolution and belongs to the space group C121, with two molecules in the asymmetric unit. The crystal structure reveals that the inhibitor targets the PEX5 binding interface. The binding surface consists of two hydrophobic sub-pockets separated by *π*-stacked phenylalanines (Phe17 and Phe35) and the inhibitor is found mimicking the binding pose of PEX5’s WxxxF motif (Figure 2a).^59^ The pyrazole-[4,3-c]pyridine scaffold of compound **1** is located on top of the phenylalanines, whereas the hydrophobic cavities are filled by the methylsulfonylphenyl and methoxynaphthalene groups which decorate the central scaffold. The methylsulfonylphenyl is involved in CH-π interactions with the side chains of residues Lys9, Arg10, Asn13, Lys38, and Leu40 (Figure 2c). The methoxynaphthalene group T-stacks with Phe17 and Phe34. It also interacts with the side chains of Arg22, Val23, Thr26, and Ser30 through CH-π stacking. The aminoethyl tail at position R4 forms a salt bridge with Glu16. Other interactions between the ligand and the polar residues surrounding the binding pocket (Asn13, Asp20, and Lys38) are water-mediated H-Bonds (Figure 2b).

In contrast with previously solved crystal structures, the pyrazole-[4,3-c]pyridine scaffold of compound **1** is rotated by ∼90° (Figure 2e).^56,57^ The high-resolution crystal structure alone could not explain this experimental observation nor assemble any Quantitative Structure-Activity Relationships (QSAR). We set up different *in-silico* experiments with IPA to elucidate individual features of binding and rationalize differences towards previously published structures. In general applications, one should perform the curation of experimental structures to remove unphysical structural deformation.

### Base Validation

For the main manuscript, we focus on validation procedures that contribute to the understanding of the binding surface of PEX14. The base validation of the algorithm may be found in the supplementary material. This starts with ligand-only refinements and includes: i) parameter studies and parameter impact over molecular torsional angles and bond distances (S2 + S3); ii) impact of iteratively using IPA refinement over deformation energies (S4); iii) IPA refinement excluding protons from the RMSD bias potential (S5); iv) impact of IPA on atomic charges, motivating the use of IPA refined structures for MD-ligand parametrization; v) impact of the initial geometry over structural modification studies (S→O replacement); vi) impact of using different implicit solvation dielectric constants; vii) impact of structure resolution over IPA. We also perform an analysis with partial inclusion of the protein. Here our focus is to determine whether removing structural elements from the protein impacts the bound conformation of the protein-ligand complex. This is presented in S10. We see in our results that structural refinement without IPA introduces novel artifacts from using incomplete structural models, that lack biological relevance. Structural refinement of protein-ligand complexes including a few surrounding residues is consequently unadvisable. We stress once more, that though our validation is based on DFTB methods, the concept behind IPA is not bound to any underlying energy function. Technically, IPA may be combined with any quantum chemical method for which analytical gradients are also available (Hartree-Fock, Density Functional Theory, Coupled Cluster Theory, *etc*.), or even with force fields.

### Group-specific tension

Using the atom-selectivity of IPA’s constraining potential we restrained specific groups of compound **1** for group-specific tension evaluation (Figure 2d). Ring torsions are expected to bring minimal stress, as these rotations are controlled by linking methylene groups. This is corroborated by the calculations. Similar considerations apply to the rotation of the carboxamide. Rotation around the sulfur atom breaks the flow of π-electrons, hindering π-delocalization and leading to tension of 4.3 kcal/mol for the ligand to assume the binding conformation. Lastly, there is tension associated with the primary ammonium, which results from breaking an intramolecular H-Bond (dashed line in Figure 2d). As this interaction does not necessarily exist in solution, we consider it an artifact of using an implicit solvation model. This leads to an accumulated tension of 8.5 kcal/mol. Except for the sulfur atom, the protein distributes pressure over the ligand’s surface, which is consistent with the binding mode and interface. Further inclusion of the broken intramolecular H-Bond with the primary ammonium leads to a total stress of 14.5 kcal/mol, in good agreement with the total strain obtained using IPA on the whole structure (14.2 kcal/mol).

### The role of sulfur and simple SAR to relieve tension accumulated on sulfur

To understand which chemical modifications could improve the overaccumulation of tension on the sulfur atom, IPA was used to replace the sulfur with oxygen (compound **2**, *c.f*. Figure 2f and Table 1). The resulting strain is similar in magnitude, 14.4 kcal/mol. The difference concerning ligand **1** stems from the starting geometry: when generating the ether, we simply replaced the sulfur with an oxygen atom. Consequently, the optimization of **2** started from (experimental) CS bond distances, requiring 4 IPA iterations to obtain meaningful structures with the oxygen atom (see supporting material, S7).

Removing the sulfide altogether and introducing a proton instead (compound **3**) leads to a strain of 11.7 kcal/mol. This is larger than the expected tension by 1.6 kcal/mol, which results from the fact that we did not freely optimize the protons. Thus, also the proton replacement in compound **3** contains strain from the original experimental structure. When leaving all protons unconstrained, deviations between models for compound **3** drop below 0.2 kcal/mol. Interestingly, the binding of compounds **1**-**3** to the protein’s pocket is numerically identical (see Table 1). This clearly shows that the tension applied to the sulfur is compensated by interactions with the protein and that the electronic effect offered by this substituent is of relevance to binding. Further, calculated data shows slightly lower binding energies for compound **3** compared to compound **1**. The relative order of binding energies agrees with AlphaScreen EC_50_ data for compounds **1** (27.4 μM, see table S13.2) and **3** (17.5 μM).^57^ These calculations hinted that reducing the tension on this fragment of the ligand might require a change of hybridization for the atom replacing the sulfur.

The sulfur atom was consequently replaced by a methylene group (compound **7**). The total tension on the ligand lowers to 10.0 kcal/mol (reduction by 3.9 kcal/mol). Binding improves by 0.2 kcal/mol, which is far from the tension the protein applies on the sulfide group. This indicates that the sulfur atom is of relevance for electronic effects/interactions with the protein. Next, the methyl group in compound **7** is replaced by a fluorine atom (compound **15**, see Figure 4d). The total interaction energy lowers by 2.6 kcal/mol, potentially via halogen-π interactions. Using chlorine instead of fluorine (**16**), independently of the chemical stability of the respective ligand, impairs binding. This is likely caused by an increase in repulsion with the protein.

The sulfide was furthermore replaced by a chlorine atom (**9**, see also Figure 4e), which improved binding by 0.5 kcal/mol. Again, this is far from the total tension applied to the sulfur atom. The improvement in binding comes from lipophilic contacts with nearby residues, though the interactions are not as directed as in the case of compound **15**. We also investigated the bromine (**10**) and fluorine (**11**) variants of ligand **9**, but in both cases, binding was disfavored: in the case of bromine, the repulsion between protein and ligand should be the main issue, *i.e*., the lack of space in the binding pocket to accommodate the bromine atom; in the case of fluorine, its electron-withdrawing properties over the aromatic ring should weaken binding. The results of these calculations are corroborated by experimental EC_50_ data. For compound **14** we recorded an AlphaScreen EC_50_ of 80 μM against *Tb*PEX14 (see table S14.3); for the analog of compound **11** with a hydroxyl in position R_4_ the EC_50_ is 257 μM (the EC_50_s of this assay are typically one order of magnitude higher than K_i_).^57^ The structure shows that the sulfur’s main contact is Arg10’s alkyl chain. The sandwich stack with Asn13, together with the positive charge on Arg10 and Lys38 makes the electron-giving properties of sulfur to the aromatic group most likely the key aspect for binding. Supplementary Information S15 provides additional data.

In summary, the calculations revealed that the hybridization of sulfur is the main element leading to overaccumulation of tension. This tension may be relieved by effective change of hybridization, *e.g*., by replacing the sulfur with a methylene. Such changes will however impact the electron-giving properties of the aromatic group in position R1.

### Interactions in the crystal

Earlier in the manuscript several interactions were identified based on the classical geometric analysis of a crystal structure. Their relevance for binding could not be quantified, so the analysis remained purely qualitative. Here, IPA is used to analyze inhibitor-residue interaction energies including the role of water molecules (Figure 3).

**Figure 3.**
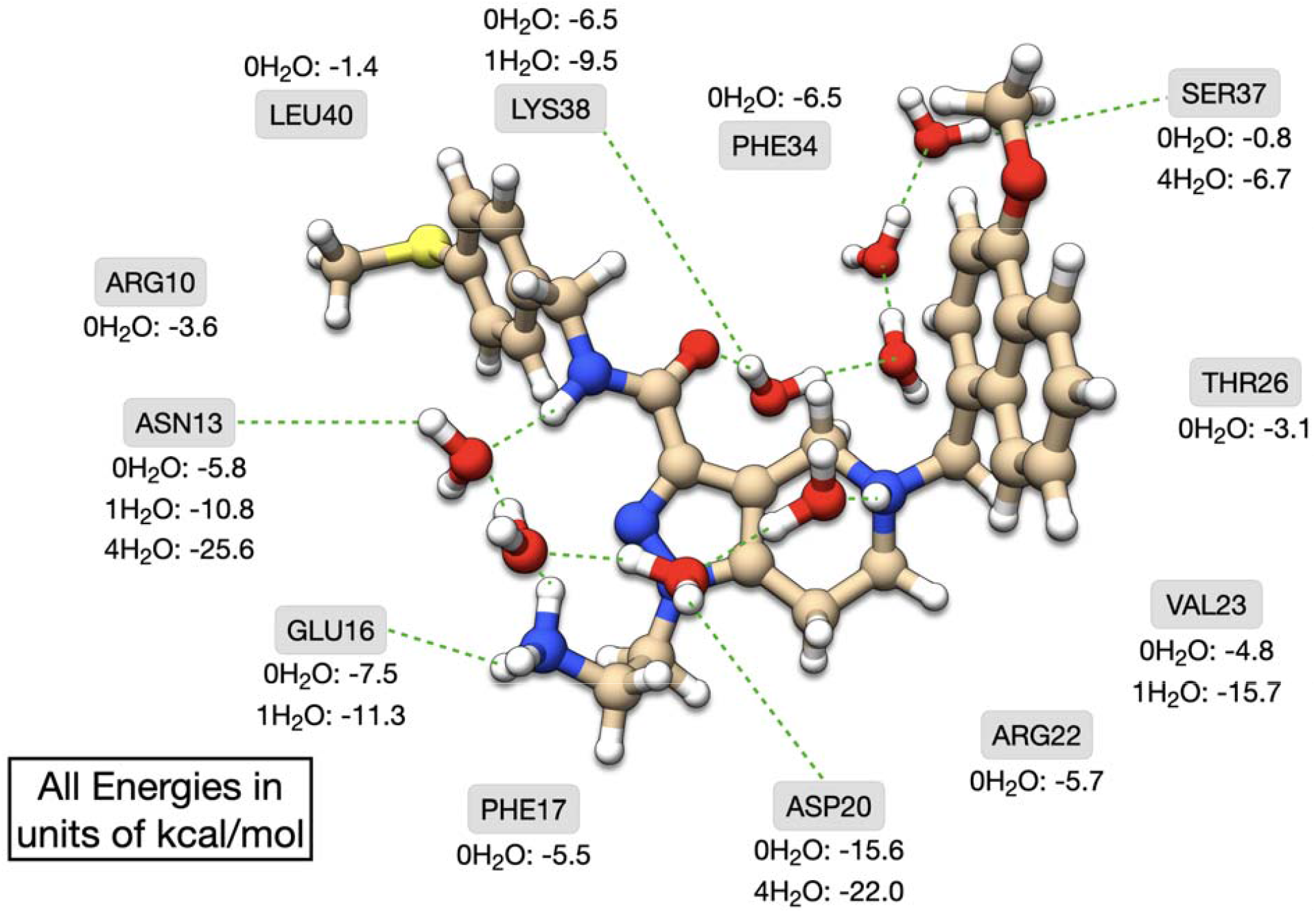
Quantification of specific ligand-residue interaction energies and the eventual role of water in those interactions. Green, dashed lines represent hydrogen bond networks. Different numbers of water molecules are used in the calculations according to their proximity to the interaction being studied. For instance, the trailing waters connecting the ligand to Ser37 are excluded when studying the interaction between the ligand and Asn13. In the case of Asn13, there is a calculation with one molecule of water, the closest to the residue. Details on the water exclusion concept are provided in the supplementary information (S10).

The crystal structure reveals two sets of water molecules in the complex’s envelope (Figure 2b): one, linking amide and tertiary ammonium in the ligand (intra); the other linking the ligand’s amide with Ser37 (inter). The interaction energies between ligand and those waters are, respectively, –22.0 and –6.7 kcal/mol. Interestingly, the second value matches the ligand-Ser37 interaction energy in the presence of explicit water, in agreement with the weak interaction recorded when no water is included (implicit solvation only). This indicates that the interaction with Ser37 is purely water-mediated, and this residue plays a stabilizing role in holding the water network together. Other inhibitor-residue interactions investigated show that the inclusion of water favors binding, even in the case of the lipophilic Val23, where only the sidechain comes close to the ligand. This shows that the role of the water envelope goes beyond bridging H-Bonds. Our calculations reveal that the trail of waters connecting ammonium and amide in the ligand donates ∼0.12 electrons to the ligand, though the total change in electronic populations amounts to ∼0.42. Including the whole protein system (and waters) shows a net transfer of 0.17 electrons to the ligand, while the total change in ligand electronic populations is 0.84 electrons. But if only the protein is considered (no water), then the total amount of electrons transferred zeroes. We, therefore, conclude that water promotes charge transfer and electronic exchange between the main interaction partners: by supplying the ligand electronically, which in turn supplies the protein too, the water envelope acts as a vector that promotes the covalency of interactions between protein and ligand, though no true covalent bond is established.

Additional calculations we performed reveal that the aromatic rings on the ligand establish strong stacking interactions, not only with Phe17 and Phe34 but also with nearby arginines. The contribution of the stacking interaction with Arg22 is in the range of the phenylalanines already mentioned. The interaction with Arg10 is also of significance, which justifies the requirement of PEX14 for electron-rich aromatic rings in the sub-pockets previously observed.^57^

The present analysis provides quantitative indications of the role of water and particular residues for binding. The analysis is further extended to all residues composing the pocket (Supplementary Information S11) and it shows how to analyze interactions between ligand fragments and specific residues. An energy-based evaluation of single-site ligand-residue interactions provides key additional information that may be used to assemble a quantitative binding model. To make this available to the SBDD community, a Python script was released in the ULYSSES repository. The script performs ligand-deformation and protein fragmentation analyses. A benchmark of the method used for energy evaluation against higher-level Density Functional Theory methods is available in (S11), showing equivalent quality.

### The role of the primary ammonium

It was previously verified that the primary ammonium in position R_4_ is key for high affinity.^56,57^ The affinity data on compounds **1** and **14** further corroborate these observations. The crystal structure of compound **1** shows Glu16 in the vicinity of the ligand with N^+^---O^−^ distances of 2.89 and 5.50 Å. In our attempt to rationalize the role of this group, we ran a calculation with the respective amine. Alkalinization hindered binding by 0.3 kcal/mol. The ammonium/amine was then replaced with a hydroxyl group (**14**). Surprisingly, binding was favored by 0.7 kcal/mol compared to **1**, in clear contradiction with the experimental data (tables S14.2 and S14.3: the EC50 of **14** is larger than that of **1** by a factor of 3).^56,57^ To ensure the result was not an artifact caused by the structural deformation, we let the hydroxyl group free during IPA. The oxygen used the additional freedom to move apart from the glutamate’s carboxylate (5.48 Å → 6.86 Å), and binding improved by almost 2.5 kcal/mol. This implies that the improvement in binding observed is driven by the minimization of repulsion and not by the strengthening of attractive forces.

Next, an explicit water molecule from the crystal structure was added to the calculations (Figures 4a,b). The protons were unconstrained during the optimization. The binding energies of ligands **1** and **14** improved, in the former case to –26.5 kcal/mol, and in the latter to –24.3 kcal/mol. This difference of 2.2 kcal/mol is remarkably close to the 1.6-1.8 kcal/mol obtained experimentally,^56^ despite the structural simplicity of the model used. A closer look at the optimized geometries reveals that the ammonium group acts as an H-Bond donor to the key structural water molecule. Inspection of the structure of **14** with water indicates that the hydroxyl group tries to act as an H-Bond donor to the carboxylate and as an acceptor to the key structural water. We thus conservatively conclude from our calculations that this structural water, conserved in most complexes characterized thus far by X-ray crystallography, favors pyrazole substituents with strong H-Bond donor character.

### A change in the binding pose

The novelty of ligand **1** is its binding pose compared to earlier determined poses of compounds built around the same scaffold (Figure 2e),^56,57^ which was puzzling to our understanding of the protein-ligand interactions. Our initial guess was that the pyramidal inversion of the tertiary amine would increase repulsion with the protein, thus disfavoring binding. To great surprise, binding improves by 0.8 kcal/mol. Detailed inspection of the structures reveals that several alkyl chains favor the amine inversion.

Further examination of the crystal structure reveals that in both protein-ligand complexes in the asymmetric unit, there is a molecule of water between the protein and the tertiary amine. In one of the cases, there is a trail of solvent molecules bridging between the two amine groups in the ligand (Figure 4c). This water envelope, exclusive to this complex, forms strong interactions with the polar residues on the protein side (Arg and Asp), accounting for the new binding pose. The inversion of the tertiary amine *i)* breaks an H-Bond with the entrapped water molecule, *ii)* places a ligand’s methylene group close to that same molecule of water and *iii)* increases repulsion.

#### Semi-Empirical Quantum Chemical Docking

IPA was used to investigate swapping the orientation of the amide group in compound **1**, *i.e*., compound **1**f (Figure 2g). EC_50_s were collected for **1** and **1**f, and this data is provided in supplemental (S13). Despite the simplicity of the modification, EC_50_s are larger for **1**f by a factor of 2.7 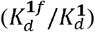. IPA was initially used to generate a binding pose for **1**f that kept the H-Bond donor-acceptor orientation of **1**. This led to unrealistic binding energies since it either distorted the amide or disrupted the π-stacking of aromatic rings. We then investigated whether the inversion of the donor-acceptor atoms would be more consistent. Taking advantage of the capabilities of the new algorithm, we docked compounds **1** and **1**f to the protein and the resulting structures showed relative dissociation constants of 3.1 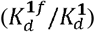. In this case, the water envelope was treated implicitly. Consequently, we see the agreement between calculated and experimental values rather conservatively and focus on the relative order of binding. Still, we can see that IPA-based docking can provide useful results. This can be, for example, used for post-screening scoring of *in silico* hits from high-throughput software. Supplemental S16 contains another case study.

### Thermodynamics of Binding

To further evaluate the impact of solvation waters on binding, several models were assembled for compounds **1** and **14**. For compound **1** we tested starting directly from the crystal structure (setD), as well as from in-pocket (setIPA) and unconstrained optimization (setOPT). For compound **14** only the last two variants were tested. Details on the affinity calculation protocol are provided in the SI (S17). The calculations with the full protein and all waters detected in the crystal structure reinforce the conclusion of water as a key element for determining affinity (Figure 5a,b) since the binding energies are always lower. The relative affinities show similar trends, though the effect is less pronounced because entropies of binding are negative. Reducing the protein part used in the calculations impacts energies and affinities (Figures 5a,b). Both using setD and setIPA there is a tendency for the calculations to exaggerate the role of water for binding. This evidences the importance of missing residues in determining the strength of protein-ligand-water interactions. In setOPT differences reduce beyond what is expected given the full construct, and this applies to both pocket-cuts studied. Interestingly, the complexes in setOPT indicate in some cases that water is not beneficial for binding, corroborating the role of distal residues for a balanced description of interactions. This shows that limiting the pocket too extensively causes artifacts and disagrees with the calculations using the full protein construct and previous data.^56,57^ It also exemplifies the importance of the entire water network for precise structure analysis. Entropy corrections from the two pocket cuts show, interestingly, a similar behavior, which is visible in the results in Figure 5b. Using the smaller pocket leads to overly large entropy penalties, which render results counterintuitive even for the full protein construct.

**Figure 4.**
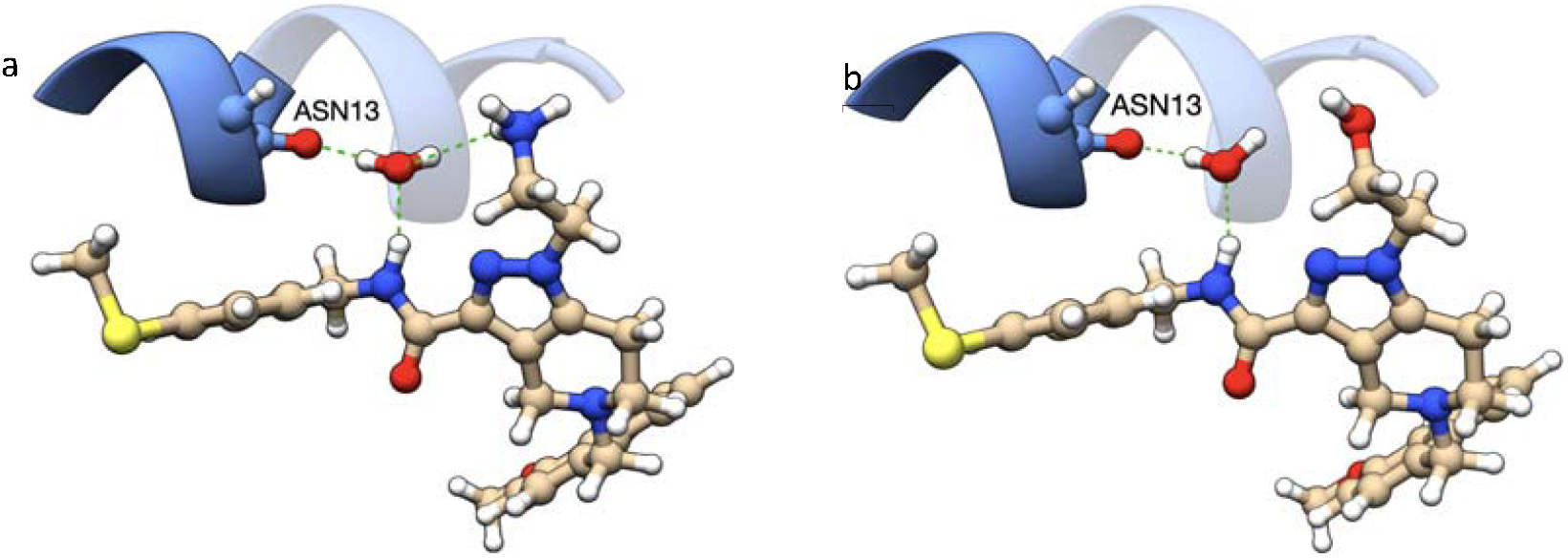
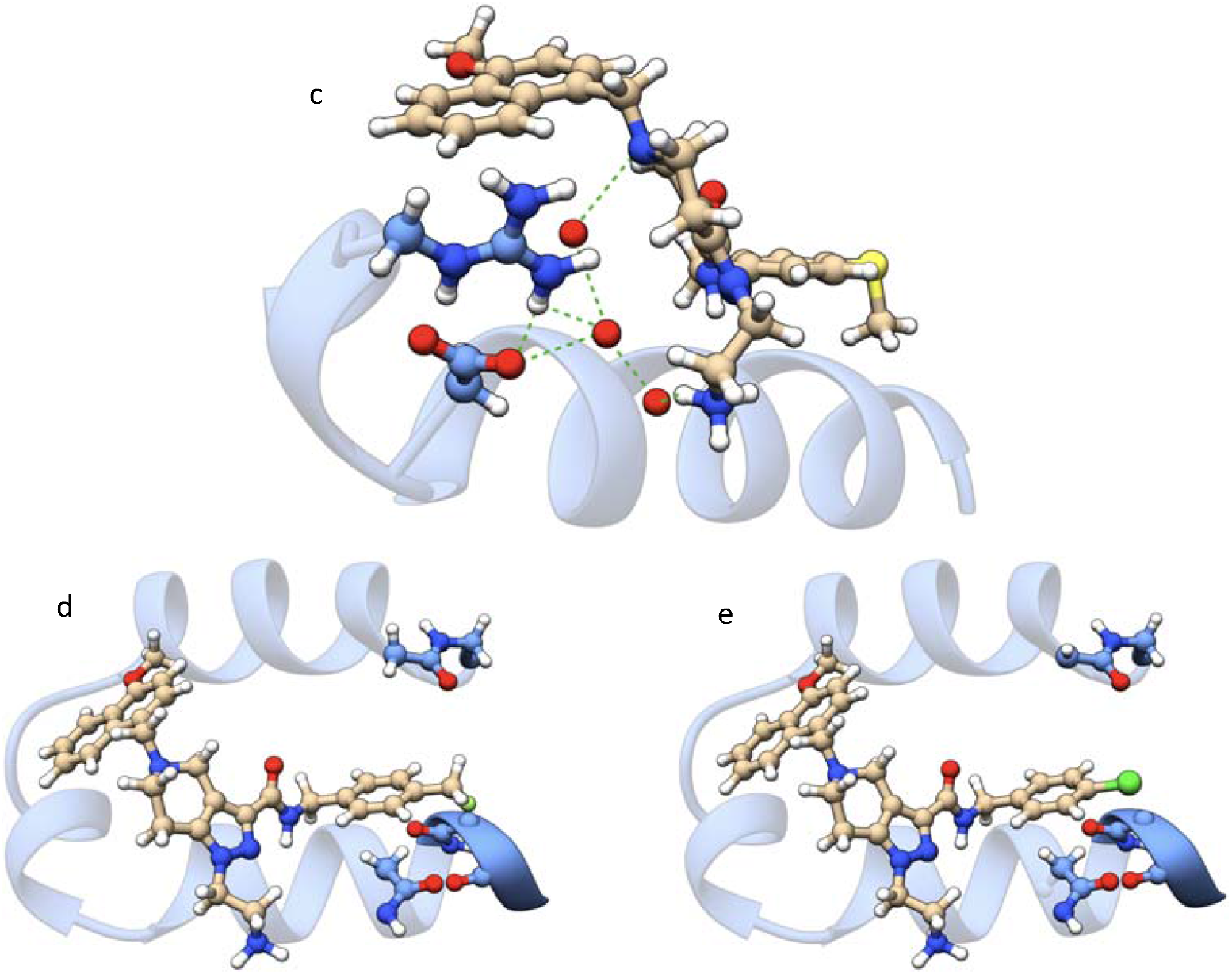
In-pocket optimized structure of a) 1 and b) 14 with a key molecule of water; c) the trail of water molecules in the crystal structure; in-pocket optimized binding poses for d) 15, e) 9. In all subfigures, the protein’s backbone scaffold is shown as blue ribbons. H-Bond networks are marked with green dashed lines.

**Figure 5.**
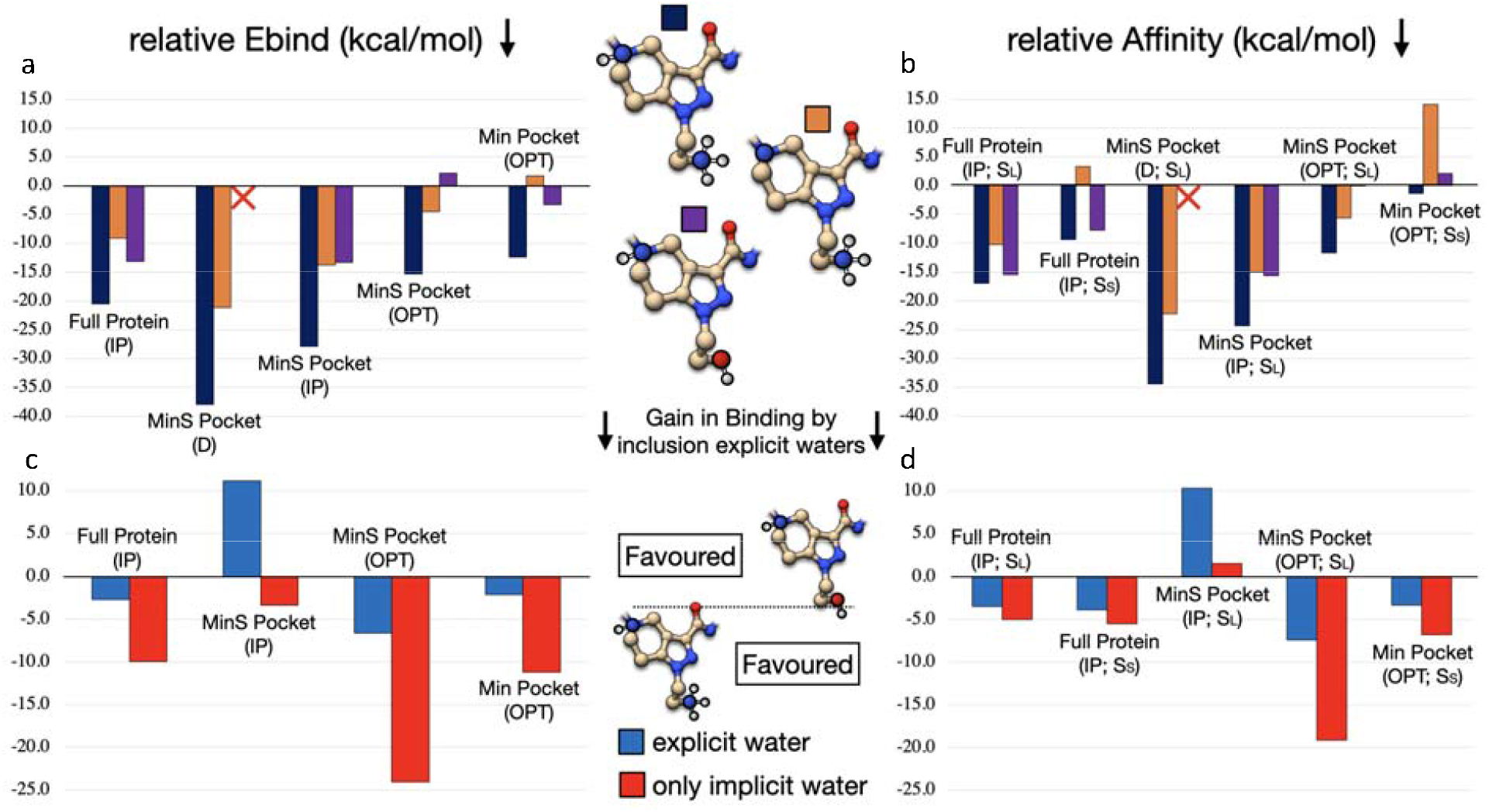
a) Influence of solvation waters onto relative binding energies for ligand 1 in two protonation states and ligand 14. b) Influence of solvation waters onto relative affinities, considering in this case the entropic and enthalpic corrections, including conformational entropies of the ligand only. **c)** R**elative binding energies for ligands 1 and 14. d) Relative affinities of ligands 1 and 14, using similar calculation methods as in case a)**. In the figure, IP stands for in-pocket optimization, OPT for unconstrained optimization and D is the direct calculation, using the crystal structure with protons added by Chimera. Note that direct calculations are not possible for ligand **14**. The MinS pocket consists of the first row of amino acids interacting with the ligand with all their side chains. The smaller pocket Min retains only those side chains that are directed to the ligand (8 Å). Enthalpy and entropy corrections were calculated from both pocket constructs after full optimization. This is marked with S_L_ or S_S_ in the plots, corresponding to either MinS or Min pockets.

We also performed calculations on the relative affinity of compounds **1** and **14** (Figures 5c,d). Using the complete protein with IPA leads to meaningful results matching the experimental data. The entropy corrections to the relative affinities are minimal. However, using a pocket instead of the full protein leads to strong discrepancies in the results, which invalidates any conclusion from those reduced models. Curiously, we observe that entropy corrections are in both pocket cases qualitatively suitable. This appears contradictory with observations from the previous paragraph, however, unlike the previous data, we compare systems with equivalent hydration states. Contrary to enthalpy corrections and entropies, data for ligands **1** and **14** shows that using small pocket constructs will systematically lead to poor results, mainly due to poor energetics, no matter which optimization algorithm is employed. Hence, these findings suggest that substantial constructs are essential for a realistic assessment of binding energies in biological systems; otherwise, energy calculation becomes a limiting factor in evaluating binding affinities. This is particularly interesting in the case we study since the ligands analyzed target a highly lipophilic pocket of PEX14, where the impact of electrostatics is expected to be minimal. SI sections S17 and S18 contain additional test cases and additional entropy considerations.

## DISCUSSION

The present manuscript shows IPA as an extremely versatile multi-purpose tool for investigating the mechanisms underlying experimental observations: corroborating hypotheses, and revealing information encoded in experimental structural models. In the analysis we performed, we show how chemistry in general, and quantum chemistry in particular, provide a detailed description of complexes between PEX14 and a series of inhibitors, defining a quantitative binding model. The role of positively charged residues and their targeting of electron-rich aromatic rings was explained.^56,57^ The analysis, not restricted to aromatic groups of the ligand, was started with structurally refined crystallographic data.

IPA allows selective optimization of atoms and functional groups composing a structure. Unlike previous methods, where the protein backbone is held fixed from the crystal structure (*cf*. reference 60), all bonds and angles might be refined if they are found violating chemical principles, while torsions are effectively retained. We believe that this feature will allow IPA to be crucial in assembling reference data from biological databases like PDBbind,^61^ since *i)* it avoids problems of fixing atoms during geometry optimization or using too short subconstructs,^62,63^ but also *ii)* it helps keeping the structures as close as possibly to experimental data.^64^ Only with the use of this multifaceted tool we were able to establish a series of *in silico* experiments that allowed us to investigate the nature of interactions for a PPI-inhibitor system quantitatively. A quantum mechanical role was established for different fragments of the ligand, but also for the solvent. The latter, according to our calculations, is crucial for promoting protein-ligand contacts since it enhances the electron-giving properties of the ligand and acts as an electronic mediator between inhibitor and target.

IPA allowed us to construct minimal computational models that could reproduce experimental observations quantitatively. These allowed us to pose new questions related to the refinement of binding affinity, which matched closely prospective experimental data. It showed limitations in some of the hypotheses used in building oversimplified thermodynamic models of binding. Altogether, this will open the door to more rational approaches in drug discovery and in generating computational models to explain and rationalize non-bonded complexes.

## CONCLUSION

We introduce a biased optimization technique, in-pocket analysis, that is useful in curating experimental structures, for understanding protein-induced tension, explaining SAR data, docking, and understanding the role of molecules or functional groups for binding. The bias potential introduced by in-pocket affects primarily bond distances, then bond angles, and lastly (and minimally) the torsions. This is key for reducing unphysical tension in bonds and angles while retaining molecular conformation. The extent to which these quantities are affected is controlled with two parameters, *κ* and *α*.

The method is used extensively in the analysis of crystal structures solved in our groups and in explaining SAR data around the biological target, allowing the extraction of the maximal amount of quantifiable information. This permits understanding the role of functional groups in the protein-ligand complex, building a more realistic QSAR model, and understanding which structural modifications of the original ligand could improve binding, as exemplified in this manuscript.

## METHODS

### Implementation and computational details

Ultimately, the problem we are interested in solving consists of a geometry optimization of the substructure of a protein-ligand complex under the constraint of the experimental data. Defining these constraints as penalty functions permits solving the problem using unconstrained optimization techniques.^65^ For efficiency, the quaternion method of Coutsias and coworkers is used to evaluate gradients,^66^ whereas the Hessian of the bias potential is evaluated numerically.

An easy-to-use interface for IPA is available in our in-house code *ULYSSES*.^58^ Besides experimental/reference systems, the only input parameters required are κ and α. Users are encouraged to directly use the values determined in this work.

All quantum chemical calculations performed here used exclusively GFN2-xTB.^67^ Other semi-empirical or QM methods could equally be employed. Solvation was modelled with ALPB.^68^ UCSF Chimera and ChimeraX, as developed by the Resource for Biocomputing, Visualization, and Informatics at the University of California, San Francisco, with support from NIH P41-GM103311 was used to generate the visualizations of molecular models.^69-72^

### Protein expression and purification

#### Trypanosoma brucei

N-terminal domain PEX14 (*Tb*PEX14; 19-84) was cloned into a pETM11 (EMBL). The expression was carried out in *E. coli* BL21 (DE3) using an autoinduction medium.^73^ *Tb*PEX14 was purified from the crude lysate using Immobilized Metal Affinity Chromatography (IMAC). The eluted protein was incubated with TEV protease overnight at room temperature to cleave the His-tag. Protease and the cleaved His-tag were removed by reverse IMAC. The protein contained in the flow-through was further purified by size exclusion chromatography (SEC) using crystallization buffer (10 mM Tris pH 8.0, 100 mM NaCl, 5 mM β-mercaptoethanol).

### Crystallization and X-ray structure solution

Purified *Tb*PEX14 was mixed with a 10-fold molar excess of compound **1** (50 mM solution in DMSO) and the mixture was incubated for 1h at room temperature. The excess of ligand and DMSO were removed by washing the complex with the storage buffer at 4 °C on a 10 kDa-cutoff Centricon concentrator. Before crystallization, the complex was concentrated to about 30 mg/ml. The initial crystallization trials were set up using commercial kits in the TTP LabTech Mosquito automated crystallization workstation. The crystals suitable for testing were transferred into a cryo-protectant solution containing the harvesting solution supplemented with 25% (v/v) ethylene glycol and cryo-cooled in liquid nitrogen. Diffraction data were collected at the Swiss Light Source at Paul Scherrer Institut (SLS, Villigen, Swiss) beamline PXIII. The best diffracting crystal grew in 0.2M sodium malonate dibasic monohydrate and 20% (w/v) PEG 3350 at 4°C for 17 months. The experimental data were processed using XDS software,^74^ before further processing through STARANISO^75^ for anisotropy correction to give a 1.1 Å dataset. The crystal belonged to the space group C 1 2 1. The Matthews coefficient analysis suggested the presence of two molecules in the asymmetric unit.^76^ The structure was solved by molecular replacement using Phaser^77^ with *Tb*PEX14 structure (PDB code: 6SPT)^57^ as a search model. The analysis of the electron density calculated with (Fo – Fc) and (2Fo – Fc) coefficients allowed us to build the initial model and unambiguously place the ligand using COOT.^78^ The starting model was refined by iterations of manual and automated refinement cycles using Refmac5^79^ at R/Rfree 12.6/16.2%. Throughout the refinement, 5% of the reflections were used for the cross-validation analysis, and the behavior of R_free_ was employed to monitor the refinement strategy.^80^ Statistics of data collection and refinement are reported in the supplementary material (S13).

### Synthesis of compounds 1, 1f, and 14

Compounds **1** and **14** were prepared according to the protocols previously defined for similar ligands.^56,57^ Target compound **1f** on the other hand required the development of a completely new synthesis route, which is displayed in Scheme 1. The detailed synthesis procedures and the spectroscopic data are reported in the following paragraphs. Recorded NMR spectra of target compound **1f** and its precursors **17-21** can be found in the supplementary material (S20).

#### General methods

Air and water-sensitive reactions were performed in flame-dried glassware under an argon atmosphere. Solvents used for column chromatography, extractions, and recrystallization were purchased in technical grade and were distilled before use. Solvents used for the reversed-phase chromatography and HPLC-MS analyses were purchased from Thermo Fisher Scientific in HPLC-quality. Reagents and dry solvents were purchased from Sigma Aldrich (Merck), ABCR, Alfa Aesar, Thermo Fisher Scientific, TCI, and Carl Roth and were used without further purification. Analytical thin layer chromatography (TLC) was performed on silica-coated plates (silica gel 60 F_254_) purchased from VWR. Compounds were detected by ultraviolet (UV) irradiation at 254 or 366 nm. Manual flash column chromatography was performed using silica gel 60 (particle size: 0.040-0.063 mm) available from VWR. Automated preparative chromatography was performed on a Grace Reveleris Prep purification system using linear gradient elution and Büchi Reveleris Silica 40 μm cartridges for normal-phase and Büchi Reveleris C18 40 μm cartridges for reverse-phase separations. ^1^H, ^19^F and ^13^C NMR spectra were recorded at room temperature on a Bruker AV-HD400 spectrometer operating at 400 MHz. The NMR peaks are reported as follows: chemical shift (*δ*) in parts per million (ppm) relative to residual non-deuterated solvent as internal standard (CHCl_3_: *δ* (^1^H) = 7.26 ppm, *δ* (^13^C) = 77.2 ppm; DMSO: *δ* (^1^H) = 2.50 ppm, *δ* (^13^C) = 39.5 ppm; CH_3_OH: *δ* (^1^H) = 3.31 ppm, *δ* (^13^C) = 49.0 ppm), multiplicity (s = singlet, d = doublet, t = triplet, q = quartet, m = multiplet and br s = broad signal), coupling constant (Hz) and integration. HPLC-UV/MS analyses were performed on a Dionex UltiMate 3000 HPLC system coupled with a Thermo Scientific™ ISQ™ EC Single Quadrupole Mass Spectrometer, using the following method: Thermo Scientific™ Accucore™ RP-MS LC-column (2.1 x 50 mm, 2.6 μm); gradient: 5 to 95% of acetonitrile + 0.1% formic acid v/v in water + 0.1% formic acid v/v over 5 min period; flow rate: 0.6 mL/min; UV detection at 254 nm. All target compounds exhibited a purity greater than 95%, which was determined by HPLC-UV/MS analyses.

### Synthetic procedures and spectroscopic data for flipped amide 1f

#### Ethyl 3-((*tert*-butoxycarbonyl)(2-cyanoethyl)amino)propanoate (17)

A solution of β-alanine ethyl ester (32.00 g, 208.3 mmol, 1.0 eq) and NaOH (8.32 g, 208.3 mmol, 1.0 eq) in CH_3_OH (200.0 mL) was stirred at rt for 30 min. After the addition of acrylonitrile (16.32 mL, 15.28 g, 249.9 mmol, 1.2 eq) at rt, the reaction mixture was heated until reflux for 4 h. The reaction mixture was then cooled to 0 °C and Boc anhydride (45.48 g, 208.3 mmol, 1.0 eq) was added. The resulting reaction mixture was warmed to RT and stirred overnight. The next day the precipitate was filtered off, washed with methanol, and the filtrate was concentrated under reduced pressure. The formed solid was filtered off again and washed with EtOAc. The filtrate was taken up in EtOAc (150 mL) and H_2_O (150 mL). The two layers were separated, and the aqueous phase was extracted with EtOAc (3 x 100 mL). Afterward, the combined organic layers were washed with brine, dried over Na_2_SO_4_, filtered, and concentrated under reduced pressure to give the desired product **17** as a colorless oil (56.11 g, 207.6 mmol, 100% over 2 steps). *R*_f_ = 0.60 (pure EtOAc, KMnO_4_ stain). ^1^H NMR (400 MHz, CDCl_3_): *δ* = 4.09 (q, *J* = 7.2 Hz, 2H), 3.55 – 3.45 (m, 4H), 2.62 – 2.49 (m, 4H), 1.42 (s, 9H), 1.21 (t, *J* = 7.2 Hz, 3H) ppm. ^13^C NMR (101 MHz, CDCl_3_, two sets of rotamers): *δ* = 172.0, 171.6, 154.9, 154.5, 146.7, 118.2, 117.8, 85.1, 80.8, 60.7, 51.7, 44.7, 44.1, 34.2, 33.6, 28.3, 27.4, 17.5, 16.9, 14.1 ppm.

**Scheme 1.**
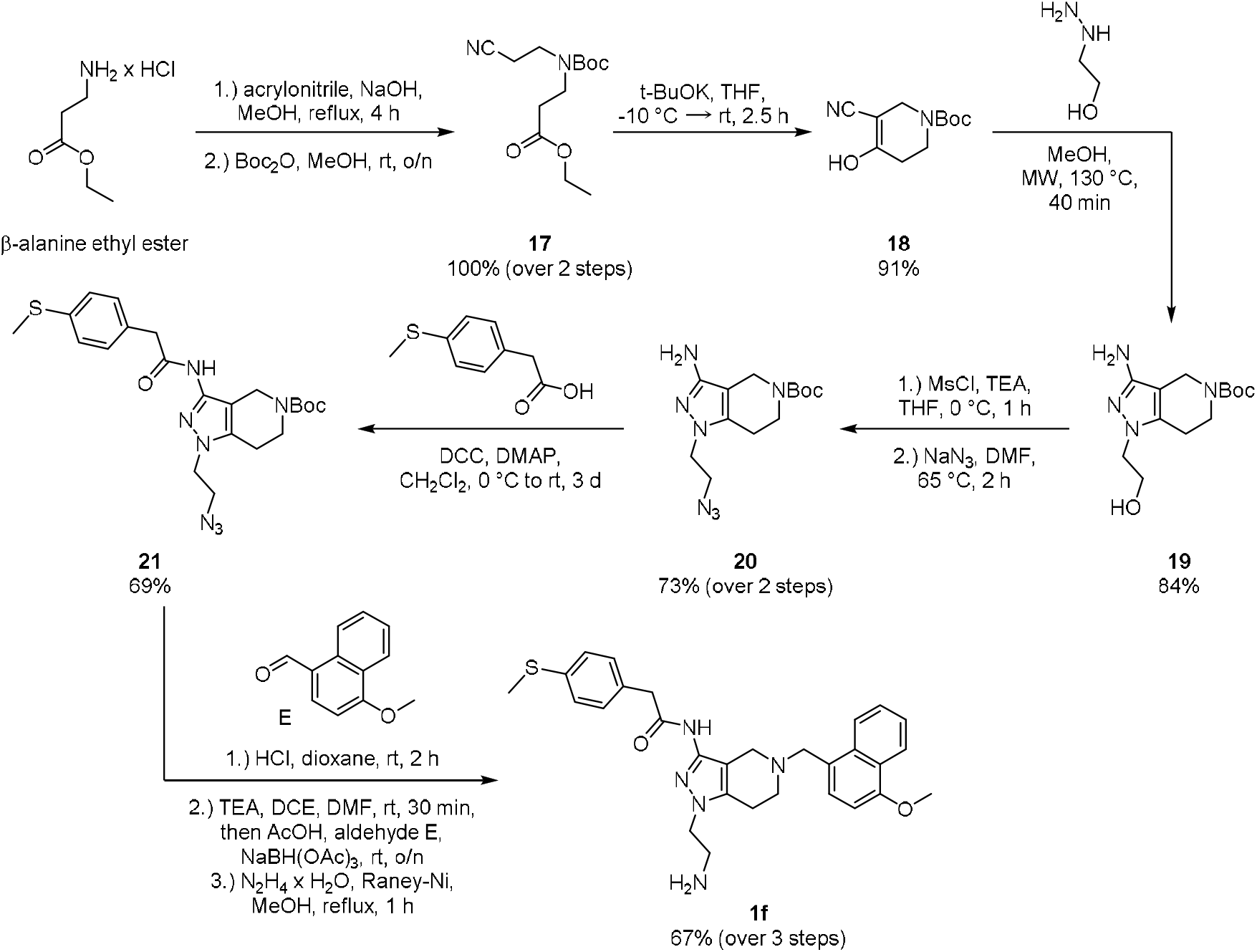
Synthetic route for the preparation of the flipped amide **1f**

#### *tert*-Butyl 5-cyano-4-hydroxy-3,6-dihydropyridine-1(2*H*)-carboxylate (18)

Nitrile **17** (55.96 g, 207.0 mmol, 1.0 eq) was dissolved in dry THF (300.0 mL) and cooled to −10 °C. Afterward, *t*-BuOK (27.88 g, 248.4 mmol, 1.2 eq) was added at −10 °C and stirred for 30 min. The reaction mixture was allowed to reach rt and stirred for an additional 2.5 h. After this period, water (100 mL) was added and the pH was adjusted to 3-5 with 2N hydrochloric acid. The aqueous phase was extracted with EtOAc (3 x 100 mL) and the combined organic layers were washed with brine (100 mL), dried over Na_2_SO_4_, filtered, and concentrated under reduced pressure. The crude product was purified *via* gradient flash column chromatography (c-Hx/EtOAc: 4:1, 2:1, 1:1, pure EtOAc) and reprecipitation from EtOAc/Et_2_O/c-Hx to obtain piperidinone **18** as a white solid (42.34 g, 188.8 mmol, 91%). *R*_f_ = 0.38 (hexane/EtOAc 1:1, UV). ^1^H NMR (400 MHz, DMSO-d6): *δ* = 10.86 (br s, 1H), 3.89 (s, 2H), 3.46 (t, *J* = 5.6 Hz, 2H), 2.28 (t, *J* = 5.2 Hz, 2H), 1.41 (s, 3H) ppm. ^13^C NMR (101 MHz, DMSO-d6): *δ* = 167.0, 153.92, 117.8, 80.0, 78.1, 41.8, 28.7, 28.4 ppm.

#### *tert*-Butyl 3-amino-1-(2-hydroxyethyl)-1,4,6,7-tetrahydro-5*H*-pyrazolo[4,3-*c*]pyridine-5-carboxylate (19)

To a solution of piperidinone **18** (2.000 g, 8.918 mmol, 1.0 eq) in dry CH_3_OH (16.0 mL) hydroxyl ethyl hydrazine (667 μL, 747 mg, 9.810 mmol, 1.1 eq) was added at rt. The resulting reaction mixture was heated for 40 min in a microwave reactor at 130 °C. Afterward, the solvent was removed under reduced pressure and the crude product was purified *via* gradient flash column chromatography (CH_2_Cl_2_/CH_3_OH: 1, 2, 3, 4, 5, 7, 10% CH_3_OH) to obtain pyrazole alcohol **19** as a white solid (2.453 g, 8.687 mmol, 97%). *R*_f_ = 0.29 (CH_2_Cl_2_/CH_3_OH 9:1, UV). ^1^H NMR (400 MHz, CDCl_3_): *δ* = 4.18 (s, 2H), 4.00 (br s, 1H), 3.95 (t, *J* = 4.8 Hz, 2H), 3.79 (t, *J* = 4.8 Hz, 2H), 3.60 – 3.50 (m, 2H), 2.52 (t, *J* = 6.00 Hz, 2H), 1.40 (s, 9H) ppm. ^13^C NMR (101 MHz, CDCl_3_): *δ* = 155.2, 145.6, 141.1, 97.1, 80.1, 62.1, 49.8, 42.5, 39.4, 28.5, 23.9 ppm. HPLC-MS (ESI): *m/z* = 283 [M+H]^+^; retention time: 2.52 min.

#### *tert*-Butyl 3-amino-1-(2-azidoethyl)-1,4,6,7-tetrahydro-5*H*-pyrazolo[4,3-*c*]pyridine-5-carboxylate (20)

Pyrazole **19** (10.57 g, 37.44 mmol, 1.0 eq) was dissolved in dry THF (75.0 mL) under an argon atmosphere and cooled to 0 °C. After addition of triethylamine (10.38 mL, 7.575 g, 74.88 mmol, 2.0 eq) and methanesulfonyl chloride (3.767 mL, 5.574 g, 48.67 mmol, 1.3 eq), the resulting reaction mixture was stirred for 1 h at 0 °C. Afterward, the reaction mixture was filtered through celite, and the solvent was removed under reduced pressure. The crude mesylate was redissolved in dry DMF (45.0 mL) and NaN_3_ (7.302 g, 112.3 mmol, 3.0 eq) was added. The final reaction mixture was then heated to 70 °C and stirred for 2 h. After this period, the reaction was quenched by the addition of sat. NaHCO_3_ (100 mL). The aqueous phase was then extracted with EtOAc (3 x 100 mL) and the combined organic layers were washed with water (50 mL) and brine (50 mL), dried over Na_2_SO_4_, filtered, and concentrated under reduced pressure. The crude product was purified *via* gradient flash column chromatography (c-Hx/EtOAc: 50, 75, 100% EtOAc; EtOAc/CH_3_OH: 2, 4, 6, 10% CH_3_OH) to obtain pyrazole azide **20** as an off-white solid (8.442 g, 27.47 mmol, 73%). *R*_f_ = 0.33 (EtOAc/CH_3_OH 9.8:0.2, UV). ^1^H NMR (400 MHz, CDCl_3_): *δ* = 4.28 (s, 2H), 4.06 (t, *J* = 5.2 Hz, 2H), 3.73 (t, *J* = 5.6 Hz, 2H), 3.69 – 3.49 (m, 4H), 2.64 (t, *J* = 5.6 Hz, 2H), 1.46 (s, 9H) ppm. ^13^C NMR (101 MHz, CDCl_3_): *δ* = 155.2, 146.5, 140.4, 98.5, 80.1, 51.8, 46.6, 42.4, 39.3, 28.6, 24.0 ppm. HPLC-MS (ESI): *m/z* = 308 [M+H]^+^; retention time: 2.88 min.

#### *tert*-butyl 1-(2-azidoethyl)-3-(2-(4-(methylthio)phenyl)acetamido)-1,4,6,7-tetrahydro-5*H*-pyrazolo[4,3-*c*]pyridine-5-carboxylate (21)

A solution of 4-(Methylthio)phenylacetic acid (385 mg, 2.12 mmol, 1.0 eq) and DMAP (77.5 mg, 0.635 mmol, 30 mol%) in dry CH_2_Cl_2_ (25.0 mL) was cooled to 0 °C. After the addition of DCC (480 mg, 2.33 mmol, 1.1 eq) and pyrazole azide **20** (650 mg, 2.12 mmol, 1.0 eq) under argon, the reaction mixture was slowly warmed to room temperature and stirred for 1 d. After this period, the precipitated dicyclohexylurea was removed by filtration, and the solvent was removed under reduced pressure. The residue was purified by gradient flash column chromatography (c-Hx/EtOAc: 20, 35, 50, 70, 90, 100% EtOAc) and thereby the desired pyrazole amide **21** was obtained as a white foam (687 mg, 1.46 mmol, 69%). *R*_f_ = 0.35 (pure EtOAc, UV). ^1^H NMR (400 MHz, CDCl_3_): *δ* = 7.77 (s, 1H), 7.30 – 7.23 (m, 4H, coinciding with solvent signal), 4.32 (s, 2H), 3.92 (s, 2H), 3.76 – 3.60 (m, 6H), 2.72 – 2.62 (m, 2H), 2.48 (s, 3H), 1.47 (s, 9H) ppm. ^13^C NMR (101 MHz, CDCl_3_): *δ* = 169.4, 155.1, 138.9, 131.1, 130.5, 130.1, 127.2, 80.1, 51.8, 47.5, 43.3, 42.2, 40.2, 28.6, 24.0, 15.7 ppm. HPLC-MS (ESI): *m/z* = 472 [M+H]^+^, 494 [M+Na]^+^; retention time: 3.70 min.

#### *N*-(1-(2-aminoethyl)-5-((4-methoxynaphthalen-1-yl)methyl)-4,5,6,7-tetrahydro-1*H*-pyrazolo[4,3-*c*]pyridin-3-yl)-2-(4-(methylthio)phenyl)acetamide (1f)

Boc-deprotection: Pyrazole azide precursor **21** (163 mg, 0.459 mmol, 1.0 eq) was dissolved in dry dioxane (4.0 mL) under an argon atmosphere. After the addition of HCl (4.0 M in dioxane, 1.21 mL, 6.43 mmol, 14.0 eq), the resulting reaction mixture was stirred for 2 h at rt. Afterward, the solvent was removed under reduced pressure and the off-white residual solid was dried in vacuo. Reductive amination: The deprotected crude material was suspended in a mixture of dry DCE and DMF (8.0/2.0 mL) under argon and TEA (63.6 μL, 46.4 mg, 0.459 mmol, 1.0 eq) was added. After stirring the reaction for 30 min at rt, 4-Methoxy-1-naphthaldehyde (128 mg, 0.689 mmol, 1.5 eq), AcOH (26.3 μL, 27.6 mg, 0.459 mmol, 1.0 eq) as well as NaBH(OAc)_3_ (195 mg, 0.918 mmol, 2.0 eq) were added and the resulting mixture was stirred overnight. The following day the reaction was quenched with a saturated aqueous solution of NaHCO_3_ (25 mL) and the aqueous phase was extracted with EtOAc (3 x 35 mL). Afterward, the combined organic layers were washed with brine (25 mL), dried over Na_2_SO_4_, filtered, and concentrated under reduced pressure. The obtained crude secondary amine was used for the next step without further purification. Azide reduction: An aqueous slurry suspension of Raney-nickel (1 x tip of a spatula) was washed under argon with MeOH (3 x 5.0 mL). Afterward, a solution of the crude azide in MeOH (9.0 mL) was added under argon. After the addition of 50% hydrazine monohydrate solution (143 μL, 147 mg, 1.47 mmol, 3.2 eq), the reaction mixture was heated until reflux and stirred for 1 h. Afterward, Raney-nickel was filtered off *via* celite, washed with MeOH (25 mL), and the solvent was removed under reduced pressure. The residue was purified by gradient flash column chromatography (CH_2_Cl_2_/NH_3_ (7.0 M in MeOH): 2, 3, 4, 5, 7, 8% NH_3_ (7.0 M in MeOH)) and thereby the desired flipped amide **1f** was obtained as a white solid (158 mg, 0.307 mmol, 67% over 3 steps). *R*_f_ = 0.44 (CH_2_Cl_2_/NH_3_ (7.0 M in MeOH) 9:1, UV). ^1^H NMR (400 MHz, CD_3_OD): *δ* = 8.34 – 8.14 (m, 2H), 7.60 – 7.39 (m, 2H), 7.34 (d, *J* = 7.8 Hz, 1H), 7.24 – 7.13 (m, 4H), 6.85 (d, *J* = 7.8 Hz, 1H), 4.10 – 3.95 (m, 5H), 3.89 (t, *J* = 6.1 Hz, 2H), 3.61 (s, 2H), 3.28 (s, 2H), 3.02 – 2.83 (m, 4H), 2.70 (t, *J* = 6.0 Hz, 2H), 2.46 (s, 3H) ppm. ^13^C NMR (101 MHz, CD_3_OD): *δ* = 173.4, 156.7, 147.8, 139.0, 134.7, 133.0, 132.6, 130.6, 129.3, 127.9, 127.3 (2xC), 126.8, 125.9, 125.6, 123.2, 110.7, 104.0, 60.4, 56.0, 51.9, 51.2, 49.1 (coinciding with solvent signal), 43.1, 42.4, 24.5, 15.8 ppm. HPLC-MS (ESI): *m/z* = 516 [M+H]^+^, 538 [M+Na]^+^; retention time: 2.67 min. HRMS (ESI): *m/z* calcd. for [C_29_H_34_N_5_O_2_S]^+^ 516.2428, found 516.1545.

## Supporting information

ParameterStudies

## Abbreviations

AcOH: acetic acid
Boc: *tert*-butoxycarbonyl
DCC: *N,N’*-dicyclohexylcarbodiimide
DCE: 1,2-dichloroethane
DMAP: 4-(dimethylamino)pyridine
DMF: *N,N*-dimethylformamide
DMSO: dimethyl sulfoxide
EtOAc: ethyl acetate
eq: equivalent
MeOH: methanol
MsCl: methanesulfonyl chloride
rt: room temperature
TEA: triethylamine
THF: tetrahydrofuran.

## DATA AVAILABILITY

The authors declare that the data supporting the findings of this study are available within the paper. X-ray data are deposited in the Protein Data bank (access code: 8RIB). The refined structures of the 20000 ligands available in MISATO and refined with IPA are provided in the repository https://gitlab.com/siriius/ipa-ligands.git.

## CODE AVAILABILITY

In-pocket optimization is available from the *ULYSSES* release repository, https://gitlab.com/siriius/ulysses.git.

## ACKNOWLEDGEMENTS

This work was supported by the Bundesministerium fuer Bildung und Forschung (BMBF), project SUPREME (number: 031L0268; to GMP) and by National Science Centre, Poland grant number 2020/39/B/NZ1/01551 (to GD).

## AUTHOR CONTRIBUTIONS

All authors have approved the final version of the manuscript and were involved in the reviewing and editing of the manuscript. The conceptualization of the work and the delineation of methodology were done by FM (computational) and VN (experimental). FM wrote the software. Validation was done by FM (computational) as well as VN, JDJO, GD, and GMP (experimental). The synthesis of the new inhibitors was performed by TF and SR. The original draft was prepared by FM and VN, who, together with GMP, were responsible for project supervision. Administration and funding acquisition was done by GMP and GD.

## ADDITIONAL INFORMATION

**Correspondence** and requests for materials should be addressed to FM or GMP.

