## Supplementary Information for "Quantum Chemistry in a Pocket: A Multifaceted Tool to Link Structure and Activity"

**S1 – Example of RMSD Constraint applied to the Lennard-Jones Potential**

| R | LJ | bias |
| --- | --- | --- |
| 0.7 | 5.31211 | 5.0802 |
| 0.75 | -0.535661 | -0.851858 |
| 0.8 | -2.08407 | -2.50664 |
| 0.85 | -2.21963 | -2.77319 |
| 0.9 | -1.93734 | -2.64812 |
| 0.95 | -1.57791 | -2.4725 |
| 1 | -1.25 | -2.35364 |
| 1.05 | -0.980114 | -2.31469 |
| 1.1 | -0.767053 | -2.34893 |
| 1.15 | -0.601765 | -2.43964 |
| 1.2 | -0.474308 | -2.56734 |
| 1.25 | -0.376036 | -2.71244 |
| 1.3 | -0.300034 | -2.85647 |
| 1.35 | -0.24097 | -2.98276 |
| 1.4 | -0.194806 | -3.07717 |
| 1.45 | -0.158498 | -3.12865 |
| 1.5 | -0.12976 | -3.12976 |
| 1.55 | -0.106869 | -3.07702 |
| 1.6 | -0.0885188 | -2.97089 |
| 1.65 | -0.0737201 | -2.81551 |
| 1.7 | -0.0617147 | -2.61815 |
| 1.75 | -0.0519201 | -2.38832 |
| 1.8 | -0.0438857 | -2.13691 |
| 1.85 | -0.0372608 | -1.87514 |
| 1.9 | -0.0317708 | -1.61365 |
| 1.95 | -0.0271998 | -1.36177 |
| 2 | -0.0233765 | -1.12701 |
| 2.05 | -0.0201647 | -0.914757 |
| 2.1 | -0.0174554 | -0.728239 |
| 2.15 | -0.015161 | -0.56872 |
| 2.2 | -0.0132104 | -0.435786 |
| 2.25 | -0.0115462 | -0.327744 |
| 2.3 | -0.0101213 | -0.242035 |
| 2.35 | -0.00889722 | -0.175626 |
| 2.4 | -0.00784233 | -0.125334 |
| 2.45 | -0.00693043 | -0.088086 |
| 2.5 | -0.00613981 | -0.0610867 |

**Figure S1.1:** Plot of the 12-6 Mie potential (Lennard-Jones) and a bias variant thereof.

As evidenced by the picture above, adding the bias potential creates a new minimum on the energy surface and minimally affects the original minimum. In the case of in-pocket optimization, the minimum of the native energy surface is anyway uninteresting since we wish to study the properties of an out-of-equilibrium structure (from the perspective of the free species). The depth and position of the new minimum depend on the reference structure and the parameters used.

Deviations are expected in the position of the actual minima because in-pocket optimization always includes some relaxation from all the atoms in the ligand, leading to small cumulative effects. Additionally, the parameter $N$ in equation (1) of the main manuscript will impact slightly the strength of the bias potential and, therefore also the relative position of all minima.

**S2 – Benchmarking the Algorithm for Simple Cases**

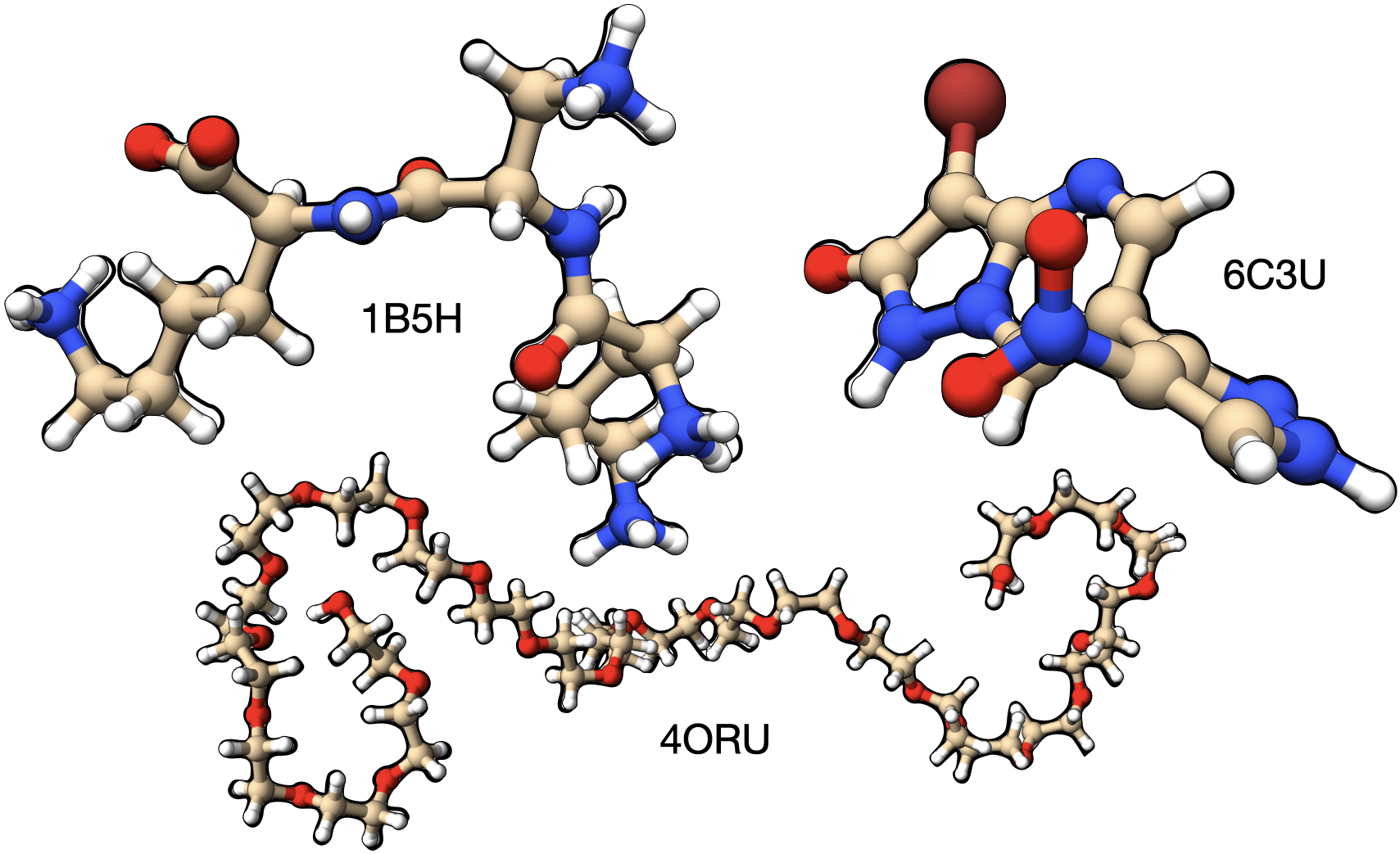

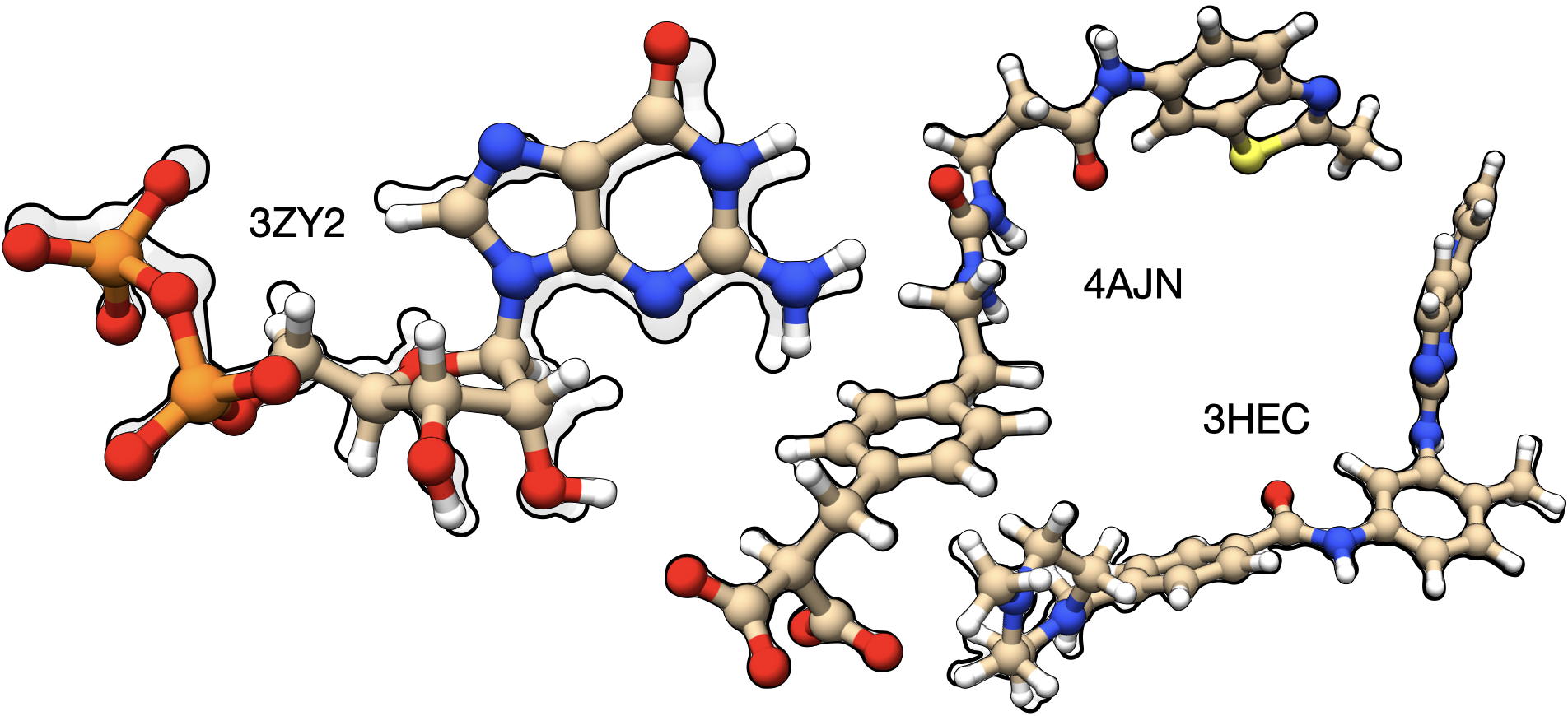

**Figure S2.1:** Structures of the 6 ligands we considered as downloaded from the MISATO database. The silhouette in the background represents the original ligand structure, and the colored molecules show the in-pocket optimized structures.

The values of κ and α determine primarily how much the original atomic positions are relaxed, and secondarily how much the conformation is changed. From a conceptual viewpoint this means that, for most potentials, the new minimum is always very close to the reference structure. For our test calculations, we selected 6 ligands from the PDB database: 1B5H [59], 3HEC [60], 3ZY2 [61], 4AJN [62], 4ORU [63], and 6C3U [64]. These structures evidence different features, which range from simple to severe geometric distortions, unnatural conformers, and large flexibility. Figure S2.1 contains the initial structures of these ligands (silhouette in the background) with the protonation state defined in the MISATO database [21], compared against the respective in-pocket optimized structures.

To determine whether a single set of parameters may be used, we begin our analysis with 6C3U. This ligand contains disruptions of π delocalization between the rings and the nitro group. Furthermore, the latter is unphysically distorted: one of the NO (NO1) bonds is 1.405 Å long, whereas the other is contracted to 1.178 Å (NO2). Besides the asymmetry, these values strongly disagree with experimental data [69] for nitromethane, 1.224 Å, and nitrobenzene, 1.223 Å.

| 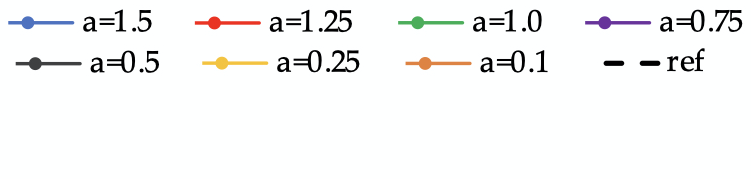 | 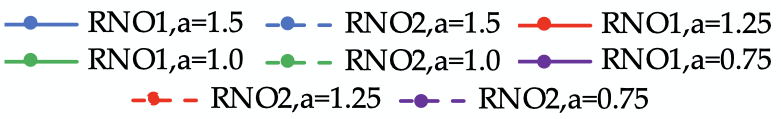 |
| --- | --- |
| (**a**) | (**b**) |

Dihedral (°)

| κ (Eh) | a=1.5 | a=1.25 | a=1.0 | a=0.75 | a=0.5 | a=0.25 | a=0.1 |
| --- | --- | --- | --- | --- | --- | --- | --- |
| 1 | 84 | 84 | 84 | 84 | 84 | 84 | 84 |
| 0.05 | 83.4 | 83.5 | 83.4 | 83.2 | 82.9 | 81.5 | 76.6 |
| 0.025 | 83.2 | 83.1 | 82.9 | 82.3 | 81.5 | 78.4 | 64.2 |
| 0.01 | 81.9 | 81.5 | 80.8 | 79.5 | 76.5 | 63.9 | 38.4 |
| 0.0075 | 81.2 | 80.5 | 79.5 | 77.5 | 72.7 | 55.3 | 32.1 |
| 0.005 | 79.4 | 78.3 | 76.4 | 72.6 | 63.5 | 44 | 25.5 |
| 0.0025 | 72.3 | 68.5 | 62.6 | 53.8 | 42.8 | 27.8 | 15.7 |
| 0.001 | 43.1 | 37.9 | 33.5 | 28.4 | 23.9 | 14.9 | 9.7 |
| 0 | -0.7 | -0.7 | -0.7 | -0.7 | -0.7 | -0.7 | -0.7 |

| κ (Eh) | ref |
| --- | --- |
| 0.05 | 84 |
| 0.025 | 84 |
| 0.01 | 84 |
| 0.0075 | 84 |
| 0.005 | 84 |
| 0.0025 | 84 |
| 0.001 | 84 |

RNO (Å)

| κ (Eh) | RNO1,a=1.5 | RNO2,a=1.5 | RNO1,a=1.25 | RNO2,a=1.25 | RNO1,a=1.0 | RNO2,a=1.0 | RNO1,a=0.75 | RNO2,a=0.75 |
| --- | --- | --- | --- | --- | --- | --- | --- | --- |
| 0.1 | 1.405 | 1.178 | 1.405 | 1.178 | 1.405 | 1.178 | 1.405 | 1.178 |
| 0.05 | 1.239 | 1.217 | 1.236 | 1.216 | 1.232 | 1.217 | 1.228 | 1.217 |
| 0.025 | 1.228 | 1.217 | 1.226 | 1.217 | 1.224 | 1.217 | 1.222 | 1.217 |
| 0.01 | 1.221 | 1.217 | 1.22 | 1.218 | 1.219 | 1.218 | 1.218 | 1.218 |
| 0.0075 | 1.219 | 1.218 | 1.219 | 1.218 | 1.218 | 1.218 | 1.218 | 1.218 |
| 0.005 | 1.218 | 1.218 | 1.218 | 1.218 | 1.218 | 1.218 | 1.218 | 1.219 |
| 0.0025 | 1.218 | 1.219 | 1.218 | 1.219 | 1.218 | 1.22 | 1.22 | 1.222 |
| 0.001 | 1.221 | 1.224 | 1.221 | 1.224 | 1.222 | 1.224 | 1.222 | 1.225 |
| 0 | 1.226 | 1.226 | 1.226 | 1.226 | 1.226 | 1.226 | 1.226 | 1.226 |

**Figure S2.2:** a) Change of dihedral angle for the atoms represented as spheres according to several combinations of the κ and α (a) parameters. b) Change of $R_{NO}$ bond distances according to several combinations of the κ and α (a) parameters.

Figure S2.2 a) shows changes in a dihedral associated with the nitro group according to different parametrizations of the bias potential. The experimental value for this dihedral is 84°, whereas with unconstrained optimization - the free ligand in water’s dielectric - the dihedral is -0.7°. For the largest κ tested, 0.05 E_h_, results for α ≥ 0.5 Å^-2^ are consistent and deviations in the dihedral are of at most 1.1°. For κ = 0.025 E_h_, α must take values of at least 1.0 Å^-2^ to ensure a similar quality. Figure 3 b) shows the change in the NO distances for the same combinations of κ but α ≥ 0.75 Å^-2^. Though not entirely obvious from the scale of the plot, we stress that irrespective of the parameters, in-pocket structures are far superior to the original bond lengths. $R_{NO1}$ is reasonably close to reference data [69] and also to the minimum from the GFN2-xTB surface. The improvement of $R_{NO2}$ is not as spectacular, however, the initial bond length was already acceptable (δ ≈ 0.04 Å). From the joint analysis of the two plots above, we would suggest using κ and α in the range of

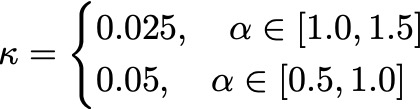

Strain or deformation energies ($E_{strain}$, Figure 4) are also used to evaluate the quality of in-pocket optimized structures. The calculated stress of the experimental structure is 51.453 kcal/mol, which is extremely large for such a small molecule with rotated single bonds between aromatic rings and a nitro group. The total tension in in-pocket optimized geometries lies between 10.743 and 12.093 kcal/mol (11.257 ± 0.837 kcal/mol), in much better agreement with the expected literature values [23].

Estrain (kcal/mol)

| κ (Eh) | a=1.5 | a=1.25 | a=1.0 | a=0.75 | a=0.5 | a=0.25 | a=0.1 |
| --- | --- | --- | --- | --- | --- | --- | --- |
| 0.05 | 13.2297 | 12.7029 | 12.0933 | 11.4376 | 10.7434 | 9.93108 | 9.07408 |
| 0.025 | 11.4266 | 11.1029 | 10.7362 | 10.3491 | 9.92799 | 9.30776 | 7.51713 |
| 0.01 | 10.0933 | 9.91869 | 9.71727 | 9.46788 | 9.06431 | 7.48693 | 3.95731 |
| 0.0075 | 9.81824 | 9.65603 | 9.46434 | 9.19225 | 8.61317 | 6.26192 | 3.18909 |
| 0.005 | 9.45722 | 9.29144 | 9.05179 | 8.59998 | 7.43186 | 4.68544 | 2.47996 |
| 0.0025 | 8.56121 | 8.099 | 7.31112 | 6.05023 | 4.53383 | 2.73816 | 1.54578 |
| 0.001 | 4.57355 | 3.89742 | 3.35414 | 2.82172 | 2.29428 | 1.4753 | 0.913478 |
| 0 | 0 | 0 | 0 | 0 | 0 | 0 | 0 |

**Figure S2.3:** Change in strain energy ($E_{strain}$) for 6C3U according to the parameters used in in-pocket optimization.

Data for other ligands is available below. Irrespective of the ligand, similar observations were made. Based on the data collected, we conclude that κ = 0.025 E_h_ and α = 1.0 Å^-2^ should provide a good choice for general use. For more conservative choices, we recommend using α = 1.5 Å^-2^.

**S3 – Parameter study for other ligands**

Analysis data for 1B5H:

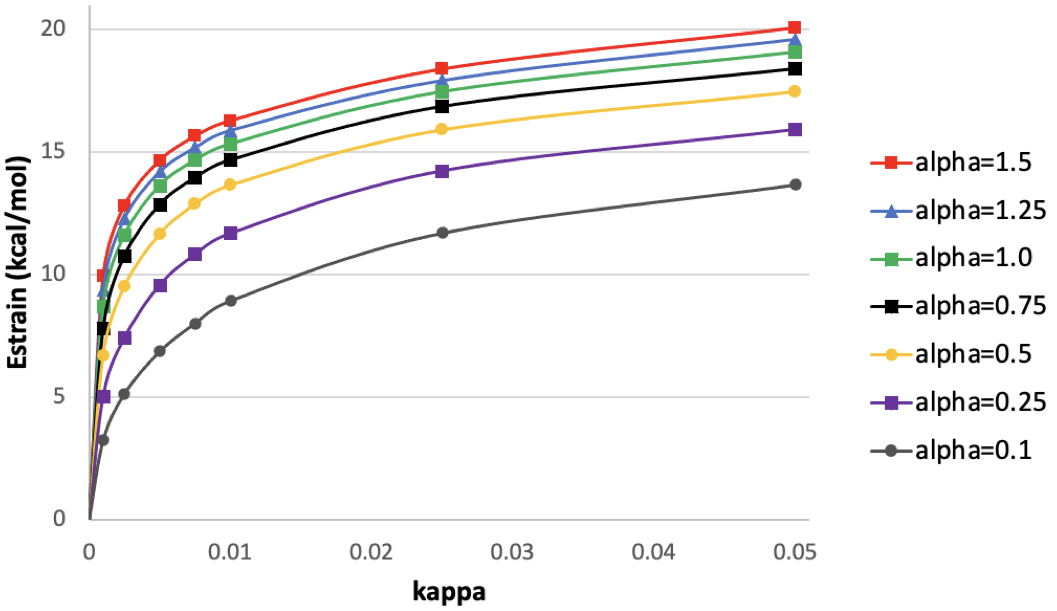

**Figure S3.1:** Change in strain energy (E_strain_) for 1B5H according to the parameters used in in-pocket optimization.

1B5H leads to the same conclusions as the main test case, 6C3U. The use of in-pocket optimization reduces tension, the decrease is however also not as significant because there are no distorted bonds and angles: from 29.335 kcal/mol, tension lowers to 18.116 ± 0.957 kcal/mol. The RMSD changes between 0.019 and 0.027 Å. This should be compared against the values for 6C3U, which go from 0.044 to 0.052. The accompanying file ParameterStudy_1b5h.xlsx contains more information on the calculations performed for this ligand.

Analysis data for 3HEC:

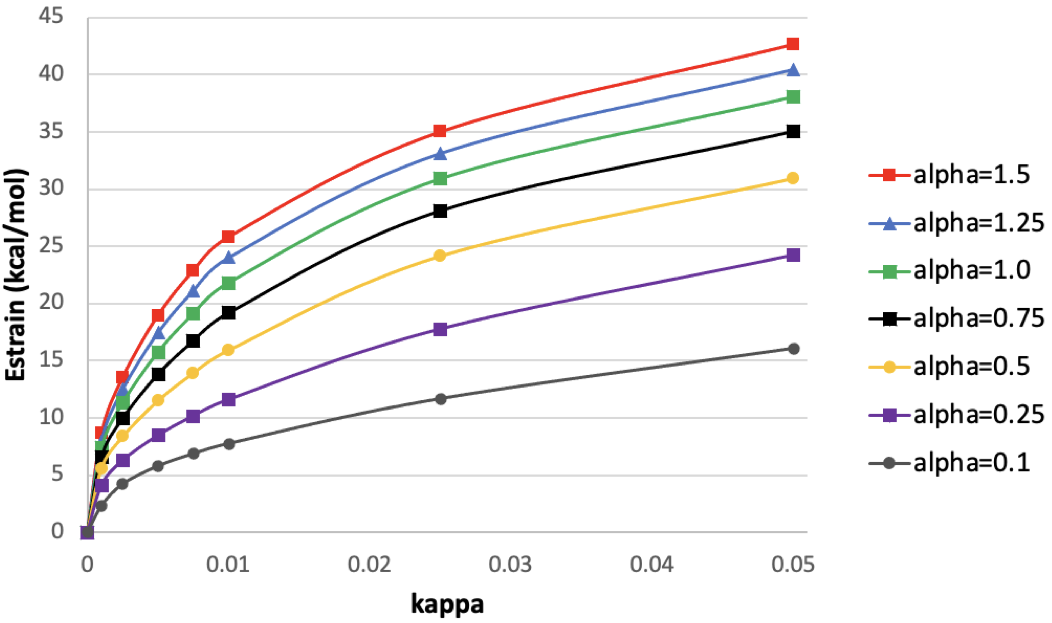

**Figure S3.2:** Change in strain energy (E_strain_) for 3HEC according to the parameters used in in-pocket optimization.

The structure of 3HEC has significantly more tension accumulated in the original geometry. According to our calculations, this takes values around 85.91 kcal/mol. Despite significant strain relief, the final deformation energies are still quite large, at about 33.837 ± 4.205 kcal/mol. The RMSD to the initial structure is between 0.038 and 0.053. The main problem associated with 3HEC is that the two tertiary amines have initially almost trigonal planar geometries. It seems thus that despite the 50 kcal/mol improvement in strain energy, improving the bond angles might not be as efficient as the improvements in bond lengths. The accompanying file ParameterStudy_3hec.xlsx contains more information on the calculations performed for this ligand.

Analysis data for 3ZY2:

3ZY2 is the most extreme case we considered. There are bonds so stretched that several graphical programs like Chimera [18] and Avogadro [50] consider them broken. On the other hand, some bonds are considerably shortened. With in-pocket optimization, we could reduce the strain energy from 152.745 kcal/mol to 76.023 ± 23.414 kcal/mol. The large error associated with the strain energy is evidence of the low quality of the structure deposited in the PDB. The strain energy that remained despite in-pocket optimization shows that further improvements are still possible. More acceptable deformation energies (below 20 kcal/mol) are obtained for κ = 0.01 and α ≤0.5, which are too weak of constraints. The RMSD for the sets of parameters we determined above varies between 0.158 and 0.196, which reflects the analysis above.

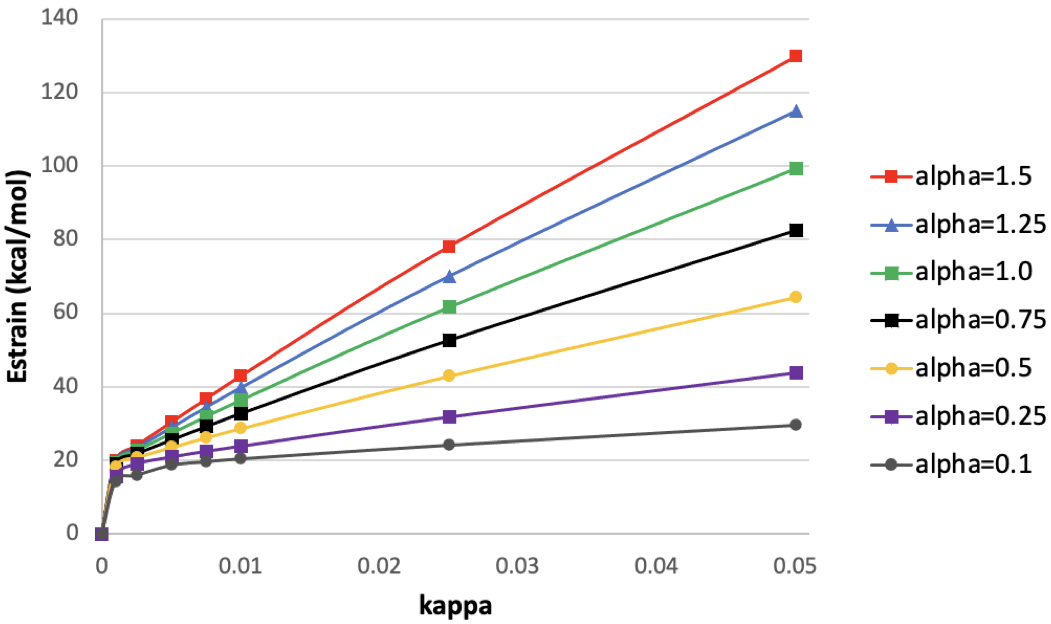

**Figure S3.3:** Change in strain energy (E_strain_) for 3ZY2 according to the parameters used in in-pocket optimization.

The accompanying file ParameterStudy_3zy2.xlsx contains more information on the calculations performed for this ligand.

Analysis data for 4AJN:

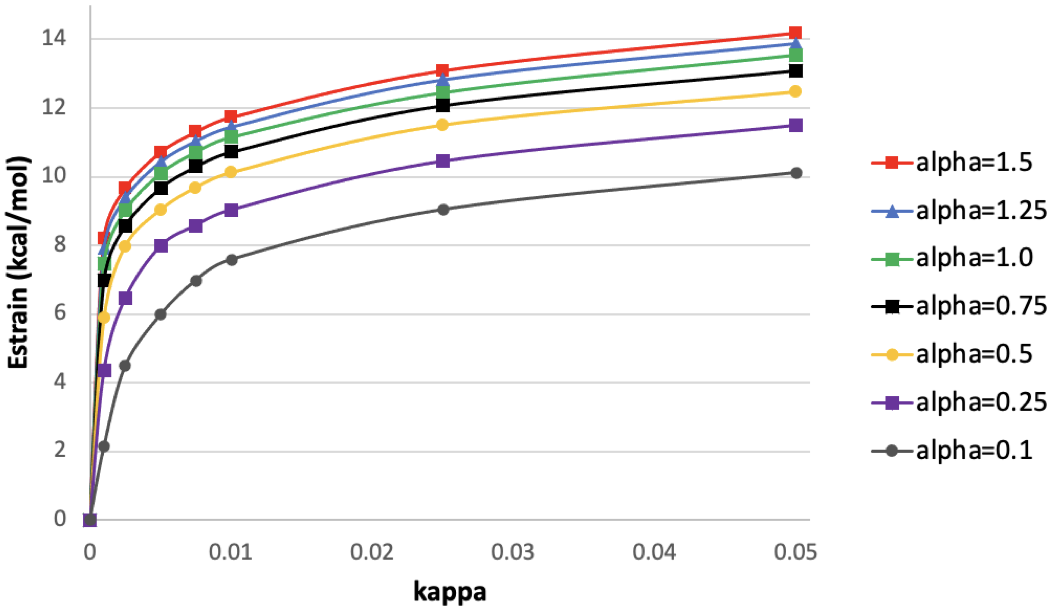

**Figure S3.4:** Change in strain energy (E_strain_) for 4AJN according to the parameters used in in-pocket optimization.

With 4AJN we return to structures that did not have much tension accumulated. The original strain energy was around 23.212 kcal/mol and after in-pocket optimization, we reduced this value to 12.915 ± 0.639 kcal/mol. The RMSD varies between 0.016 and 0.022. The accompanying file ParameterStudy_4ajn.xlsx contains more information on the calculations performed for this ligand.

Analysis data for 4ORU:

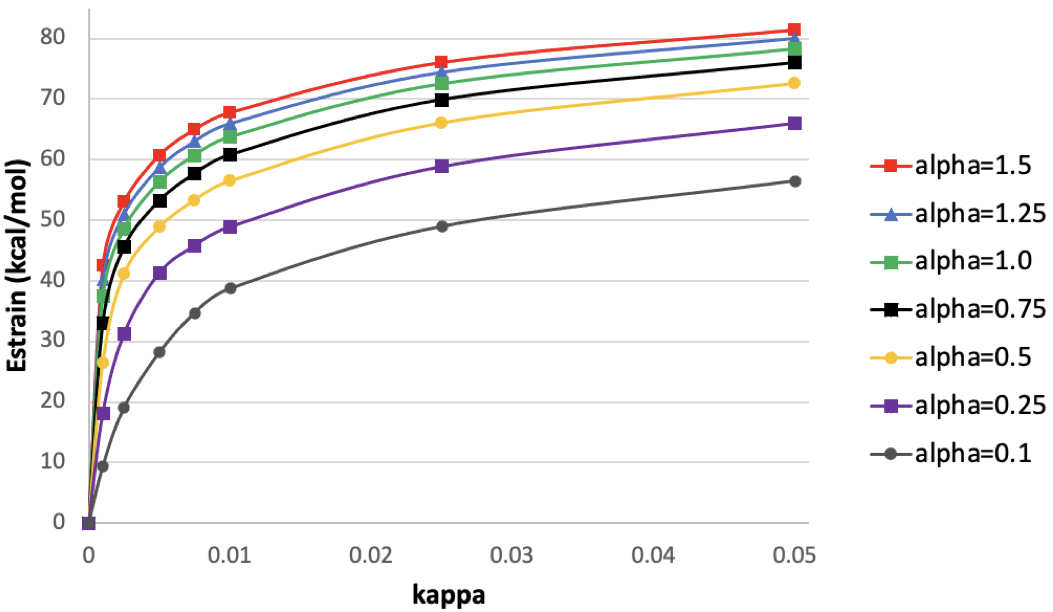

**Figure S3.5:** Change in strain energy (E_strain_) for 4ORU according to the parameters used in in-pocket optimization.

The original strain energy for 4ORU is 96.604 kcal/mol. With the parameters we defined, the strain lowers to 74.938 ± 3.325 kcal/mol. Again, using more relaxed thresholds might lower strain energies down to almost 10 kcal/mol, but this requires the lowest combinations for α (0.1) and κ (0.001). Because there are no obvious deformations on the structure, we believe that the high strain energy is caused by the dimensions of the ligand: 178 atoms and a molecular weight of 1119.33 g/mol. The accompanying file ParameterStudy_4oru.xlsx contains more information on the calculations performed for this ligand.

Analysis data for 6C3U:

The accompanying file ParameterStudy_6c3u.xlsx contains more information on the calculations performed for this ligand.

**S4 – Iterative Approach**

Despite the significant improvements brought by in-pocket optimization, structures that were initially heavily distorted still exhibited significant tension after optimization. Though visually the geometries seem acceptable, the accumulated tension is still overly large. This is the case of 3HEC and 3ZY2. An easy fix is to relax the bias potential, leading to loss of control in the optimization and lack of generality that we seek for the algorithm. In less extreme cases the binding conformation may be abnormally impaired or even lost. Here we investigate whether running a series of successive in-pocket optimization calculations could be useful. This is also of relevance when determining how stable structures are for further optimization.

6C3U

| Iteration | Estrain | dihedral | RMSD | RNO1 | RNO2 |
| --- | --- | --- | --- | --- | --- |
| 0 | 51.4534 | 84 | 0 | 14.05 | 11.78 |
| 1 | 10.7362 | 82.9 | 51.6879 | 12.24 | 12.17 |
| 2 | 9.69669 | 82 | 15.2846 | 12.16 | 12.18 |
| 3 | 9.39137 | 80.7 | 9.76163 | 12.16 | 12.18 |
| 4 | 9.22512 | 79.5 | 7.36553 | 12.16 | 12.18 |
| 5 | 9.0932 | 78.2 | 6.71734 | 12.16 | 12.18 |
| 6 | 8.95204 | 76.7 | 7.37721 | 12.16 | 12.18 |
| 7 | 8.80301 | 75.3 | 7.58637 | 12.16 | 12.19 |
| 8 | 8.63369 | 73.7 | 8.06636 | 12.17 | 12.19 |
| 9 | 8.44415 | 72.1 | 8.60419 | 12.17 | 12.19 |
| 10 | 8.24333 | 70.4 | 8.70121 | 12.17 | 12.19 |
| 11 | 8.02595 | 68.6 | 9.17145 | 12.17 | 12.2 |

3ZY2

| Iteration | Estrain | RMSD |
| --- | --- | --- |
| 0 | 152.745 | 0 |
| 1 | 61.6508 | 195.62 |
| 2 | 36.6193 | 58.1573 |
| 3 | 30.0717 | 33.4374 |
| 4 | 26.9898 | 24.1589 |
| 5 | 25.1456 | 19.1252 |
| 6 | 23.9184 | 15.7524 |
| 7 | 23.057 | 13.1749 |
| 8 | 22.4127 | 11.4809 |
| 9 | 21.9252 | 9.99021 |
| 10 | 21.5446 | 8.84837 |
| 11 | 21.253 | 7.59785 |
| 12 | 20.9927 | 7.5968 |
| 13 | 20.798 | 6.20691 |
| 14 | 20.6332 | 5.733 |
| 15 | 20.4905 | 5.36383 |
| 16 | 20.3676 | 4.96893 |
| 17 | 20.2643 | 4.43678 |
| 18 | 20.1644 | 4.56332 |
| 19 | 20.0795 | 4.10842 |
| 20 | 20.0051 | 3.76606 |
| 21 | 19.9346 | 3.73527 |
| 22 | 19.8616 | 4.04152 |
| 23 | 19.8027 | 3.36211 |
| 24 | 19.7476 | 3.2636 |
| 25 | 19.6948 | 3.21479 |
| 26 | 19.6384 | 3.55903 |
| 27 | 19.5919 | 2.99104 |
| 28 | 19.547 | 2.95475 |
| 29 | 19.5033 | 2.93698 |
| 30 | 19.4609 | 2.90441 |
| 31 | 19.4195 | 2.87538 |
| 32 | 19.4155 | 0.27758 |
| 33 | 19.4117 | 0.270201 |
| 34 | 19.4078 | 0.269738 |
| 35 | 19.404 | 0.269348 |
| 36 | 19.4002 | 0.268967 |
| 37 | 19.3964 | 0.268591 |
| 38 | 19.3926 | 0.268221 |
| 39 | 19.3888 | 0.267855 |
| 40 | 19.385 | 0.267494 |
| 41 | 19.3812 | 0.267136 |

**Figure S4.1:** Using a series of in-pocket optimizations to optimize the structures of two ligands, 6C3U (left) and 3ZY2 (right). Strain energies in kcal/mol, dihedral angles in degrees, RMSDs and NO bond distances in Å. RMSDs are multiplied by a factor of 1000 and NO bond distances by 10.

Expectedly, the dihedral angle of the nitro group in 6C3U (Figure S4.1, left) decreases monotonically with the iteration number. This is because the optimization target is updated every iteration. Furthermore, the optimized geometry gradually evolves to the free ligand. The strain energy is also reduced with the number of runs. However, in the case of 6C3U, major changes take place in the first run only. The same happens to both NO bond distances. The RMSD still shows significant changes in the second iteration, but a slow decrease follows immediately. We conclude thus that for a ligand with moderate deformations one run of in-pocket optimization suffices to stabilize most criteria. RMSDs might require more iterations to stabilize. Nevertheless, after two potential runs of in-pocket optimization, there is only degradation of the binding pose.

Despite the excessive unphysical deformation in 3ZY2, its general behavior is identical to that of 6C3U. Major structural updates take place in the first iterations, then strain and (later) RMSD stabilize. In this case, more runs of in-pocket optimization are required. After 15 iterations changes in RMSD between successive iterations are minor. For the strain energies, the same happens at about 7 runs. To show how slowly strain energies evolve with the number of in-pocket iterations, after 9 calls the strain energy is at 21.925 kcal/mol. At 30 iterations, the deformation energy is at 19.461 kcal/mol, and at 40 iterations this value lowered to 19.385 kcal/mol. However, we would not recommend going beyond 4-5 iterations if the intent is to calculate deformation energies: the improvements are minor afterward.

This analysis shows that for good experimental structures, a single run of in-pocket optimization suffices to remove most of the unphysical tension. Running successively the algorithm may lead to improvements, however, the only quantity that is sensitive enough to be used as a stopping criterion is the Cartesian RMSD to their reference structure. To avoid further degradation of the structures, it might be advisable to update the number of biasing structures, giving larger weights to the initial geometries. This path was however not explored in the current work.

**S5 – Excluding Protons**

Protein crystal structures provide an experimental basis for all atoms but the protons, which must be added *a posteriori*. The bias potential is built in such a way that protons may be excluded from the constraints. In those conditions, the latter are freely optimized. Table S5.1 compares strain energies for the 6 ligands selected using the MISATO geometry, the in-pocket one with all atoms constrained, and the in-pocket structure with free protons.

**Table S5.1:** Strain energies (in kcal/mol) for the 6 test cases, using the MISATO geometry and the in-pocket ones, with (all) and without (H free) constrained protons.

|  | **1B5H** | **3HEC** | **3ZY2** | **4AJN** | **4ORU** | **6C3U** |
| --- | --- | --- | --- | --- | --- | --- |
| MISATO | 29.335 | 85.915 | 152.745 | 23.212 | 96.604 | 51.453 |
| in-pocket (all) | 17.456 | 30.888 | 61.651 | 12.915 | 75.502 | 10.736 |
| in-pocket (H free) | 15.046 | 26.676 | 43.068 | 10.430 | 60.232 | 10.437 |

Except for 6C3U, which barely has protons (5 in total), there is a significant improvement in strain energies for all structures. Additional 18 kcal/mol are improved in a single in-pocket optimization run of 3ZY2 by simply excluding hydrogen atoms. 4ORU also contains significant tension accumulated, but this is due to the total number of hydrogens in the molecule (102). When this tension is normalized with the total number of protons, an average deformation energy of 0.118 kcal/mol per hydrogen is obtained. This clearly shows the cumulative effects of small unphysical deformation on the structures. 3HEC is a more complex case since the problems in the structure relate to two tertiary amine groups. The gain in freely optimizing the protons is about 4 kcal/mol, the penalty however is an increase of more than 20 iterations in the calculation.

Despite potential drawbacks in increased calculation times, excluding the protons from the in-pocket constraining potential seems highly advantageous. In particular, because the coordinates of the latter have no experimental basis whatsoever.

To test the selected parameters on ligands outside our benchmark, we ran in-pocket optimization to determine the tension applied by proteins on ibuprofen. We selected the structures 1EQG [70], 2BXG [71], 2PWS [72], and 2WD9 [73], and treated all molecules of ibuprofen available in those complexes. Before any optimization, we obtained an average strain energy of 22.756 (9.458) kcal/mol, which compares reasonably well with the results of Merz and coworkers [19] (average of 28.1 kcal/mol). We note that the differences may stem from the method used - semi-empirical Vs. *ab initio* - but also due to the solvation model. All factors together may also impact the local equilibrium structures used to estimate tension. The total deformation energy is reduced to 4.725 (1.969) kcal/mol when in-pocket optimization is used along with free protons. This value should be compared against the 3.7 kcal/mol reported in the literature [19].

**S6 – Comparison of Dipole Moments for Ligands 6C3U and 3ZY2**

To evaluate the effects on the charge properties of ligands we calculated the dipole moment vectors and their respective norms for 6C3U and 3ZY2 according to the three sets of geometries available: the initial MISATO, the in-pocket optimized structure, and the equilibrium structures of the ligands.

6C3U

|  | PDBBind | Pocket | Free |
| --- | --- | --- | --- |
| 1 | 2.27517 | 2.25725 | 2.42951 |
| 2 | -2.37621 | -1.89431 | -0.790685 |
| 3 | 1.44103 | 1.5163 | 1.42206 |
| 4 | 3.59157 | 3.31403 | 2.92403 |

3ZY2

|  | PDBBind | Pocket | Free |
| --- | --- | --- | --- |
| 1 | 7.04586 | 10.1931 | 1.46882 |
| 2 | 7.38033 | 10.0223 | 7.16737 |
| 3 | 4.06212 | 4.66825 | -1.71681 |
| 4 | 10.9824 | 15.0379 | 7.51505 |

**Figure S6.1:** Comparison of the dipole moment vectors and their norms for 6C3U and 3ZY2.

The case of 6C3U shows the typical behavior observed for ligands that are not overly deformed: most charges remain unaffected by in-pocket or even free optimization and most components of the dipole moment vector are unaffected. Differences come in the most distorted groups, and these might lead to differences in one of the components of the vector due to changes in coordinates. Nevertheless, the total dipole moment is barely affected, which indicates that the electrostatics and polarization forces between molecules should be identical. 3ZY2 shows an extreme case, and it shows marked differences in most partial charges. Not only the relative orientation of the ligand in the protein should be affected (all components of the dipole moment vector change) but also the electrostatics and polarization must differ since the total dipole moment is different.

**S7 – Geometry Comparison for Compound 2**

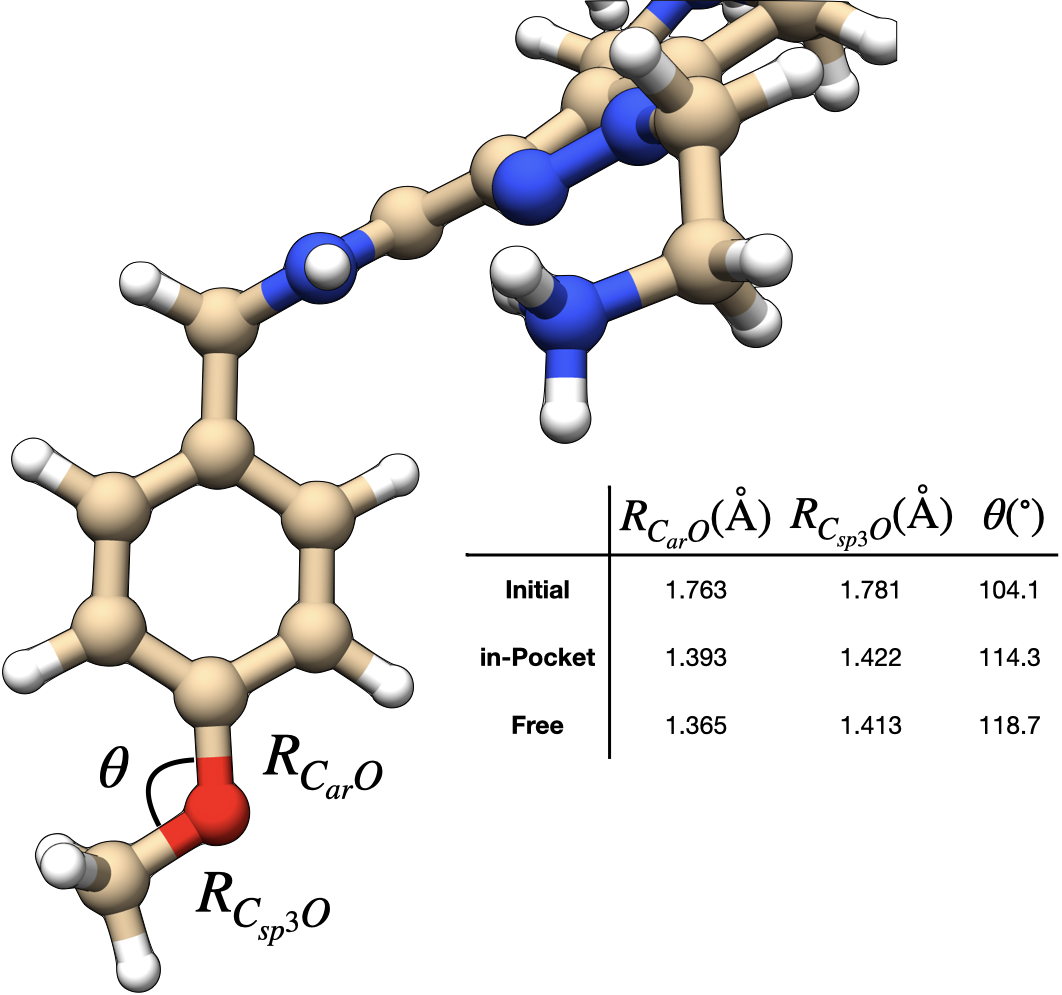

**Figure S7.1:** Comparison of the geometries for compound **2** in 3 stages: original structure pre-in-pocket optimization; in-pocket optimized; freely optimized molecule. Comparison focused strictly on the ether functionality.

The initial structure we used in this study started from the experimental coordinates, where in place of the oxygen atom there was a sulfur. Consequently, the starting coordinates reflect the thioether functional group. In-pocket optimization brings in this case both CO bond distances closer to the freely optimized one. The COC angle is in this case also refined by in-pocket, bringing the value closer to what is expected for an ether group.

**S8 – Effects of Dielectric**

The calculations discussed in the main document use water as a dielectric medium. The dielectric constant in a protein is however estimated to be significantly different than the one in bulk water. At the protein’s surface this value can take values of 20-30, which are up to 4 times smaller than in bulk water. To test the effect of dielectricity on the final geometries we ran in-pocket optimization using acetone instead (with ALPB dielectric constant of 20.7). From the practical point of view the structures in acetone were identical to the ones in water’s dielectric. The difference might however come in the energetics. Though for most structures the differences in strain energy were marginal, in the case of 3ZY2 and 1B5H deviations were more significant. The accompanying Excel files contain the results of our investigations.

**S9 – In-Pocket Optimization on 2 Å Resolution Structure of compound 1 in PEX14.**

To test the usefulness of in-pocket optimization on crystal structures of slightly lower resolution we solved the structure of compound **1** in PEX 4 at 2 Å resolution. Here we extracted one of the bound ligands and we used in-pocket to refine the structure and add hydrogen atoms. Figure S16.1 shows the atom numbering of the ligand, and Table S16.1 shows the bond distances for the high-resolution structure (main manuscript), the “low”-resolution one (2 Å) and in-pocket. Deviations and statistics are provided in Table S16.2.

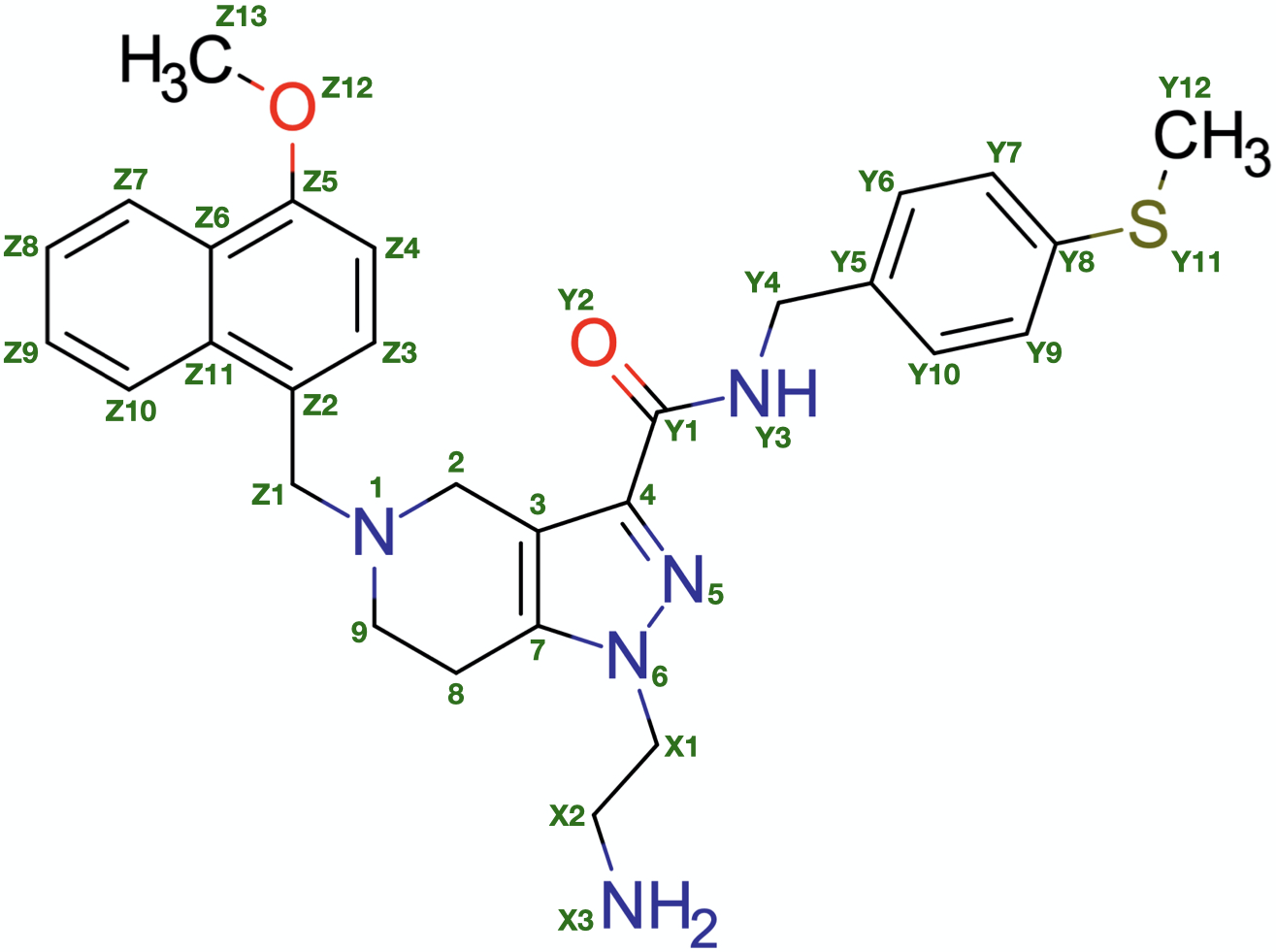

**Figure S9.1:** Atom numbering for compound **1**.

**Table S9.1:** Bond distances for the different experimental and in silico models of compound **1**. The in-pocket optimized structure considered no constraint whatsoever on hydrogen atoms. IP stands for in-pocket. Dev. is a deviation and |Dev.| is an absolute deviation.

|  |  |  | R_AB_ (Å) | | | | | | |
| --- | --- | --- | --- | --- | --- | --- | --- | --- | --- |
|  | A | B | High Res. | Low Res. | IP | Dev.  Low Res. | Dev. IP | \|Dev.\|  Low Res. | \|Dev.\| IP |
| Core | 1 | 2 | 1.46 | 1.48 | 1.47 | 0.02 | 0.01 | 0.02 | 0.01 |
|  | 2 | 3 | 1.49 | 1.51 | 1.49 | 0.02 | 0.00 | 0.02 | 0.00 |
|  | 3 | 4 | 1.36 | 1.36 | 1.40 | -0.01 | 0.03 | 0.01 | 0.03 |
|  | 3 | 7 | 1.35 | 1.34 | 1.38 | -0.01 | 0.03 | 0.01 | 0.03 |
|  | 4 | 5 | 1.35 | 1.34 | 1.33 | 0.00 | -0.01 | 0.00 | 0.01 |
|  | 5 | 6 | 1.35 | 1.36 | 1.34 | 0.01 | -0.01 | 0.01 | 0.01 |
|  | 6 | 7 | 1.37 | 1.36 | 1.35 | -0.01 | -0.02 | 0.01 | 0.02 |
|  | 7 | 8 | 1.48 | 1.48 | 1.47 | -0.01 | -0.01 | 0.01 | 0.01 |
|  | 8 | 9 | 1.53 | 1.53 | 1.54 | 0.00 | 0.02 | 0.00 | 0.02 |
|  | 9 | 1 | 1.47 | 1.49 | 1.46 | 0.02 | -0.01 | 0.02 | 0.01 |
| X | X1 | X2 | 1.51 | 1.53 | 1.53 | 0.01 | 0.01 | 0.01 | 0.01 |
|  | X2 | X3 | 1.47 | 1.50 | 1.48 | 0.03 | 0.01 | 0.03 | 0.01 |
| Y | Y1 | Y2 | 1.25 | 1.26 | 1.23 | 0.01 | -0.02 | 0.01 | 0.02 |
|  | Y1 | Y3 | 1.32 | 1.32 | 1.34 | 0.00 | 0.02 | 0.00 | 0.02 |
|  | Y3 | Y4 | 1.44 | 1.46 | 1.45 | 0.02 | 0.01 | 0.02 | 0.01 |
|  | Y4 | Y5 | 1.50 | 1.51 | 1.50 | 0.00 | 0.00 | 0.00 | 0.00 |
|  | Y5 | Y6 | 1.38 | 1.40 | 1.39 | 0.02 | 0.01 | 0.02 | 0.01 |
|  | Y6 | Y7 | 1.38 | 1.38 | 1.39 | 0.01 | 0.01 | 0.01 | 0.01 |
|  | Y7 | Y8 | 1.39 | 1.39 | 1.39 | 0.00 | 0.00 | 0.00 | 0.00 |
|  | Y8 | Y9 | 1.39 | 1.38 | 1.39 | -0.01 | 0.00 | 0.01 | 0.00 |
|  | Y9 | Y10 | 1.39 | 1.38 | 1.37 | -0.01 | -0.01 | 0.01 | 0.01 |
|  | Y10 | Y5 | 1.39 | 1.38 | 1.39 | -0.01 | 0.00 | 0.01 | 0.00 |
|  | Y8 | Y11 | 1.76 | 1.74 | 1.76 | -0.02 | 0.00 | 0.02 | 0.00 |
|  | Y11 | Y12 | 1.78 | 1.79 | 1.81 | 0.01 | 0.03 | 0.01 | 0.03 |
| Z | Z1 | Z2 | 1.52 | 1.52 | 1.52 | 0.00 | 0.00 | 0.00 | 0.00 |
|  | Z2 | Z3 | 1.35 | 1.38 | 1.37 | 0.03 | 0.03 | 0.03 | 0.03 |
|  | Z3 | Z4 | 1.40 | 1.40 | 1.40 | 0.00 | 0.00 | 0.00 | 0.00 |
|  | Z4 | Z5 | 1.37 | 1.36 | 1.37 | -0.02 | 0.00 | 0.02 | 0.00 |
|  | Z5 | Z6 | 1.42 | 1.43 | 1.43 | 0.01 | 0.01 | 0.01 | 0.01 |
|  | Z6 | Z7 | 1.40 | 1.42 | 1.41 | 0.02 | 0.01 | 0.02 | 0.01 |
|  | Z6 | Z11 | 1.44 | 1.43 | 1.43 | -0.01 | -0.01 | 0.01 | 0.01 |
|  | Z7 | Z8 | 1.35 | 1.37 | 1.37 | 0.02 | 0.02 | 0.02 | 0.02 |
|  | Z8 | Z9 | 1.39 | 1.40 | 1.40 | 0.01 | 0.01 | 0.01 | 0.01 |
|  | Z9 | Z10 | 1.35 | 1.36 | 1.37 | 0.01 | 0.02 | 0.01 | 0.02 |
|  | Z10 | Z11 | 1.41 | 1.43 | 1.42 | 0.02 | 0.01 | 0.02 | 0.01 |
|  | Z11 | Z2 | 1.43 | 1.43 | 1.43 | 0.00 | 0.00 | 0.00 | 0.00 |
|  | Z5 | Z12 | 1.36 | 1.38 | 1.37 | 0.01 | 0.01 | 0.01 | 0.01 |
|  | Z12 | Z13 | 1.41 | 1.43 | 1.42 | 0.02 | 0.01 | 0.02 | 0.01 |
| Core-X | 6 | X1 | 1.45 | 1.47 | 1.46 | 0.01 | 0.00 | 0.01 | 0.00 |
| Core-Y | 4 | Y1 | 1.45 | 1.45 | 1.47 | 0.00 | 0.02 | 0.00 | 0.02 |
| Core-Z | 1 | Z1 | 1.47 | 1.49 | 1.46 | 0.02 | -0.01 | 0.02 | 0.01 |

**Table S9.2:** Statistics for bond distance deviations presented in table S16.1.

|  | Dev. | | | | \|Dev.\| | | |
| --- | --- | --- | --- | --- | --- | --- | --- |
|  | Low Res. | | IP | Low Res. | | | IP |
| average | 0.006 | 0.005 | | | 0.012 | 0.012 | |
| st dev | 0.013 | 0.014 | | | 0.007 | 0.009 | |
| max | 0.029 | 0.035 | | | 0.029 | 0.035 | |

As evidenced by the data, the lower resolution of the crystal data did not negatively impact the structure of the ligand. Deviations are all below 0.03 Å and on average of about 0.01 Å. In such a situation, we also verify that in-pocket optimization using GFN2-xTB did not improve the ligand’s structure, because there was nothing to be improved. The average deviation and average absolute deviation between low-resolution crystal data and quantum chemical are on par with each other.

A closer look at the deviations shows that there is no systematic improvement of specific functionalities, not even for instance those related to the sulfur atom. The largest deviations on the in-pocket structure are all related to a specific carbon atom (4 in Figure S16.1). Removing this atom from the statistical analysis brings a slight shift that favors in-pocket optimization. We are nonetheless still speaking about minimal improvements. We did not investigate the reason behind the poorer results on carbon 4.

**S10 – The impact of (protein) Structural Elements over Bound conformations.**

Unconstrained geometry optimization is used to refine experimental structures and remove inaccuracies. While using full structures is feasible at lower levels of theory, for Density Functional or *ab initio* methods one requires smaller constructs containing the binding pocket or the first shells of residues. Here we analyze the impact of using smaller constructs on the integrity of the binding pocket.

We focus on the four structures in the asymmetric units of 5L8A and 5N8V, as well as the two structures in the asymmetric unit of 8RIB. The use of several protein-complexes from the same asymmetric unit gives us access to minimal statistical information over the same complexes. Two set of pockets were prepared for this analysis. We started by identifying all residues within 5 Å of the ligand. The first pocket contained only those residues capped at the N-terminus when necessary (nobbPocket). The second type of pocket consists of an extension of the first one, where the main chain atoms of clipped residues are included (bbPocket). This resulted in protein constructs containing 322 (nobb) and 386 (bb) atoms. Figure S10.1 contains a schematic representation of how we cut each type of pocket.

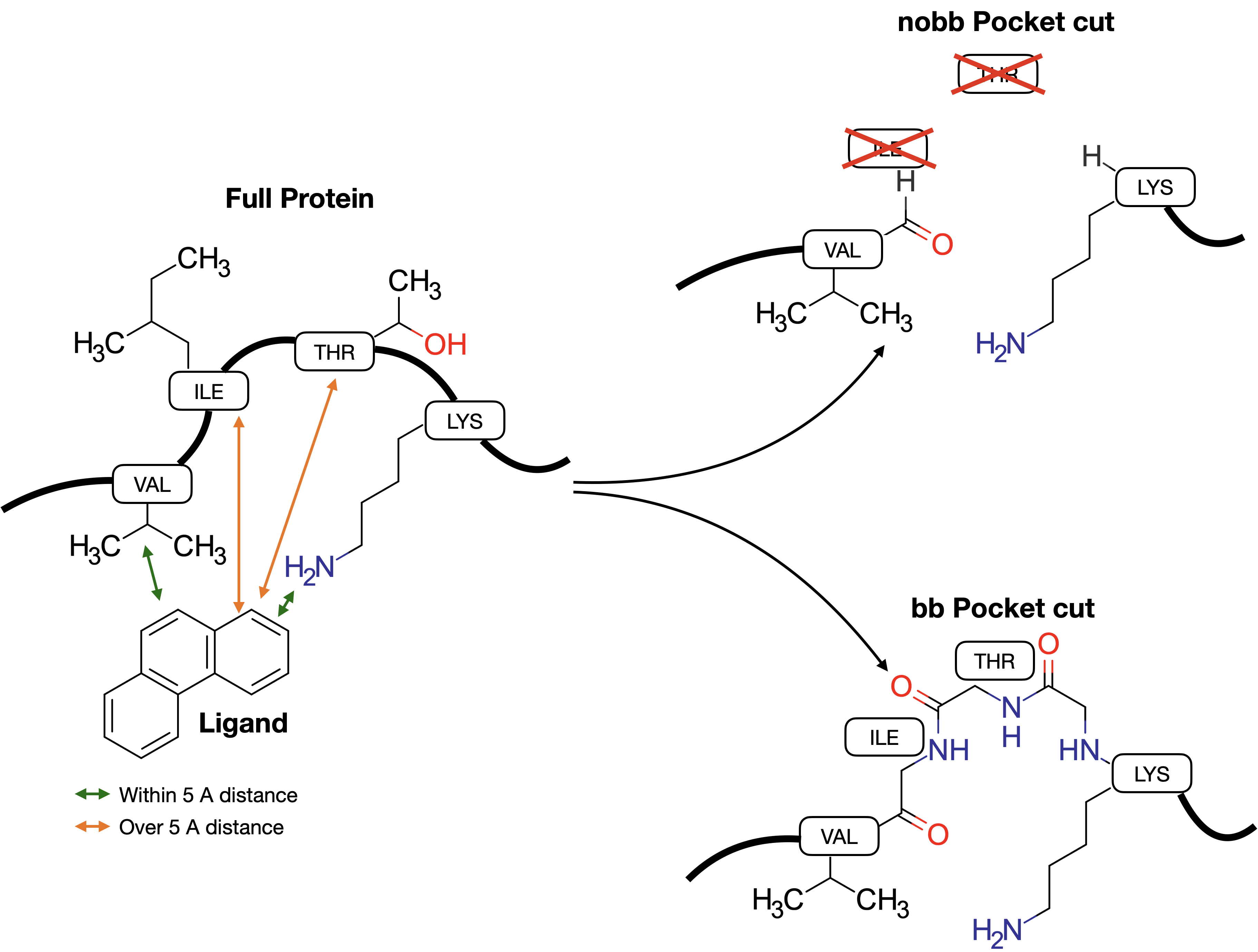

**Figure S10.1:** Schematic representation of the pocket cutting strategy for this evaluation.

Structures were optimized with In-Pocket and with free, unconstrained minimization for all cases (two pocket types, 10 complexes). Figure S10.2 compares the optimized structures for the three protein-ligand complexes, with chain A always selected. Statistical information on the respective RMSDs is provided in Figure S10.3.

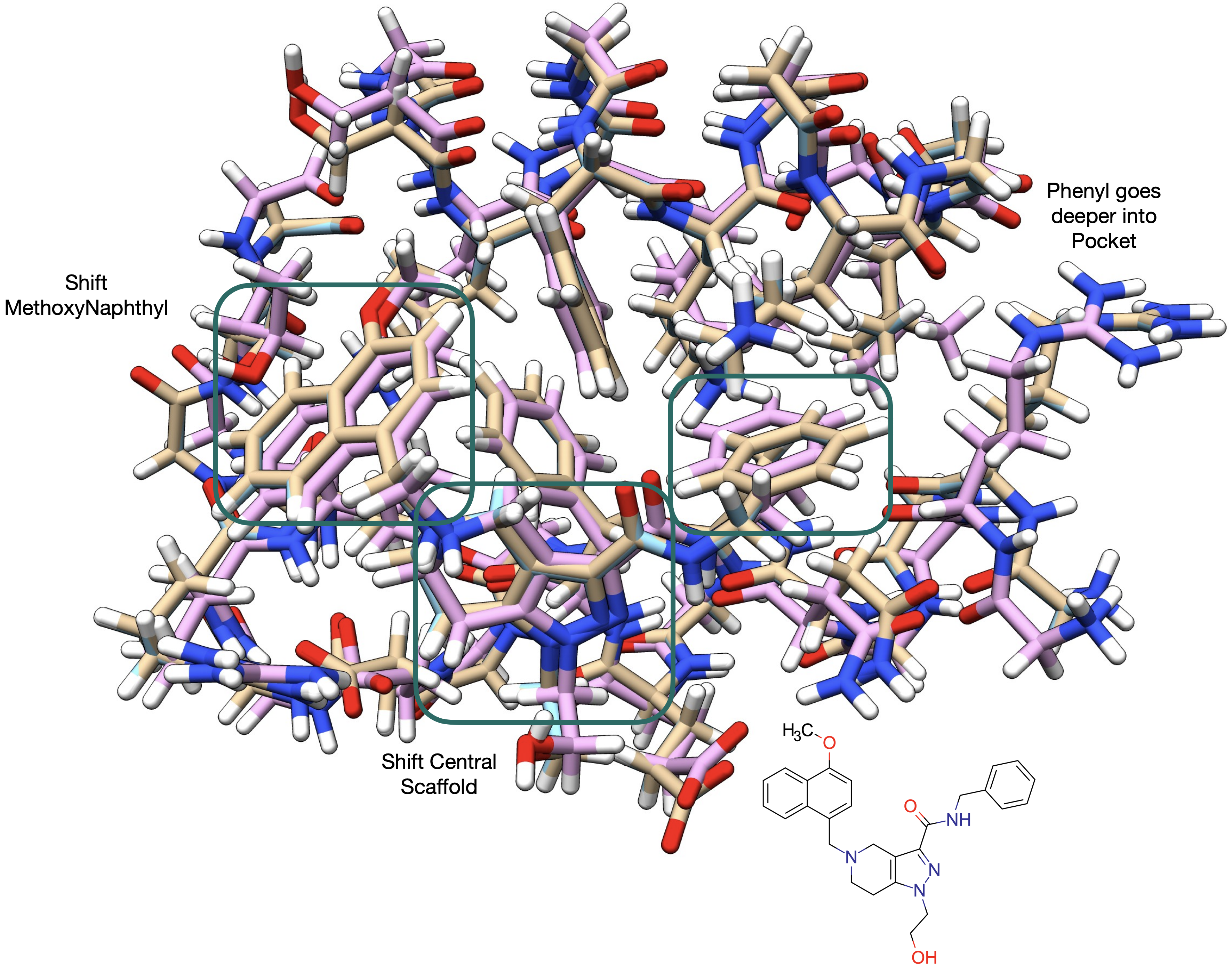

a

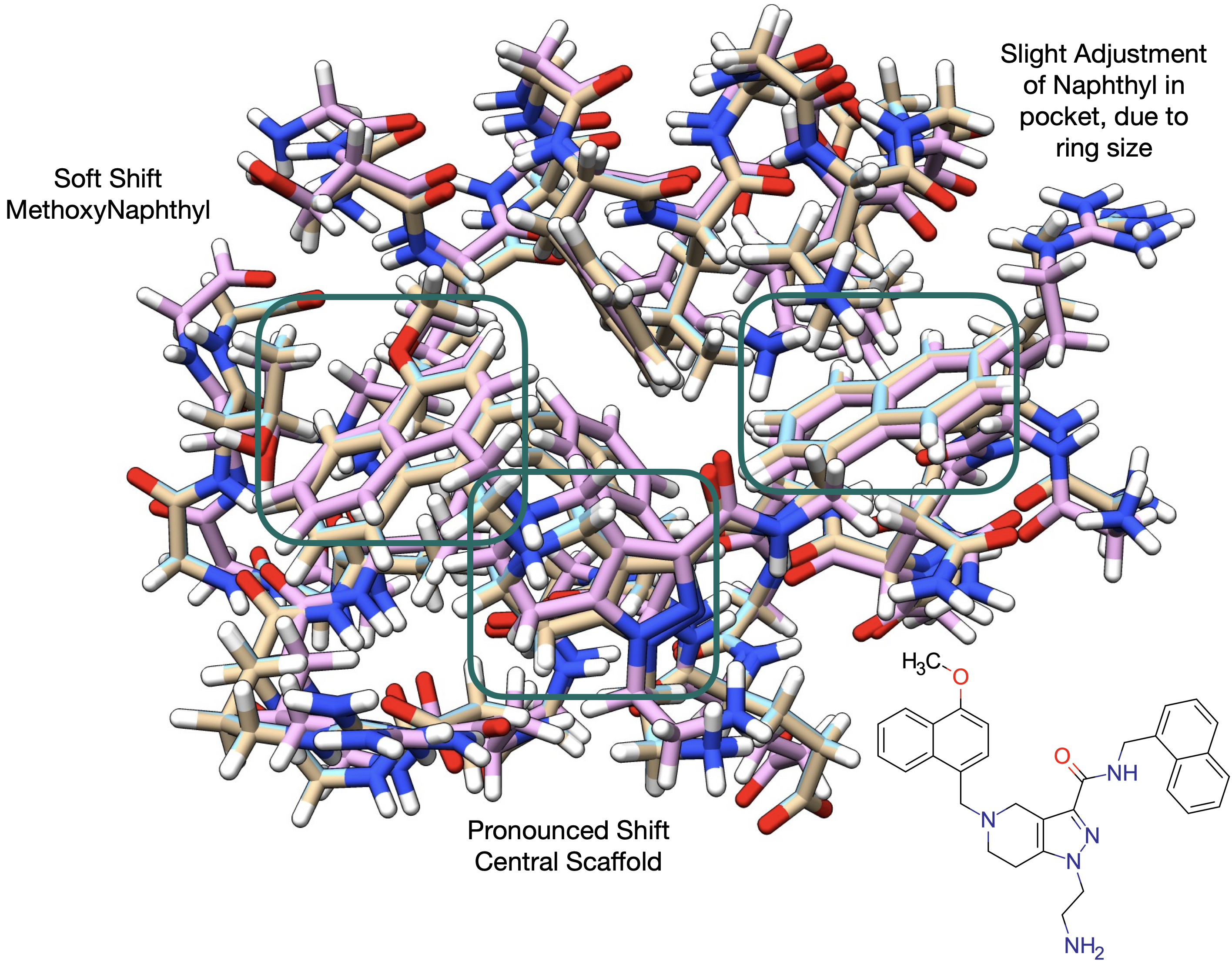

b

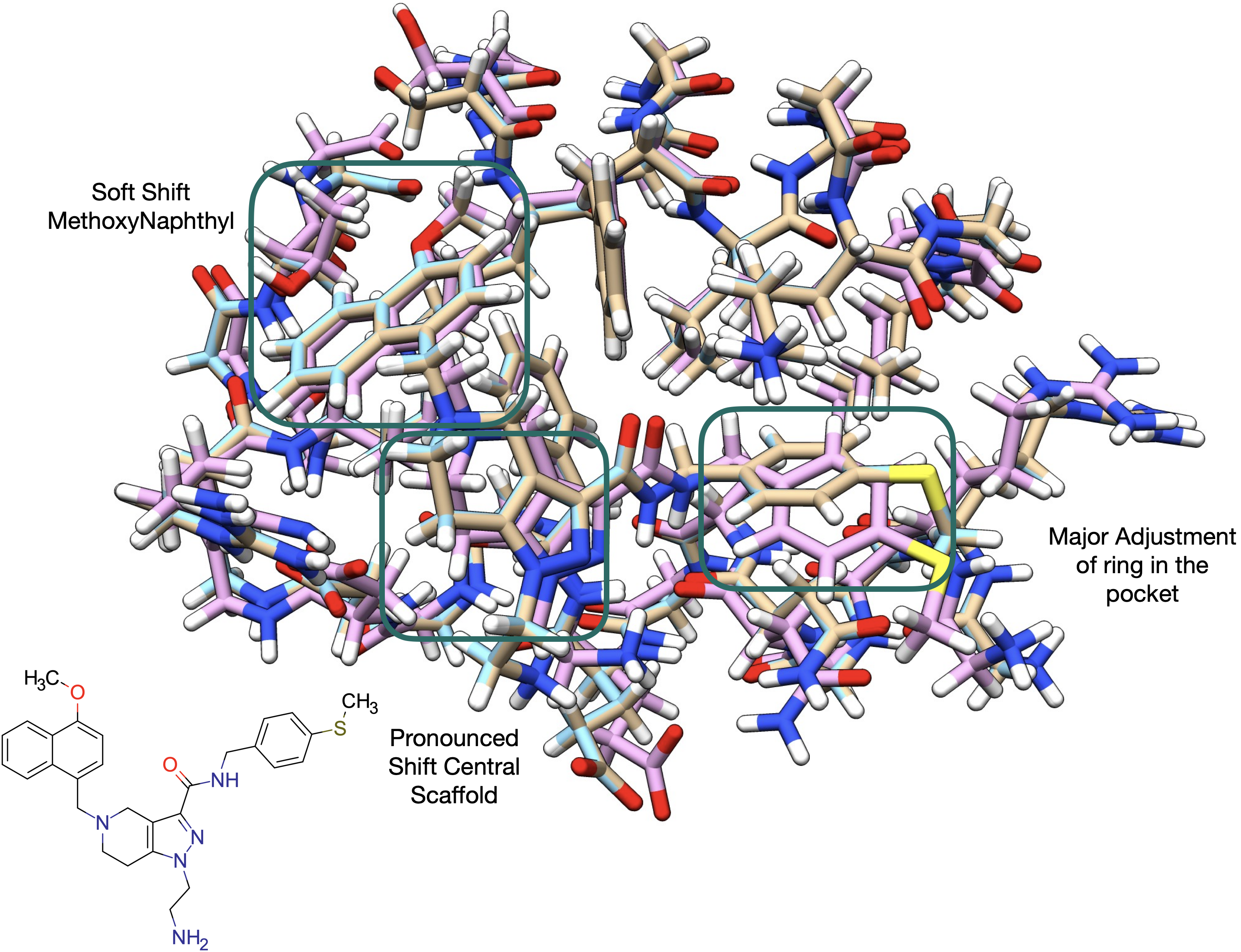

c

**Figure S10.2:** Comparison between bbPockets for a) 5L8A, b) 5N8V, and c) 8RIB. Brown represents the heteroatom coordinates from the crystal. Light blue stands for the in-pocket optimized structures, whereas pink represents the unconstrained optimized structures.

Structure refinement with In-Pocket systematically achieves minimal changes concerning the crystal structure. This contrasts with the observations for full geometry optimization, where RMSDs over non-hydrogen atoms are larger than 1 Å. In both cases, the removal of the backbone atoms of the most distal residues leads to a destabilization of the overall structures. This is because the structure capping leads to short contacts between hydrogen atoms, that must be refined through geometry optimization. Nevertheless, In-Pocket still retains the experimental crystal structure, though minimal adjustments are observed to minimize the clashes mentioned above. Geometry optimization leads to stronger changes in the structures. In the case of 5L8A, the methoxy naphthyl substituent of the ligand is slightly shifted. Stronger changes take place on the other aromatic anchoring point, the phenyl ring. Geometry optimization sits it deeper into the respective pocket. The central scaffold is in this case only slightly shifted when compared to the starting geometries. We see, however, that there are also significant changes in the overall locations and orientations of other functional groups. The RMSDs between initial and fully optimized 5N8V complexes show the least changes of all complexes studied. This is primarily motivated by the fact that this is the bulkier ligand, therefore keeping structural rearrangements like the ones described for 5L8V from occurring. Consequently, we may state that the binding pose is conserved in this case. Again, we observe the movement of side chains. This affects the whole structure. Lastly, we have the case of 8RIB. Though the methoxy naphthyl conserves its binding mode after geometry optimization, the central scaffold, and the other aromatic ring change significantly how they interact with the protein. This is most likely caused by the lack of structural waters from the geometry optimization. This stresses the observation that missing fine structural elements, no matter how fine they might be, might have catastrophic consequences for the structural refinement and the conclusions of the study. This motivates the need to work as closely as possible to the native, experimental structures, to mitigate problems from the underlying methods and models used for the calculations. We see therefore that even an RMSD change of 1.08 Å might carry significant structural modifications that might invalidate the model. Lastly, we observe that the least complete structural models (nobb Vs. bb) lead to stronger variations of the RMSD between the optimized and the initial structures.

RMSD (A)

|  | 5L8A | 5N8V | 8RIB |
| --- | --- | --- | --- |
| ip(bb) | 0.0331288 | 0.0314994 | 0.0295239 |
| ip(nobb) | 0.0496423 | 0.049694 | 0.049406 |
| opt(bb) | 1.26794 | 1.07654 | 1.34492 |
| opt_p(nobb) | 1.80034 | 1.38113 | 2.30066 |

Ebind (kcal/mol)

|  | 5L8A | 5N8V | 8RIB |
| --- | --- | --- | --- |
| ip(bb) | 26.6949 | 34.2531 | 28.1758 |
| ip(nobb) | 27.9886 | 36.4212 | 30.7592 |
| opt(bb) | 45.9397 | 56.9725 | 61.8141 |
| opt_p(nobb) | 48.104 | 62.0779 | 66.5218 |

**Figure S10.3:** RMSD (Å) and norm of the binding energy (kcal/mol) evaluation of the different optimization protocols on the different pocket cuts.

The structural changes that occurred during geometry optimization led to an improvement in the protein-ligand interactions. This is reflected in a decrease in all interaction energies. Interestingly, however, geometry optimization changes the relative order of binding energies: the value is lower for 8RIB, the complex for which more structural rearrangements took place during geometry optimization. It is then most likely that the relative order of binding energies is an artifact of the rearrangements that took place during refinement and has no biological significance. Lastly, we see that although geometry optimization should standardize structures, the complexity of protein-ligand systems prevents the optimization of a single binding energy for the same system.

Strategies to overcome these limitations are typically the zeroing of gradients for specific atoms. We see, however, that this might lead to unresolved closed contacts and clashes.

**S11 – Gas Phase Benchmark for Ligand-Residue Fragmentation and Interaction Analysis Excluding Waters**

The level of theory used in the calculations is reasonably low, so the calculations are fast. This also allows estimating interaction energies for many subsystems without requiring any special hardware. It is however important to evaluate the accuracy of the method employed, using higher-level calculations. In Figure S11.1 we benchmark GFN2-xTB gas phase binding energies for the same interactions reported in the main manuscript, Figure 5, against several dispersion-corrected density functional methods.

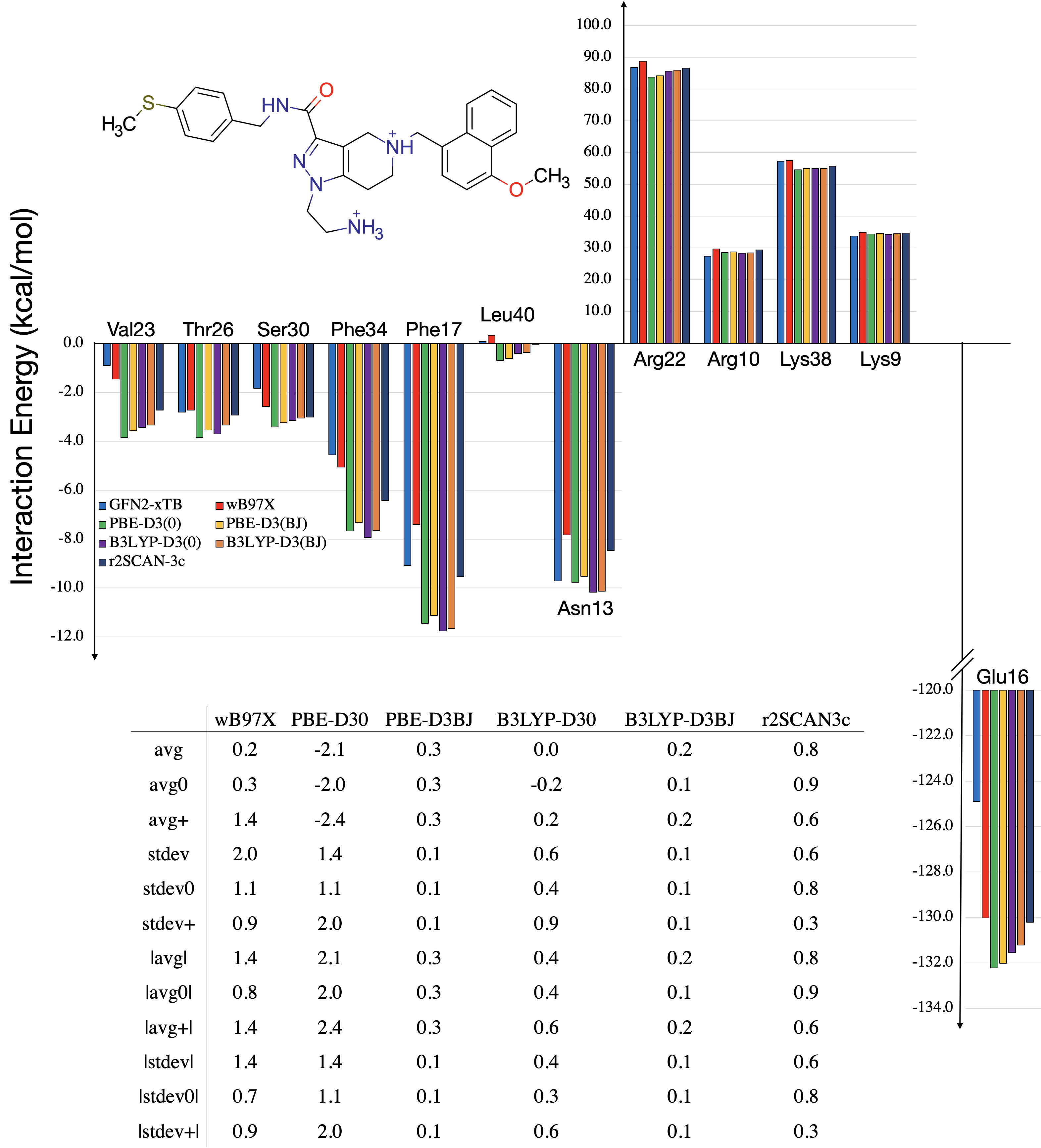

**Figure S11.1:** Benchmark of GFN2-xTB against higher-level dispersion-corrected DFT methods. Calculations in gas phase. DFT calculations using def2-TZVP basis set. In the table, avg stands for average deviation and stdev is the standard deviation. If the suffix 0 is used, then evaluation is performed on neutral residues, whereas the + is reserved for positively charged residues. Bars around the parameter indicate absolute quantities. Therefore, |avg0| is the absolute average deviation taken over neutral residues.

The results clearly show that GFN2-xTB, for the biological systems evaluated, behaves on par with the computationally more demanding DFT methods. Larger discrepancies are observed in the comparison with Glu16. We stress however that the basis set used in the DFT calculations does not include diffuse functions, meaning that the DFT calculations might be in error.

In the section “***A complex of PEX14 with compound 1: analysis of the structure***” of the main manuscript several interactions were identified based on the geometric analysis of the crystal structure. Their relevance for binding could not be quantified, so the analysis remained purely qualitative. Here, IPOPT is used to analyze the relative strength and relevance of different contributions. Figure S10.2a shows interaction energies for the ligand and those residues, whereas Figure S11.2b shows interaction energies at the ligand-functional group level. Note that here we perform an analysis of all residues of the pocket excluding water molecules seen in the crystal. In the main manuscript, the analysis is performed for a few key residues and the impact of water is also explored.

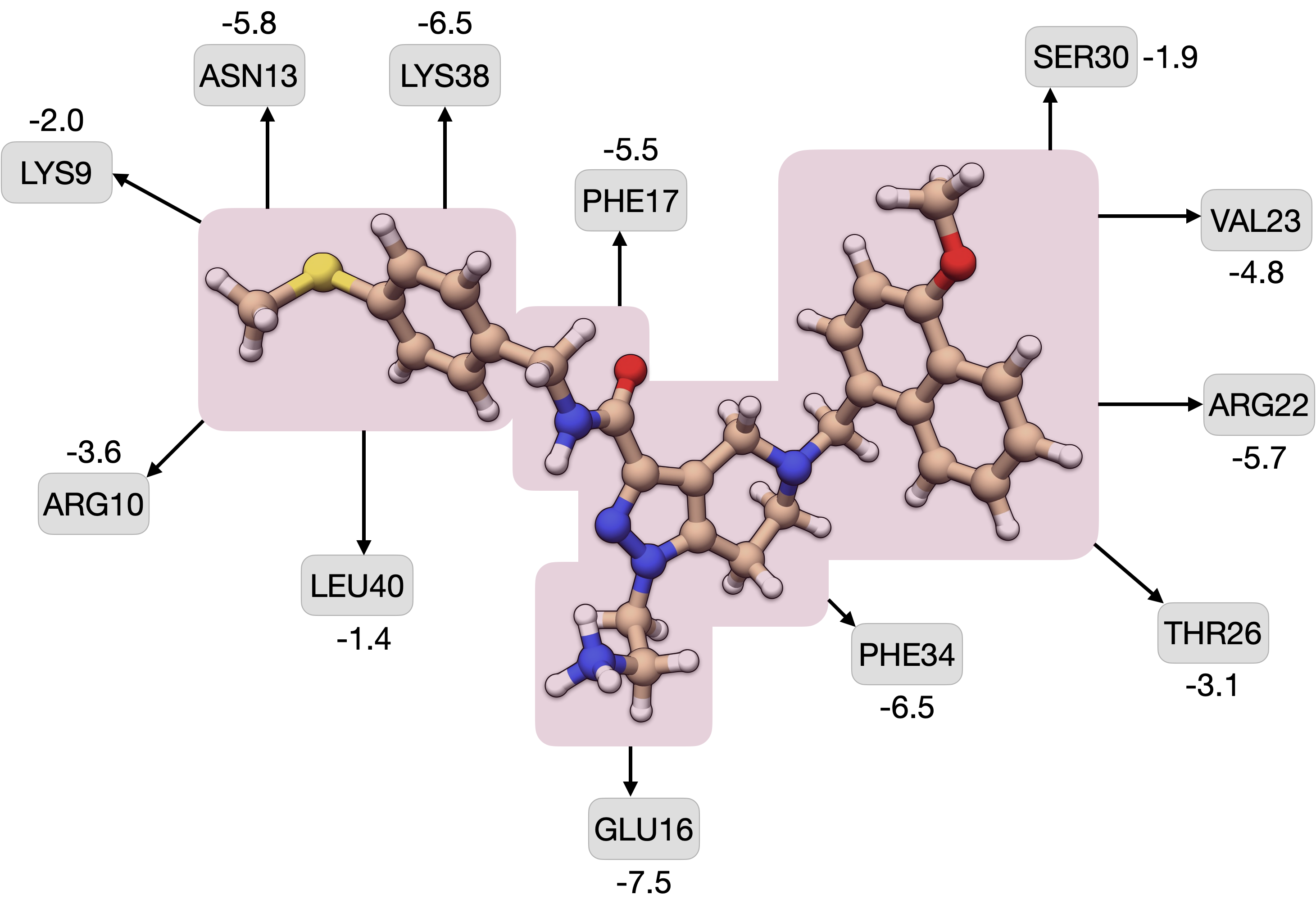

a

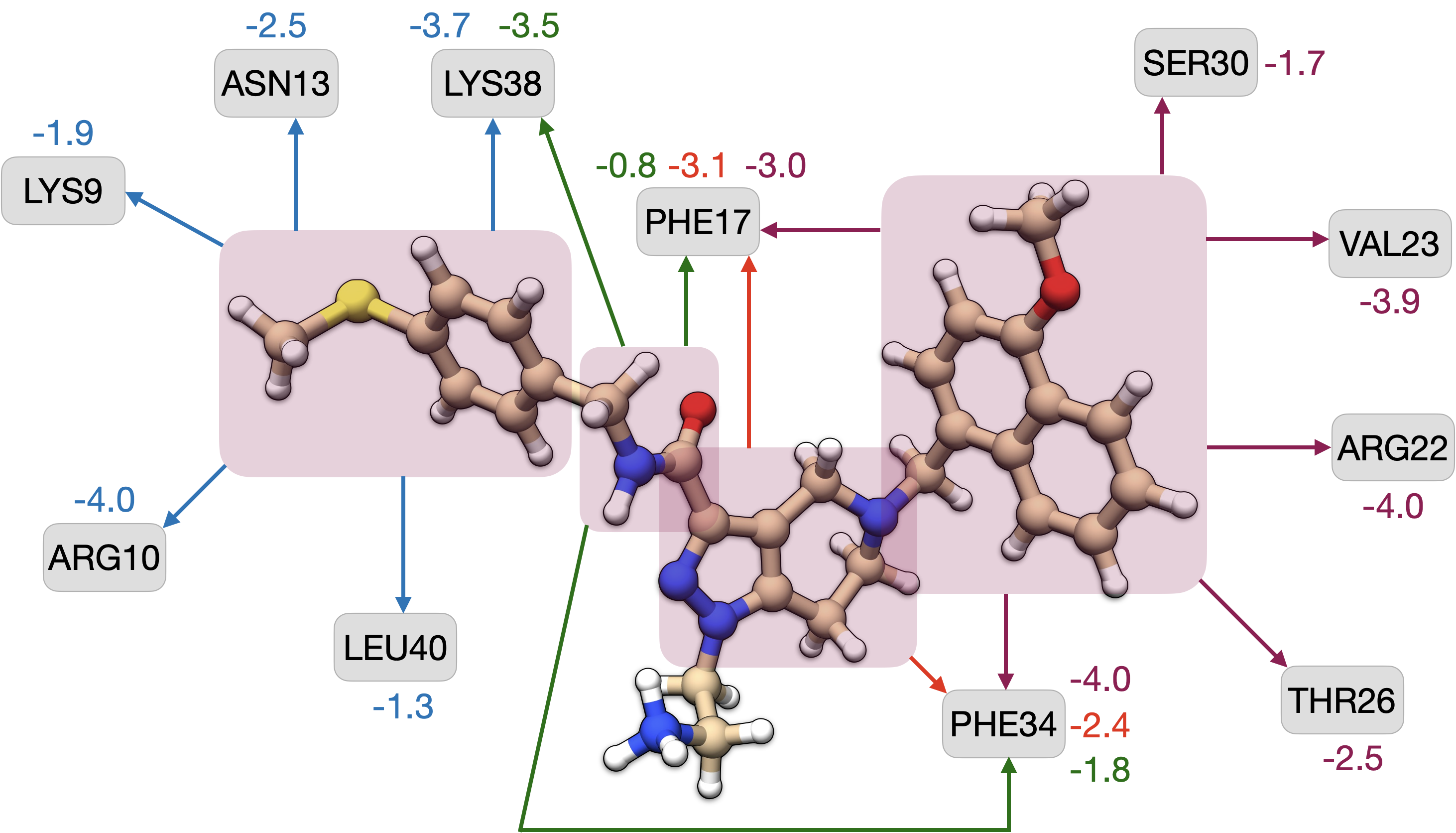

b

**Figure S11.2.** a) Quantification of the interaction energy between the ligand and specific residues previously identified from structural analysis. b) Quantification of interaction energies between residues and specific functional groups from the ligand. All energies in kcal/mol.

Of the protein-ligand contacts previously identified, the interaction between Leu40 and the aromatic sulfide is of lesser relevance. Calculations using the whole ligand or just the smaller fragment yield similar binding energies, which is in good agreement with the type of interaction analyzed. The stabilization achieved with Ser30 is also marginal, which is justified by the fact that the ligand interacts with the main chain carboxyl group. Further, this contact is established by the methyl group, which, due to its flexibility, leads to statistically vanishing attractive contributions, i.e., the spontaneity of the interaction might vanish when conformational entropies are considered. An interesting observation from Figure 3 is the contribution of the amide linker, which seems to be quite an active element for binding. This is particularly critical in the case of the interaction with Lys38, as the sulfide group and the amide linker contribute to equally strong interactions. To better understand the nature of interactions between Lys38 and the ligand, i.e., whether the stabilization of -6.5 kcal/mol results primarily from dispersion or whether this is electrostatic, in-pocket was used to replace the lysin’s ammonium with a methyl group. The interaction energy lowered to -4.5 kcal/mol, which indicates a predominantly dispersion-like interaction. We note that dispersion energies correlate with lipophilicity. Interactions with Phe34 are also of interest to analyze. Though there is a clear predominance of the T-stack with the methoxy-naphthyl group, interactions with the carboxamide linker and the main scaffold are of similar magnitude. The interaction between the primary ammonium and Glu16 is however the strongest we sampled. This corresponds to an Hbond with a high ionic character, which justifies its stability.

Overall, the main interaction points between protein and ligand are Phe34, Arg22, Val23, Phe17, Lys38, Asn13, and Glu16. It is interesting to note that, though the T-stack with Phe34 is considered the predominant interaction with the methoxy-naphthyl moiety, the nearby Arg and Val residues have also a particular relevance for defining the interaction model. Note that some of these interactions are potentiated by the presence of water. A discussion on the impact of the water envelope on binding comes in the main manuscript.

The primary concept behind the exclusion of specific waters from the crystal structure was the region of the protein-ligand complex they interacted with. This allowed us to identify two main groups, one bridging the ligand’s carboxamide with Ser37, and the other bridging the amide and tertiary ammonium of the ligand. Additional exclusions were strictly based on proximity and requirement for unbroken/uninterrupted interaction. For instance, the trail of 4 waters connecting the ligand’s amide with Ser37 was used in investigating the importance of these waters for the specific interaction with Ser37. Note that other constructs would be feasible, however, for understanding this specific interaction, only 4 or 0 waters are providing meaningful insight. Further, one of these waters is involved in contact with Lys38, and therefore, it too was investigated for its impact on the ligand-Lys38 interaction energy. The remaining 3 water molecules could have been included too. However, these waters would not have a direct participation. In the case of Phe34, there is no direct interaction with water molecules or any water-mediated contact, therefore calculations were always performed without water. A similar logic was used in the case of the other trail of water molecules.

**S12 – Optimized Structure for Compounds 1 and 14 with One Explicit Water**

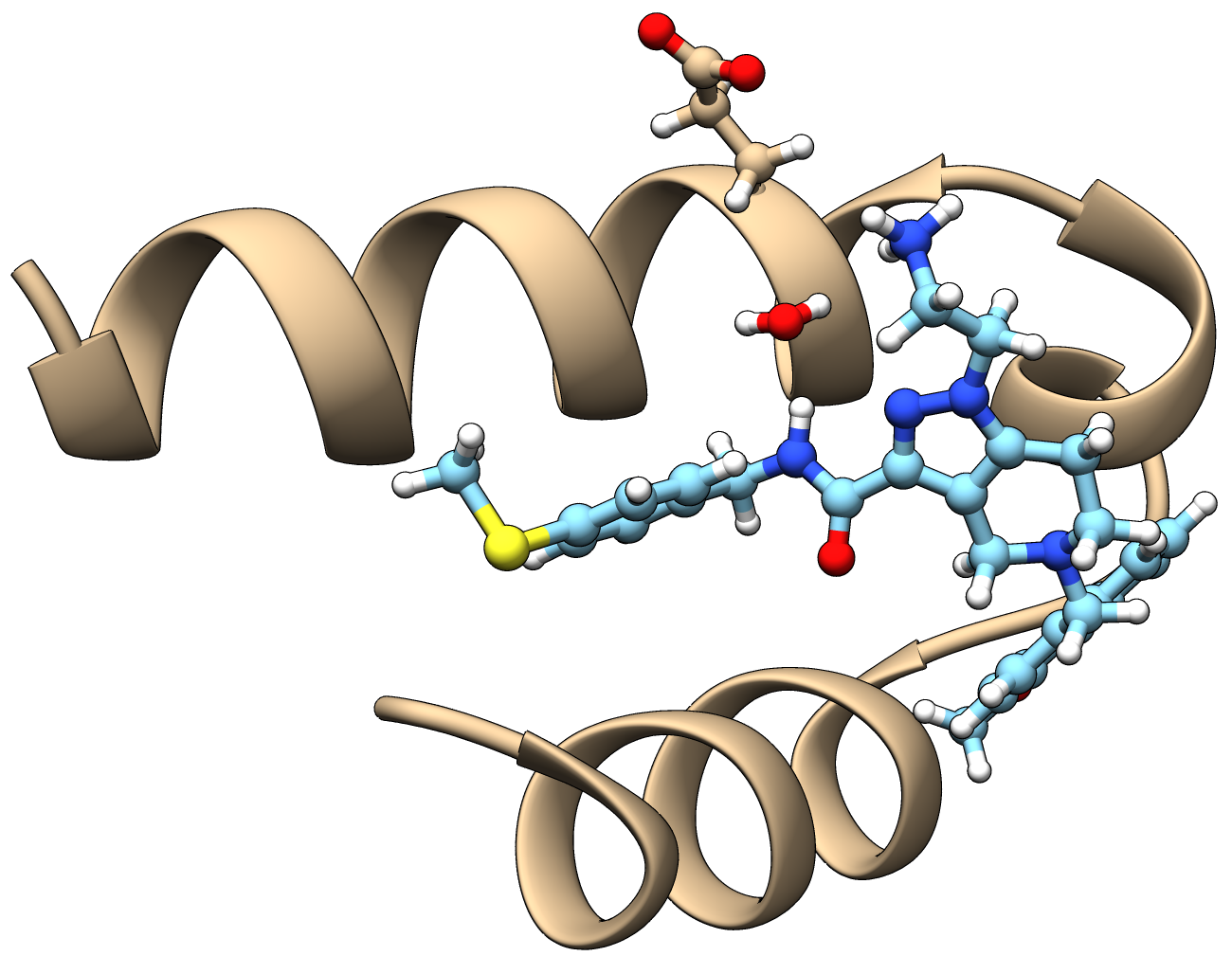

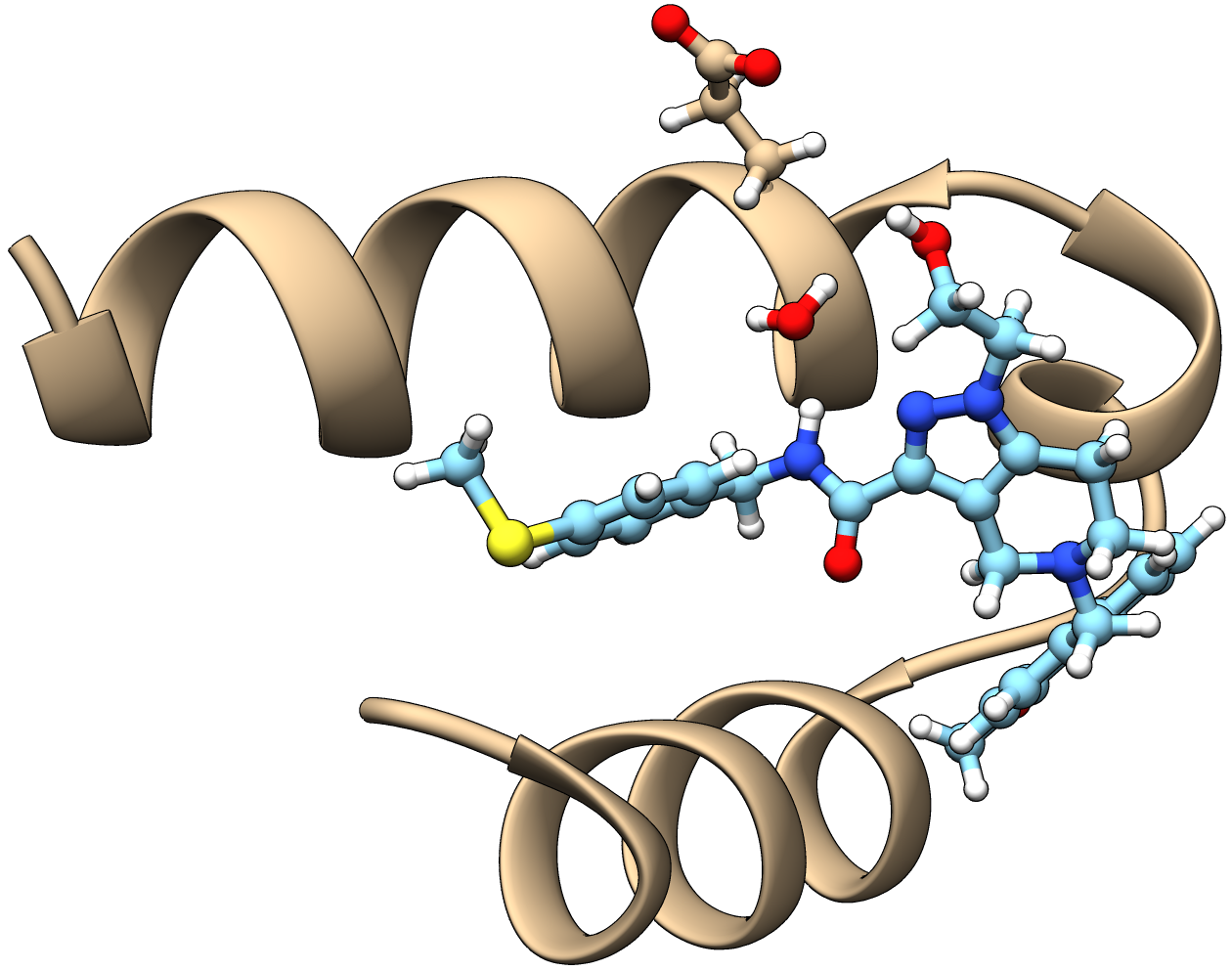

**Figure S12.1:** Structures of protein-ligand complexes with one explicit molecule of water from the crystal. On the left, the compound **1** that was recently crystallized. The molecule of water orients itself to become an H-bond acceptor. On the right, compound **14**. Water orients itself, to become an H-bond donor.

**S13 – Data Collection and Refinement Statistics**

**Table S13.1:** Data collection and refinement statistics. Statistics for the highest shell are listed in parentheses.

| PDB ID | **8RIB** |
| --- | --- |
| Inhibitor | **1** |
| Data collection |  |
| Space group | C 1 2 1 |
| Cell constants:  a, b, c (Å)  α, β, γ (°) | 56.01, 45.74, 52.62  90.00, 118.87, 90.00 |
| Wavelength (Å) | 1.000 |
| B factor (Wilson) (Å 2) | 14.48 |
| Resolution range (Å) (highest shell) | 46.08 – 1.13 (3.41-1.13) |
| Completeness (%) | 89.6 (36.2) |
| Rmerge (%) | 5.1 (54.9) |
| Rmeas (%) | 2.2 (35.1) |
| Observed reflections | 191976(4943) |
| Unique reflections | 35511(1642) |
| I/σ(I) | 14.7 (1.7) |
| CC(1/2) (%) | 99.9 (71.3) |
| Redundancy | 5.8 (3.0) |
| Refinement |  |
| Resolution (Å) | 46.08 – 1.13 |
| Number of reflections used | 35511 |
| R-factor (%) | 12.6 |
| Rfree (%) | 16.2 |
| Average B (Å 2)  Protein  Ligand  Water | 19.5  29.4  33.3 |
| RMS from ideal values  Bond length (Å)  Bond angles (°) | 0.013  1.83 |
| Ramachandran statistics (%)  Most favored regions  Additionally allowed regions  Generously allowed regions | 98.3  1.7  0.0 |
| Content of asymmetric unit |  |
| Number of protein molecules/residues/atoms  Number of ligand molecules/atoms  Number of solvent molecules | 2/67/1046  2/128  173 |

**S14 – Experimental SAR Data.**

The experimental data used to assess the effect of lactamization is presented below. EC_50_s for different Trypanosoma parasite strains are given in μM.

**Table S14.1:** Data assessing the impact of lactamization.

| Compound | 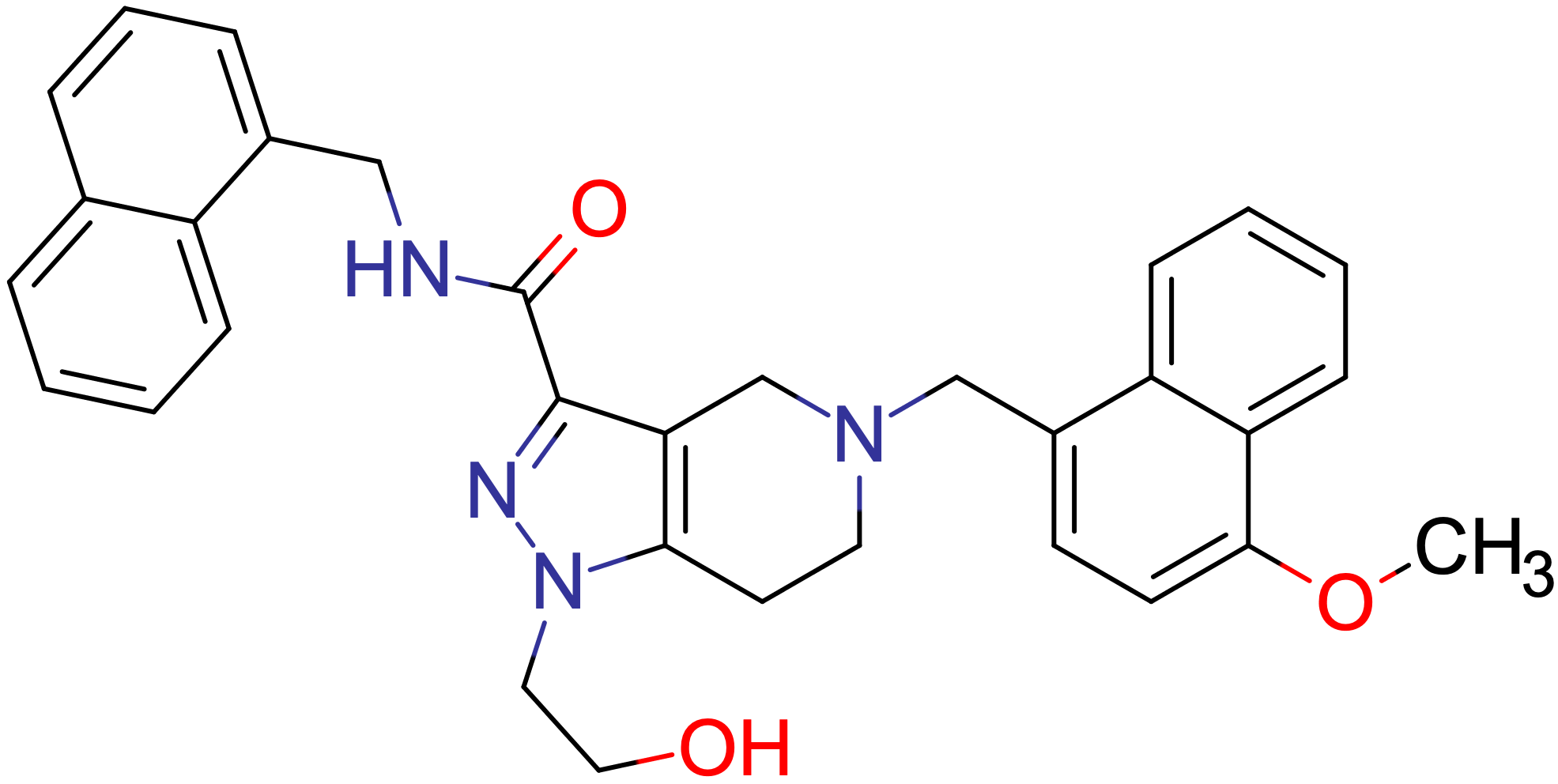 | 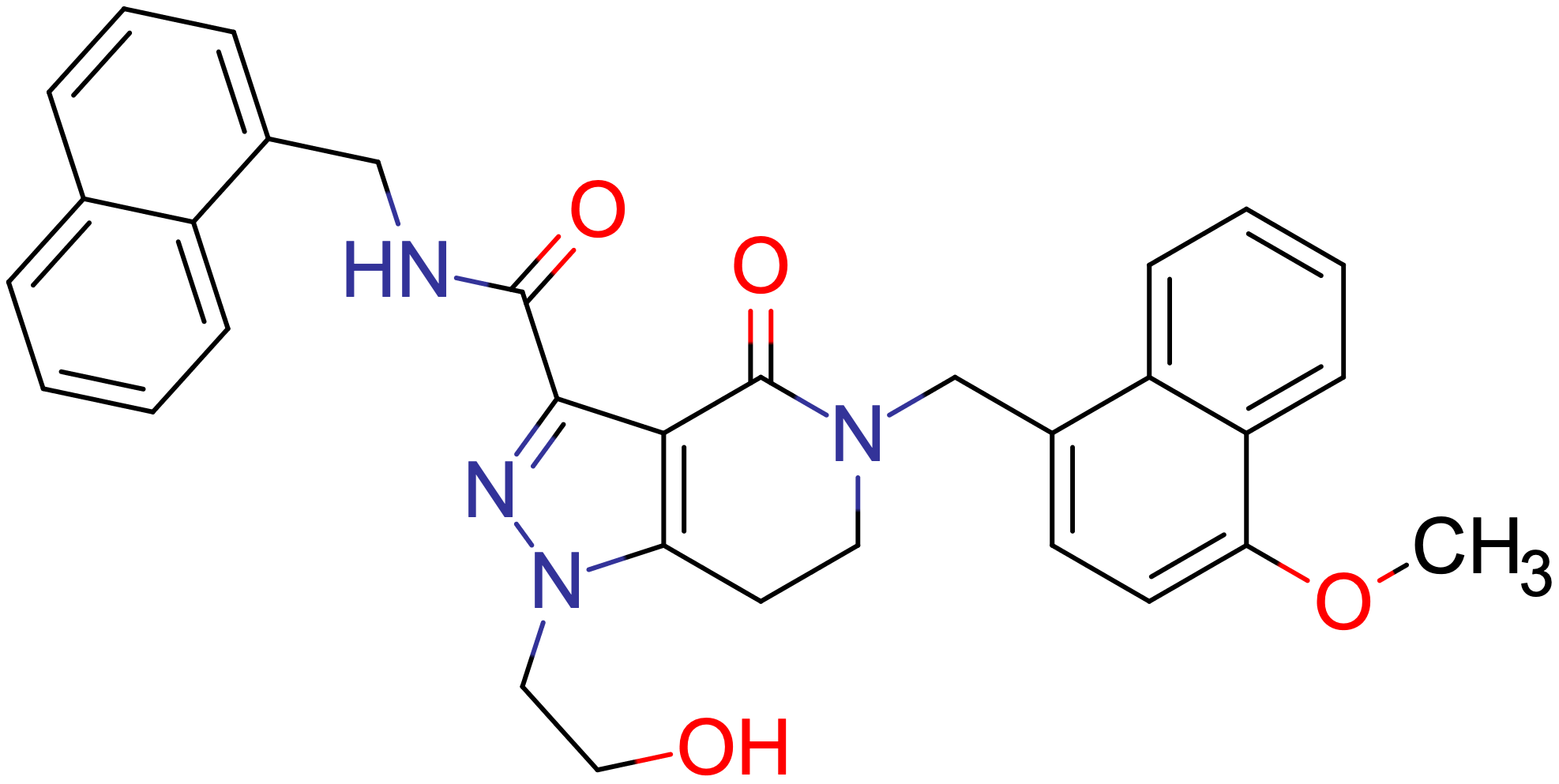 |
| --- | --- | --- |
| EC_50_ (*T. brucei*)  [cellular assay] | 3.56 | 5.26 |
| EC_50_ (*T. cruzi*)  [cellular assay] | 3.83 | 24.1 |
| EC_50_ (*T. brucei*) [AlphaScreen] | 14.8 | — |
| EC_50_ (*T. cruzi*) [AlphaScreen] | 104 | — |

The data for the flipped amide is presented in the table below.

**Table S14.2:** Data assessing the impact of flipping the amide.

| Compound | 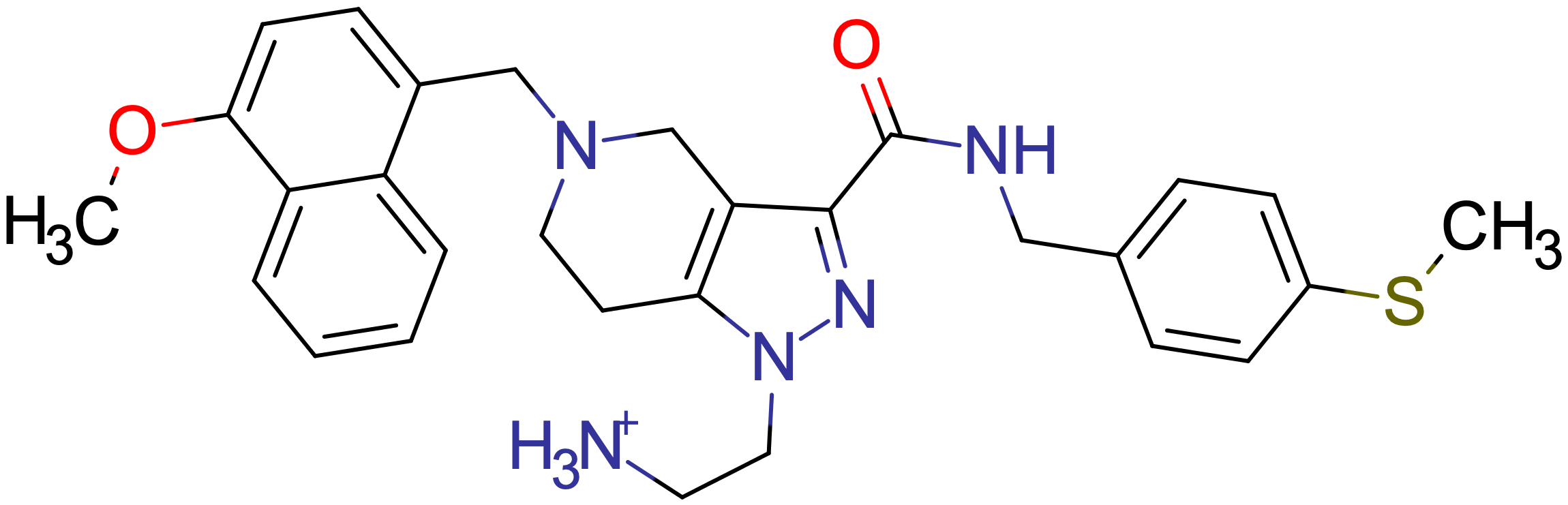 | 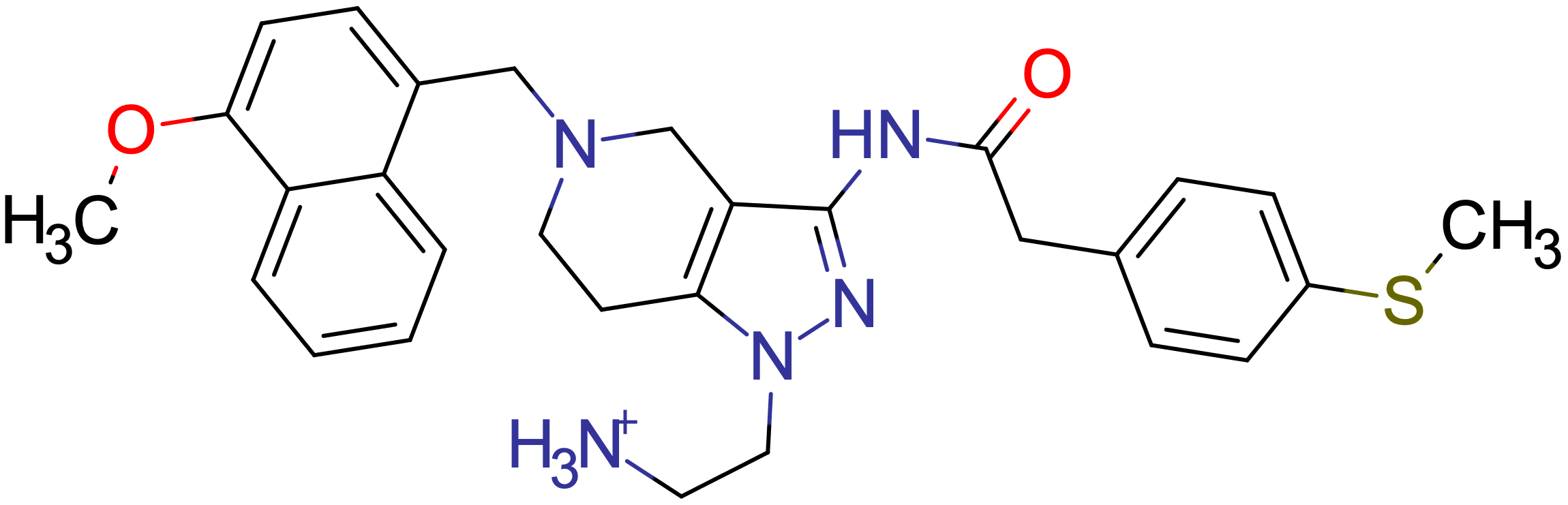 |
| --- | --- | --- |
| Name | **1** | **1**f |
| EC_50_ (*T. brucei*)  [cellular assay] | 0.122 | 0.324 |
| EC_50_ (*T. cruzi*)  [cellular assay] | 6.64 | 9.84 |
| EC_50_ (*T. brucei*) [AlphaScreen] | 27.4 | 32.8 |
| EC_50_ (*T. cruzi*) [AlphaScreen] | 37.0 | 53.4 |

The data for the alcohol is presented in the table below.

**Table S14.3:** Data assessing the binding of compound **14**.

| Compound | 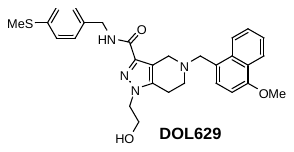 |
| --- | --- |
| Name | **14** |
| EC_50_ (*T. brucei*)  [cellular assay] | 6.96 |
| EC_50_ (*T. cruzi*)  [cellular assay] | — |
| EC_50_ (*T. brucei*) [AlphaScreen] | 80.0 |
| EC_50_ (*T. cruzi*) [AlphaScreen] | 89.0 |

The binding energies of compounds **1** and **1f** docked to PEX14 are (in kcal/mol)

| Compound | **1** | **1**f |
| --- | --- | --- |
| Complex A | -27.02 | -26.41 |
| Complex B | -23.40 | -24.44 |

Using a Boltzmann average of the binding energies, we obtain the average binding energy of -27.02 kcal/mol for compound **1** and -26.35 kcal/mol for compound **1**f. The difference in binding between the two species is -0.67 kcal/mol. This can be converted to relative binding energies of

$$\frac{K_{d}^{1f}}{K_{d}^{1}}=3.1$$

The experimental value is 2.7. The difference we assign to lack of entropy effects and vibrational contributions. It is also interesting to convert the experimental affinities to relative Gibbs free energies of binding. From the experimental data, we get the value of -0.58 kcal/mol (compared to -0.67).

The binding poses for compounds **1** and **1**f obtained from docking are presented below.

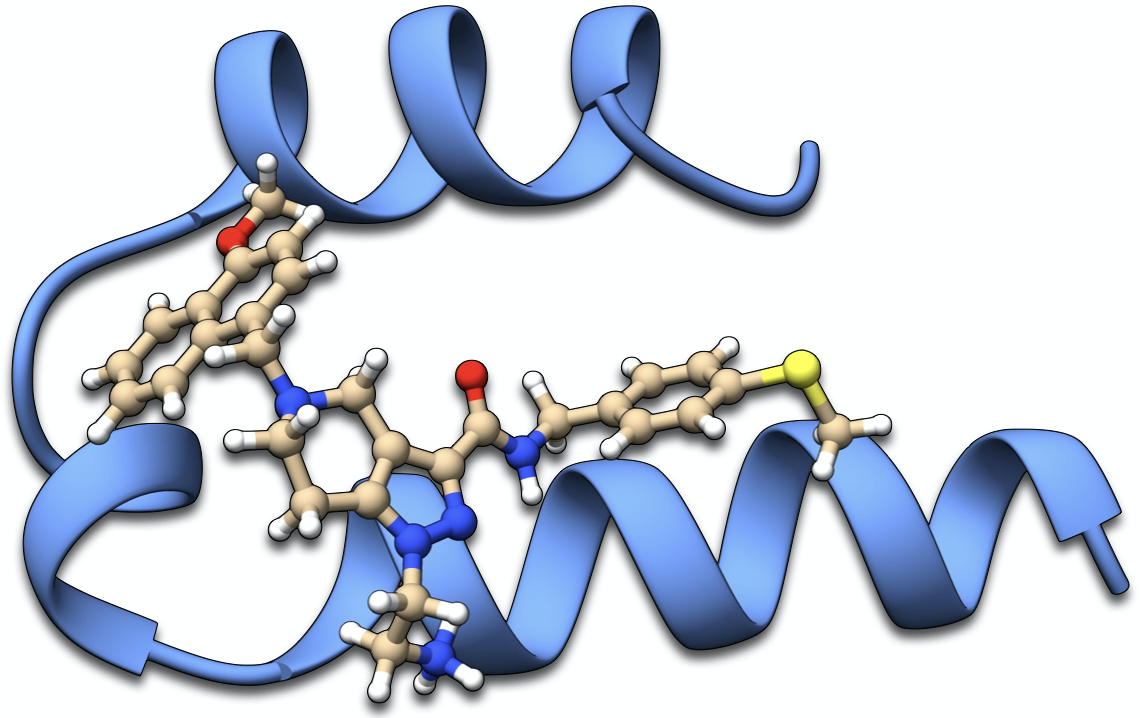

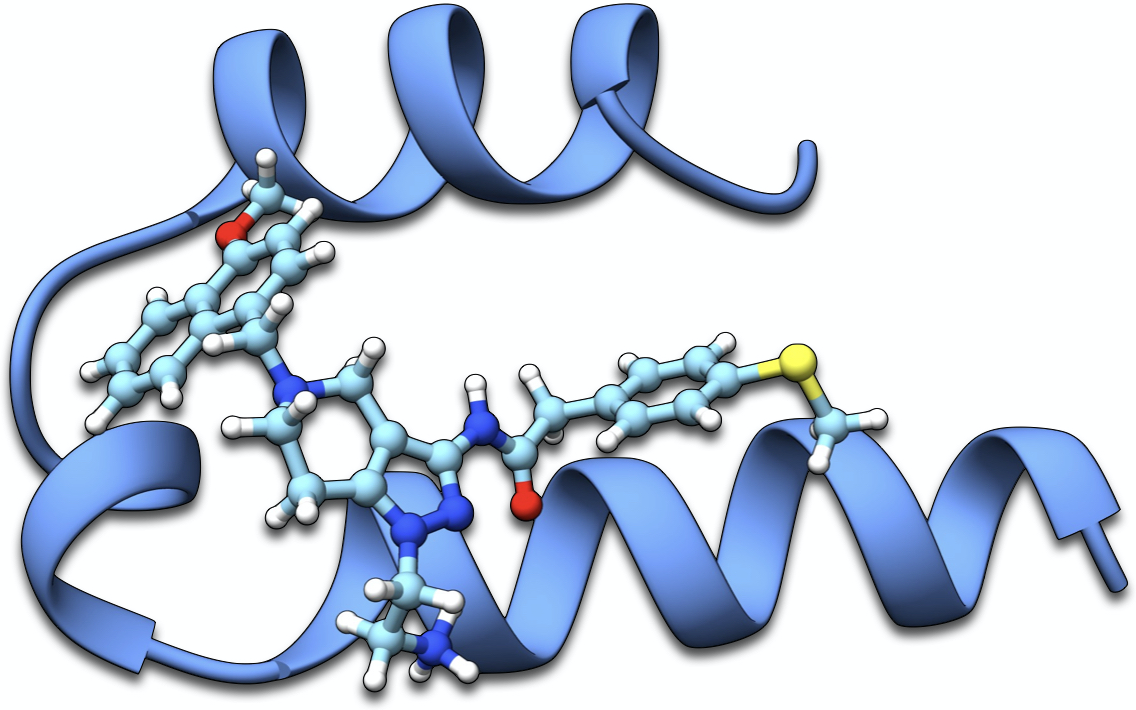

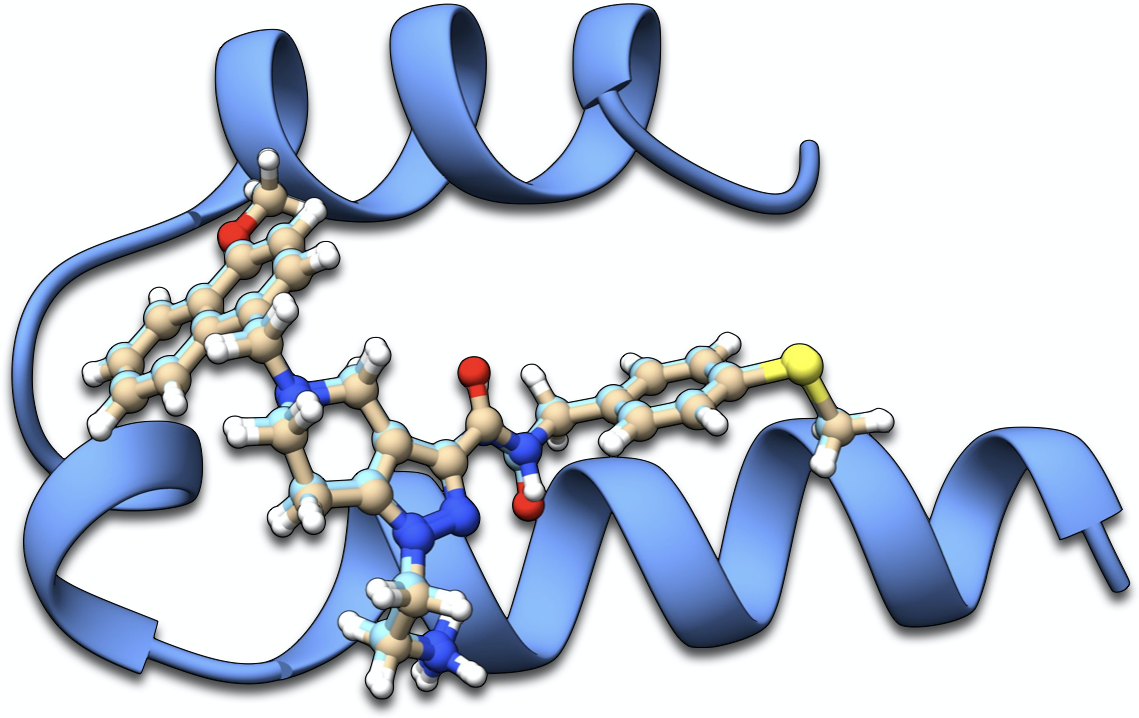

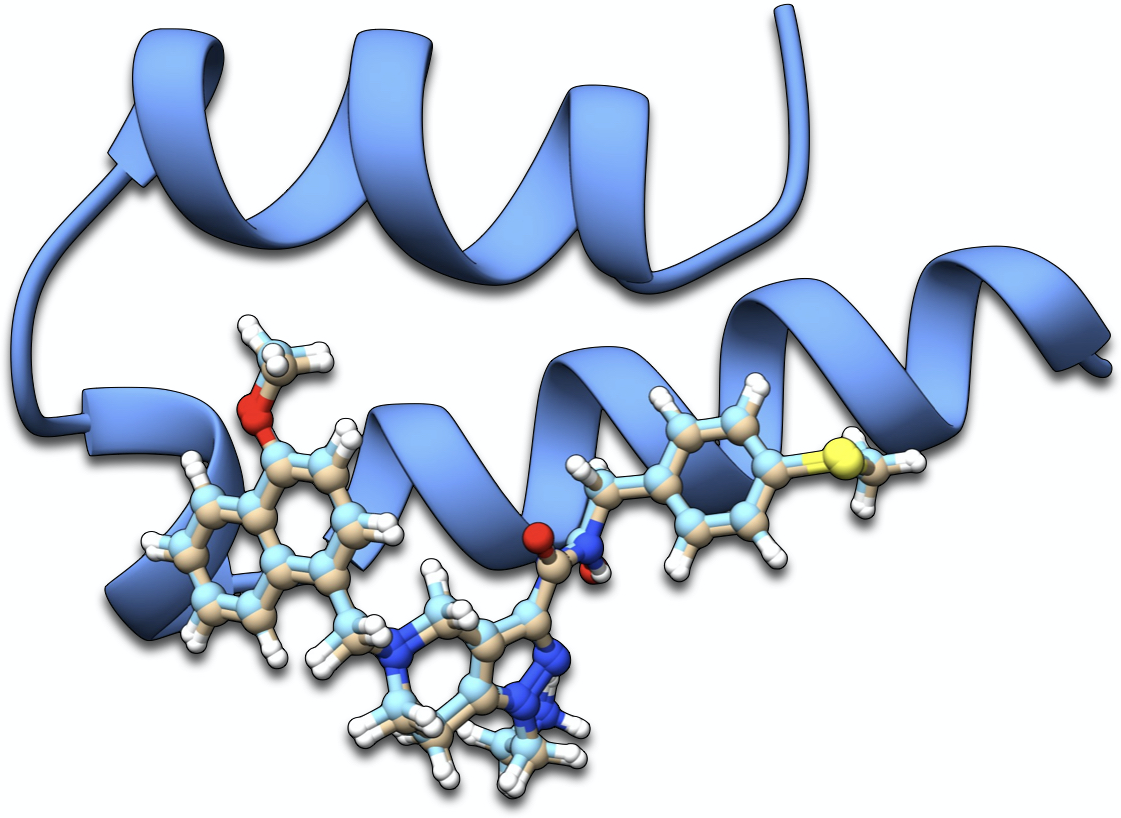

**Figure S14.1:** Binding pose of compound **1** after docking. Then the binding pose of **1**f after docking. Finally, the comparison of both binding poses.

**S15 – Further Analysis of in silico SAR data**

An alternative approach to reduce the tension on the sulfur atom is by hydrogenation of the respective phenyl ring (compound **4** of Table 1, main manuscript). This is attested by the calculations. However, removing the aromatic character impairs binding by 4.5 kcal/mol. If we further consider the absence of tension, then the impact scales to about 9 kcal/mol. We conclude that this is not a favored structural modification to improve binding. Though the binding pose we obtained with in-pocket is certainly still biased to the volume the aromatic ring requires and cannot properly accommodate the expansion necessary for the saturated ring, removal of aromaticity should severely impact the stacking with Asn13. Besides this interaction, this fragment of the ligand contributes with two additional strong contacts, one with Lys38, and the other with Arg10, both dominated by interactions with the methylene groups of the side chains.

To further explore the possibilities opened by our algorithm, we attempted more daring variations of compound **1**. In an *in-silico* environment, these modifications lead to a better understanding of the nature of protein-ligand interactions. In compound **6** (Figure S14.1b), the sulfur atom was oxidized to sulfoxide, a typical metabolic modification. The potential benefit is the promotion of interactions with two nearby Lys residues via electrostatics and hydrogen bonding. The binding for the new structure indeed improves by 3.7 kcal/mol. To determine whether the charge effect is more general, we removed the methyl group from the sulfide and transformed it into a phenyl thiolate (**8**, see also Figure S14.1a). The net effect was quite disastrous, as the binding energy increased by 7.9 kcal/mol. A closer analysis of the respective structures reveals that the sulfoxide → thiolate conversion brings the formal negative charge closer to the oxygens of carboxyl groups and further from the lysins.

Generating a pyrrole from the pyrazole (**5, 13**) improves binding even further. Including a molecule of water, the very same one that proved essential to distinguish the ammonium and hydroxyl variants of the ligand, shows however that binding is disfavored by this modification: the binding energy of **5** increases by 0.4 kcal/mol.

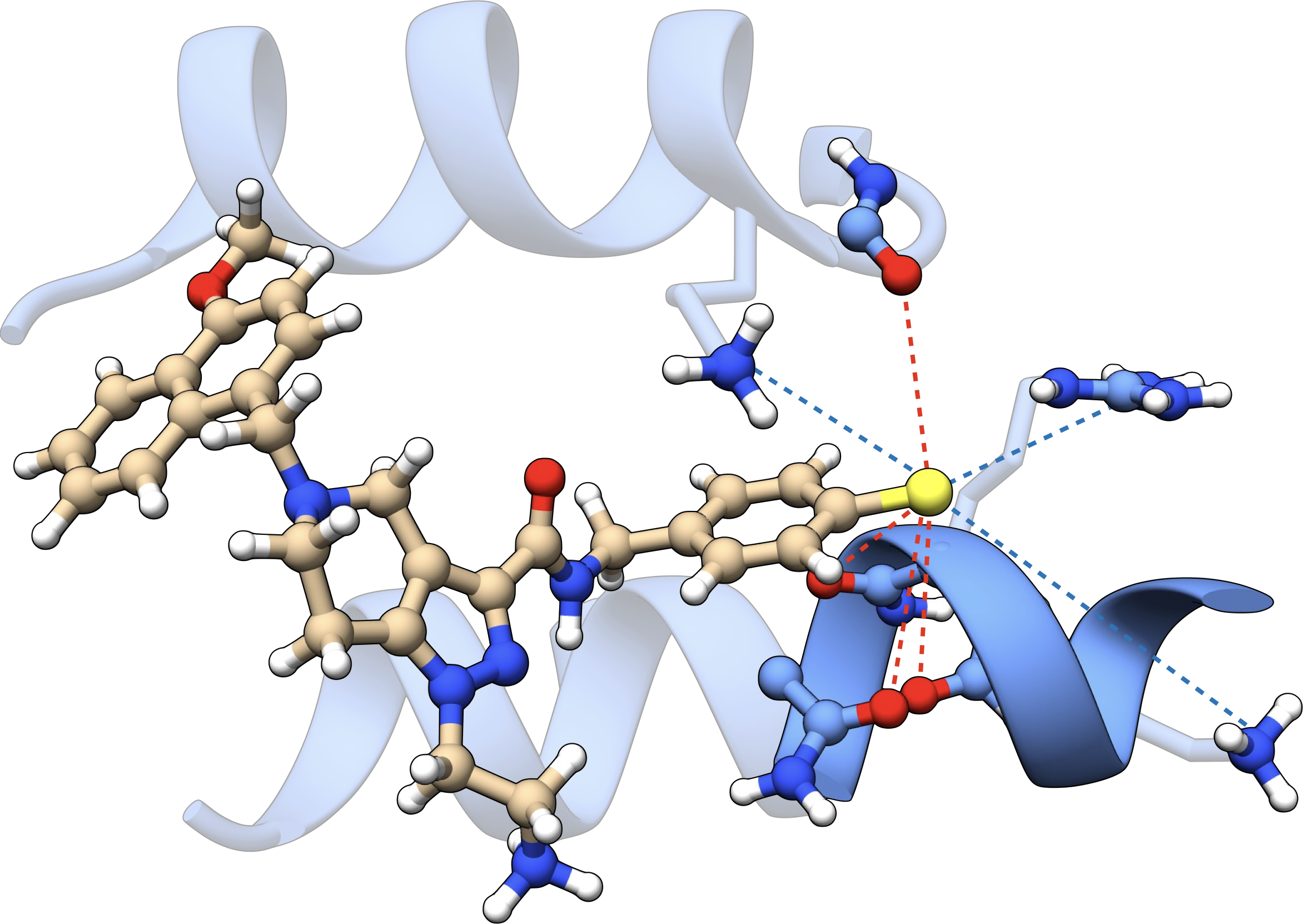

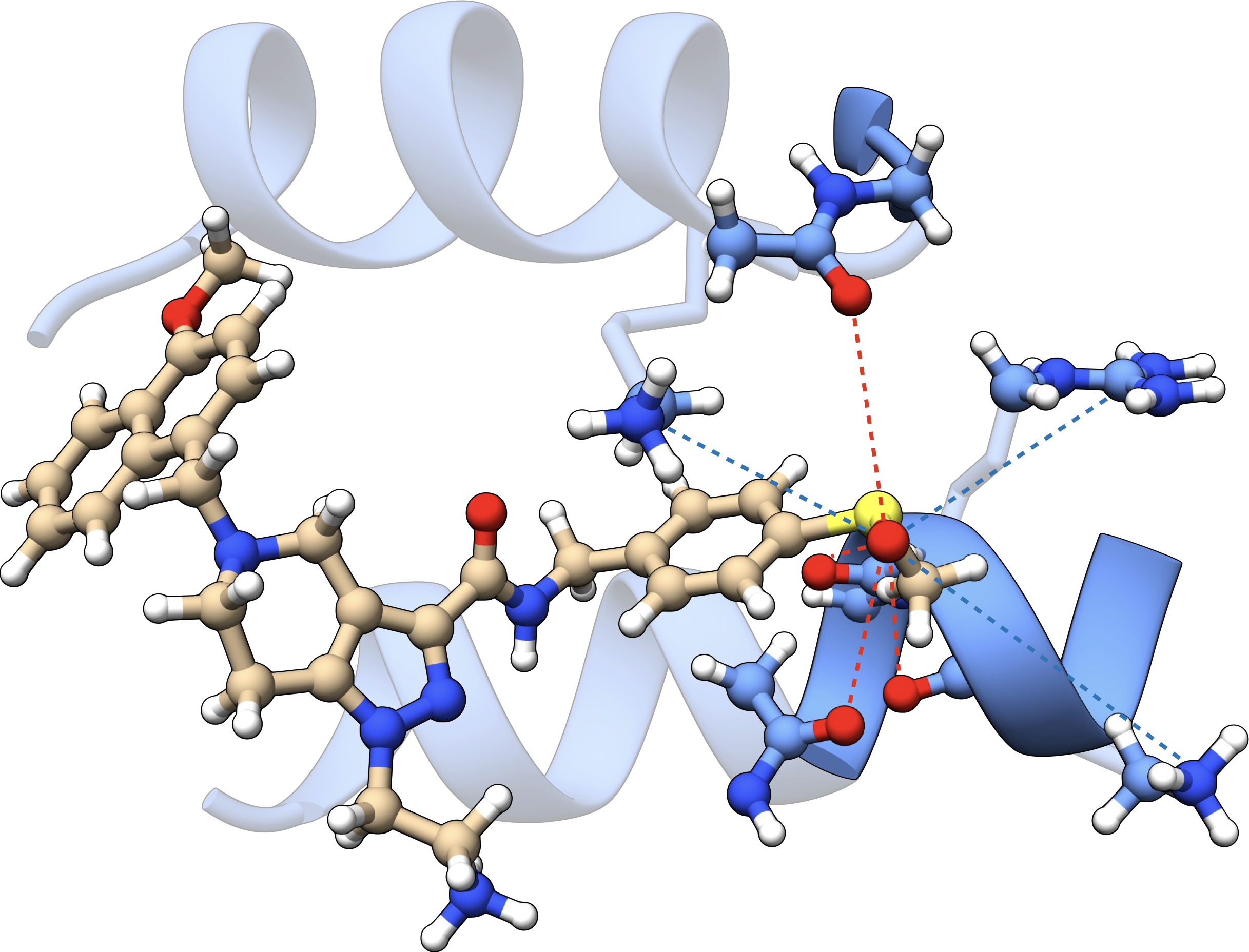

b

a

**Figure S15.1:** In-pocket optimized binding poses for compounds a) **8** and b) **6**. In the subfigures, the protein’s backbone scaffold is shown as blue ribbons. H-Bond networks are marked with green dashed lines. Attractive electrostatic interactions are marked with blue dashed lines whereas repulsive electrostatics are marked with red dashed lines.

**S16 – Further Analysis of Semiempirical docking data**

Introducing a carboxyl group in the tetrahydropyridine-pyrazole scaffold (**12**) to form a lactam also seems to be beneficial for binding. Again, we expect Hbonds and flattening of the six-membered ring to be key. Structure-Activity Relationship (SAR) studies conducted in our groups revealed however that the amine → lactam transformation is not beneficial, with the affinities increasing by 0.2 kcal/mol (AlphaScreen data for one such modification available in S13). Since the different behavior is hardly caused by the substituents on R1, we decided to extend our studies to include an explicit molecule of water. To have the best possible structures for the ligands in the protein’s pockets, we docked them using in-pocket optimization in the presence of one explicit water. The docking was performed with unconstrained protons. The resulting interaction energy of the lactam was 3.2 kcal/mol higher than the respective amine. The problem however was that the amide group was not entirely flat. Further flattening the latter leads to a deviation of 1.0 kcal/mol on the binding energy (with respect to the amine), in much better agreement with the experimental data. We then conclude that the replacement of amine with lactam is not beneficial due to the disruption of a Hbond.

**S17 – Methodology used for evaluating Binding Energies and Affinities**

Figure S17.1 shows the schematic for the methodology used to calculate affinities. Note that the exact calculation of affinities would require an exhaustive calculation of all terms indicated in Figure 16.1. In the case of energies ($\Delta E_{strain}$, $\Delta E_{bind}$, $\Delta E_{hydr}$, and ${\Delta E}_{bind}^{H2O}$) and enthalpies ($\Delta H_{TRV}, {\Delta H}_{TRV}^{H2O}$) the model employed requires averaging over several conformational states.^58^ In the case of entropies, additional terms arise to account for the stabilization provided by having a conformer ensemble for each molecule, the so-called conformational entropy.^58^ The main limitation of our calculations to the full protocol in Figure S17.1 is therefore the neglecting of conformational sampling for the protein, ligand, protein-ligand complexes, and solvent molecules. For conceptually correct affinities, considerations must be made on the presence of several protonation states for each of the molecular aggregates. Both these points are mitigated by using for calculations a representative protein-ligand complex obtained from the crystal. This is particularly suitable for the estimation of enthalpies, as these are simple average properties and contain no additional terms, unlike entropy. Additionally, entropies and corrections to transform binding energies into enthalpies of binding used smaller protein-ligand constructs, which neglect the impact of distal residues for vibrational corrections. Our goal, however, is not the accurate calculation of absolute affinities, but to have a complete enough model that accurately reproduces trends, *i.e.*, to calculate relative affinities for QSAR. Lastly, implicit solvation is always employed in the calculations to mitigate the limitations of small hydration spheres and to account for the effect of molecules in the bulk of a solution at infinite dilution.

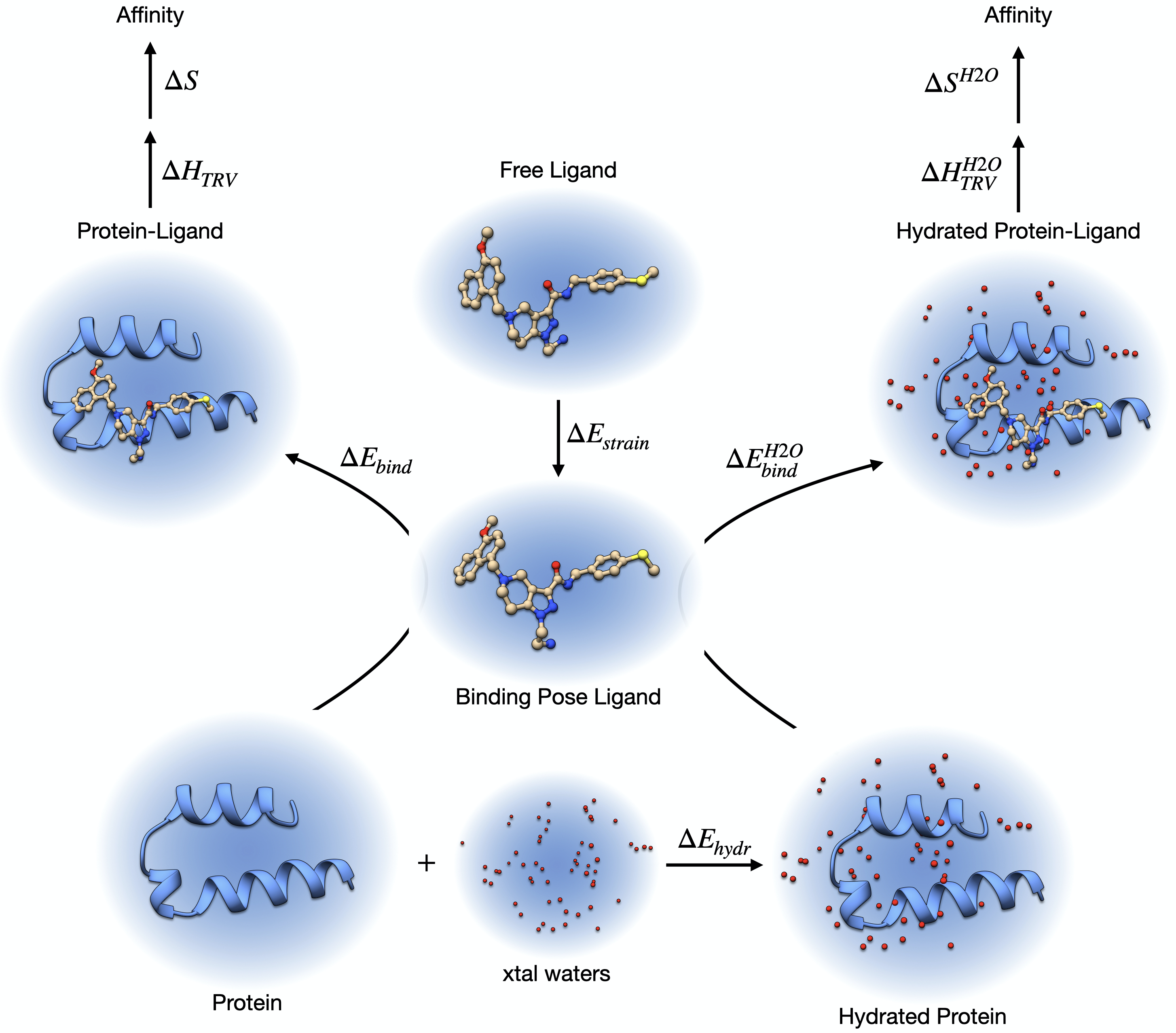

$$Affinity=\Delta G=\Delta E_{strain}+\Delta E_{bind}^{X}+\Delta H_{TRV}^{X}+\Delta S^{X}+\Delta\delta G_{solv}^{X}$$

$$\Delta S^{X}=\Delta S_{TRV}^{X}+\Delta S_{conf}^{X}$$

**Figure S17.1** Schematic representation of the methodology used to calculate affinities. Blue shaded areas represent the implicit solvation model. The protein hydration energy ($\Delta E_{hydr}$) is obtained from the free protein and the crystallization waters after in-pocket optimization. The ligand strain ($\Delta E_{strain}$) is approximated from the in-pocket optimized ligand against the closest unconstrained ligand. Binding energies ($\Delta E_{bind}, {\Delta E}_{bind}^{H2O}$) are obtained from the free ligand and the protein(-water) system. This can be in-pocket or fully optimized. To transform binding energies into affinities, two additional quantities were calculated: translation-rotation-vibration contributions to enthalpy ($\Delta H_{TRV}, {\Delta H}_{TRV}^{H2O}$), and entropy ($\Delta S, {\Delta S}^{H2O}$). The latter should ideally contain sampling contributions from protein, ligand, complex, and solvent. The equations at the bottom show how affinities are calculated according to the terms in the figure. The superscript X refers to how waters are being included in the treatment. If these are explicitly included, then the additional hydration term is included. Otherwise, only implicit solvation is employed ($\Delta\delta G_{solv}^{X}$).

Given this thermodynamic model for binding affinity, comments on the reasonability of our results in *in-silico* SAR are due (all sections in the main manuscript, except the one on the water envelope). The validity of our calculations strongly depends on several assumptions. The first and by far the most important one is that the binding pose remains unaffected by the structural modifications investigated. This may however be remedied using in-pocket to redock the compounds or by using any external docking software, potentially followed by in-pocket refinement. Furthermore, we assume the conservation of the nature of the interactions between the two species, which is not entirely guaranteed by many modifications tested. As shown in our work, the main entropic effects associated with such changes may potentially be well captured using models composed of the first layer residues in a pocket. Another point of relevance is how the chemical changes affect the interactions with the solvent of the free ligand and of/on the complex. Here the situation might be more delicate since in most cases we introduced H-Bond acceptors or charged groups. Particularly the sulfoxide and the carbonyl/carboxyl modifications could be potentially problematic, and in such cases, entropy corrections are strictly necessary.

**S18 – Analysis of protonation states of compound 1**

b

a

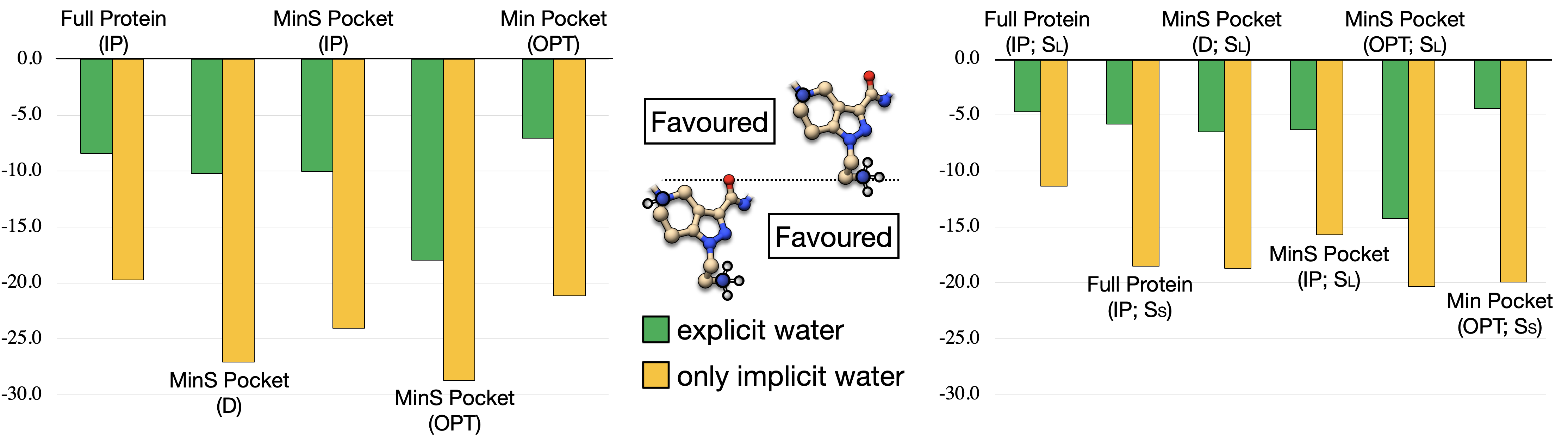

**Figure S18.1. Comparison of protonation states of compound 1 at a) the binding energy level and b) at the Gibbs free energy level, using implicit waters and eventually also explicit ones.** IP stands for in-pocket optimization, OPT for unconstrained optimization, and D is the direct calculation, using a crystal with protons added by Chimera. The MinS pocket consists of the first row of amino acids interacting with the ligand with all their side chains. The smaller pocket Min retains only those side chains that are directed to the ligand ($\lesssim$ 8 Å). Enthalpy and entropy corrections were calculated from both pocket constructs after full optimization. This is marked with S_L_ or S_S_ in the plots, corresponding to either MinS or Min pockets.

We also used the calculations to analyze the preferential protonation state of ligand **1** in the pocket of PEX14 (Figures S18.1a,b). Irrespective of the system used, the doubly protonated state has the most favorable interaction energy. As the free protein state, we use to calculate binding data is always the same, we conclude that the observed effect results from the additional stabilization arising in the complex. Consequently, one should expect the ligand in its doubly protonated state when binding takes place. This might however be the case only for the ligand here studied, due to the specificities of the water envelope shown in the complex. Conformational entropies were also estimated for the ligands here studied. We noticed however there is no advantage in including these for the calculations, most likely due to an imbalance in the model. For more details, refer to the supplemental S16.

**S19 – Conformational entropy for compounds 1 and 14 in two protonation states**

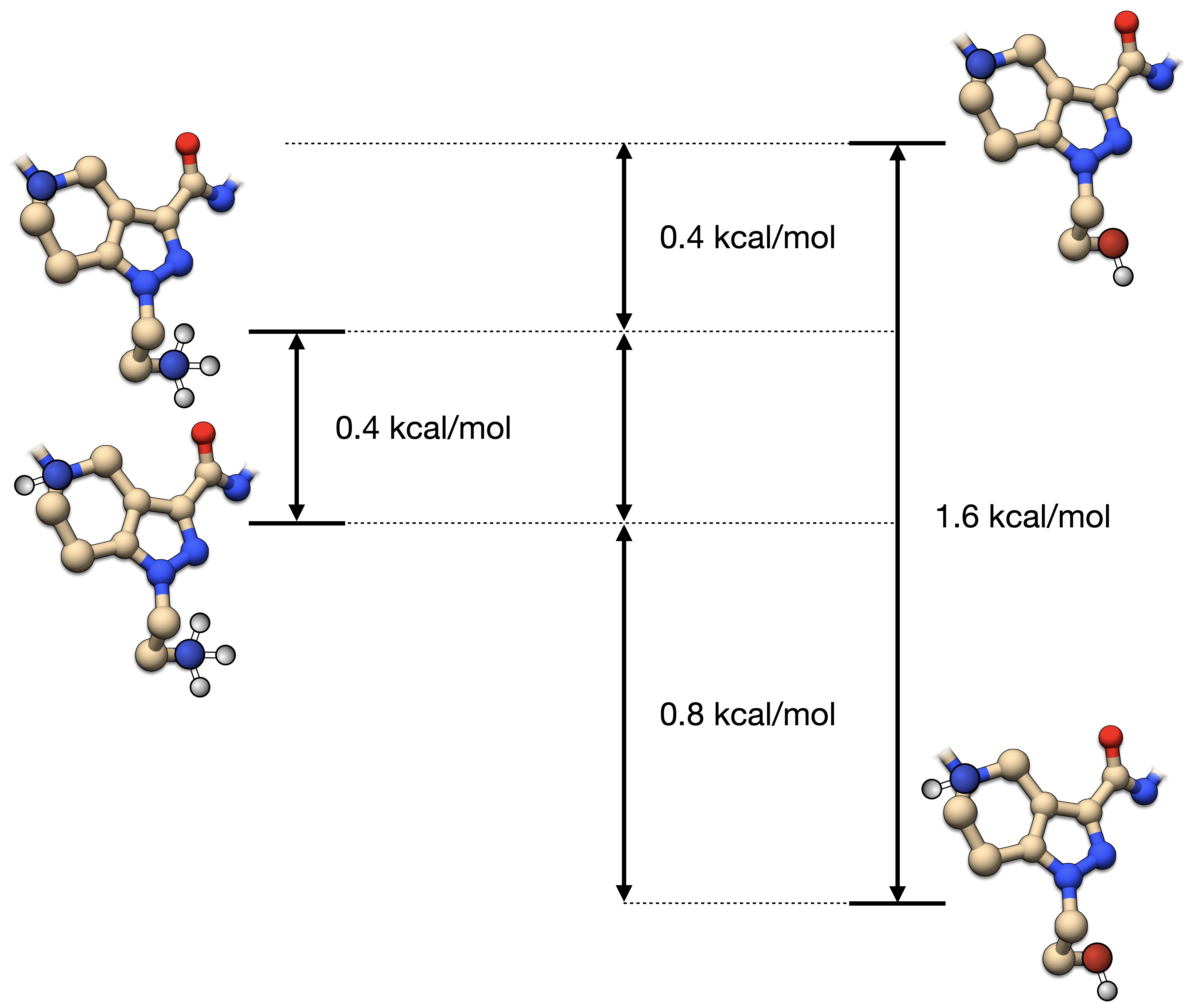

**Figure S19.1:** Conformational entropy for compounds **1** and **4** in two protonation states. In both cases, increasing the pH is expected to increase the conformational entropy of the free ligand by 0.4-1.6 kcal/mol.

**S20 – Recorded NMR spectra of target compound 1f and its precursors 17-21.**

^1^H NMR spectrum of compound **17** recorded on a Bruker AV-HD400 spectrometer (400 MHz, CDCl_3_):

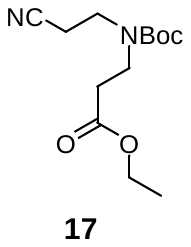

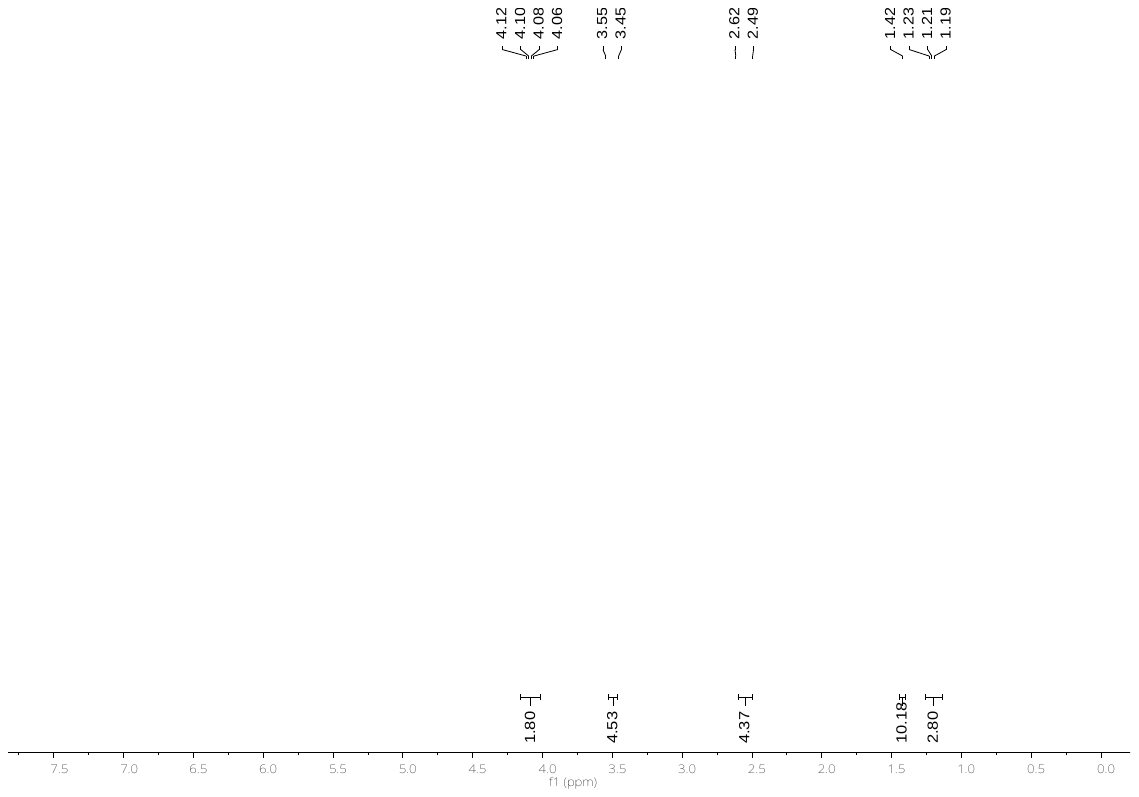

^13^C NMR spectrum of compound **17** recorded on a Bruker AV-HD400 spectrometer (101 MHz, CDCl_3_):

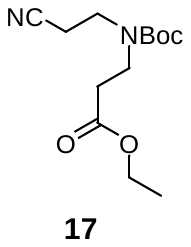

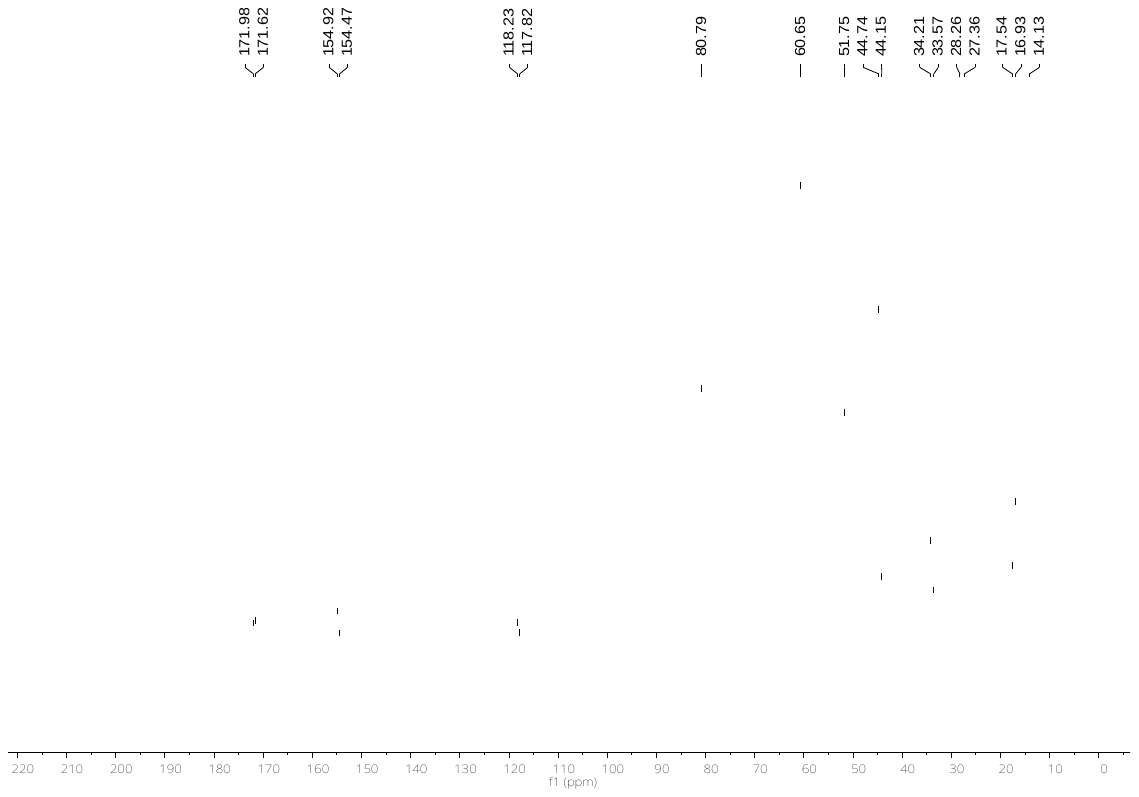

^1^H NMR spectrum of compound **18** recorded on a Bruker AV-HD400 spectrometer (400 MHz, DMSO‑d6):

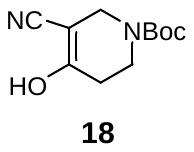

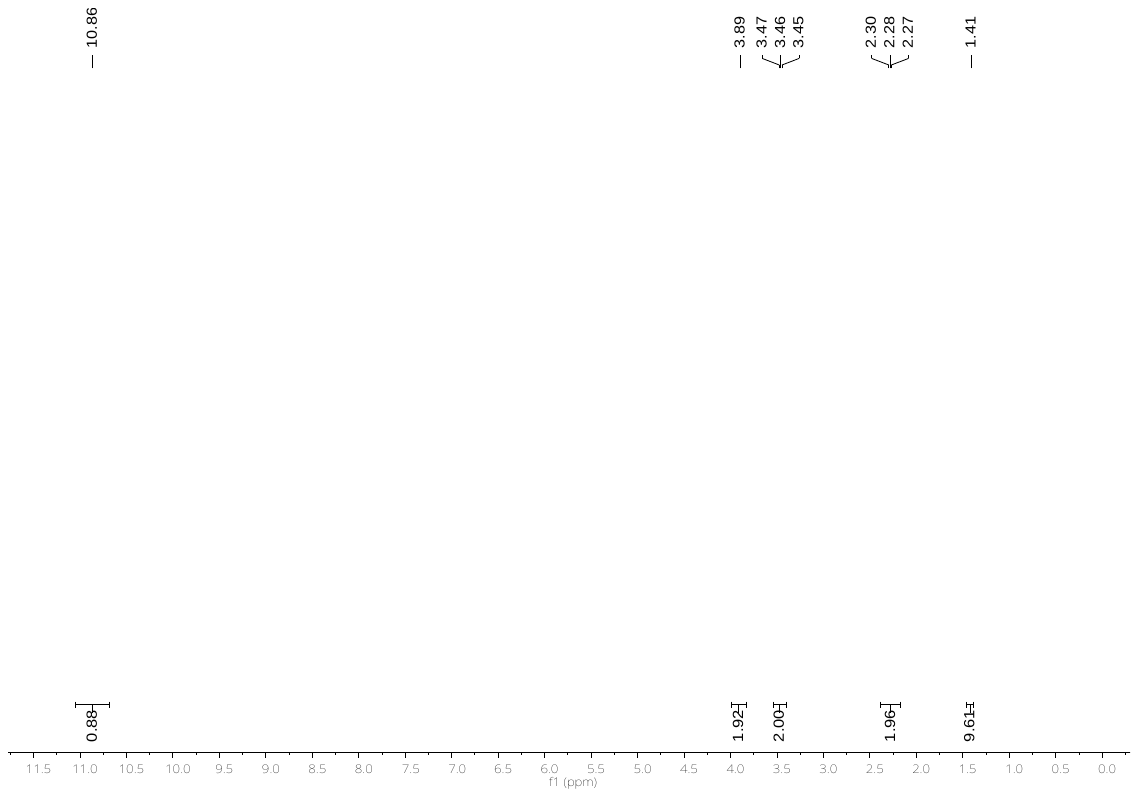

^13^C NMR spectrum of compound **18** recorded on a Bruker AV-HD400 spectrometer (101 MHz, CDCl_3_):

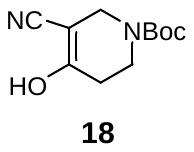

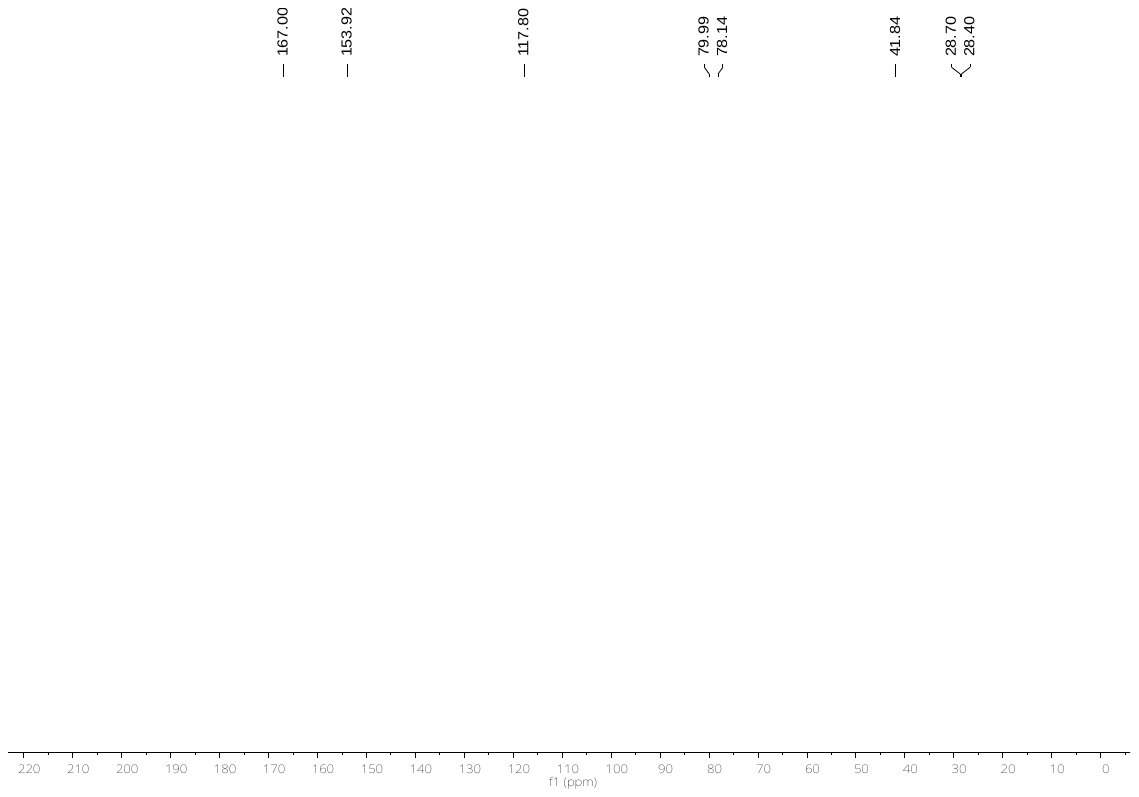

^1^H NMR spectrum of compound **19** recorded on a Bruker AV-HD400 spectrometer (400 MHz, CDCl_3_):

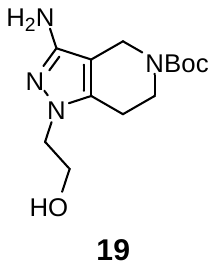

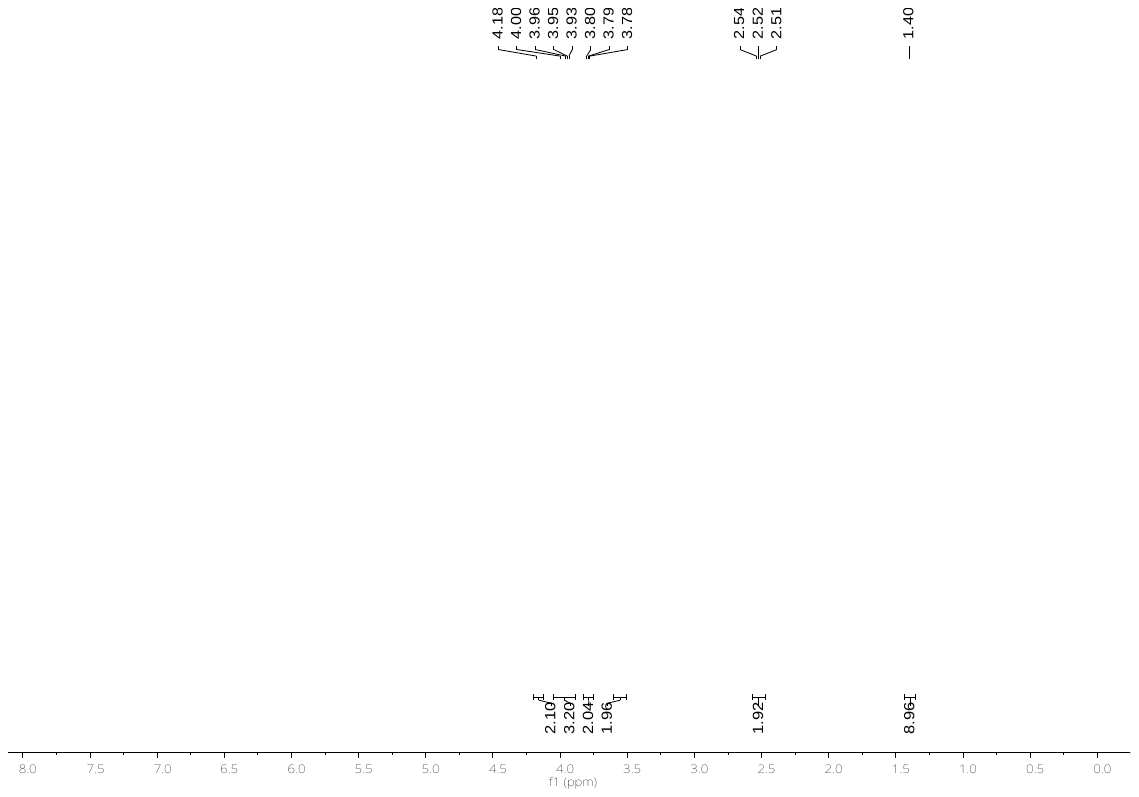

^13^C NMR spectrum of compound **19** recorded on a Bruker AV-HD400 spectrometer (101 MHz, CDCl_3_):

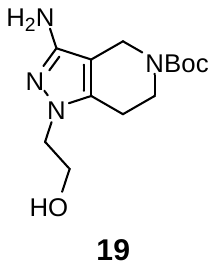

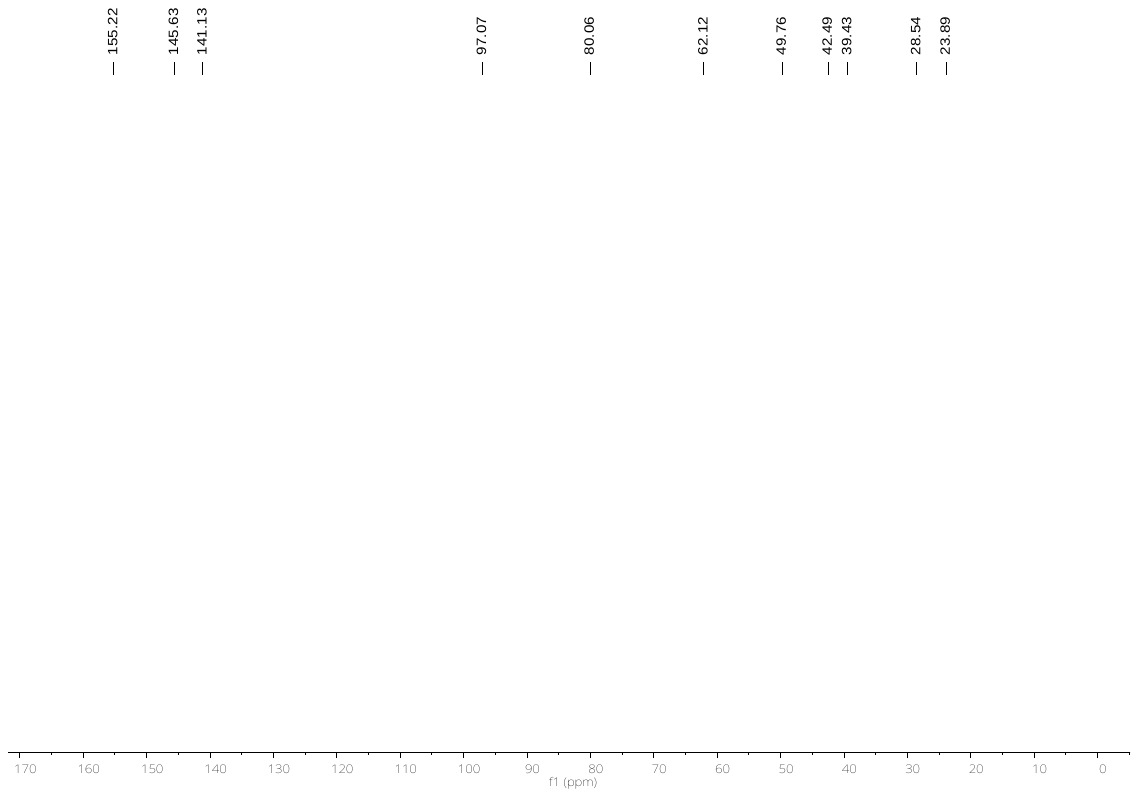

^1^H NMR spectrum of compound **20** recorded on a Bruker AV-HD400 spectrometer (400 MHz, CDCl_3_):
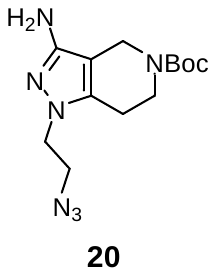

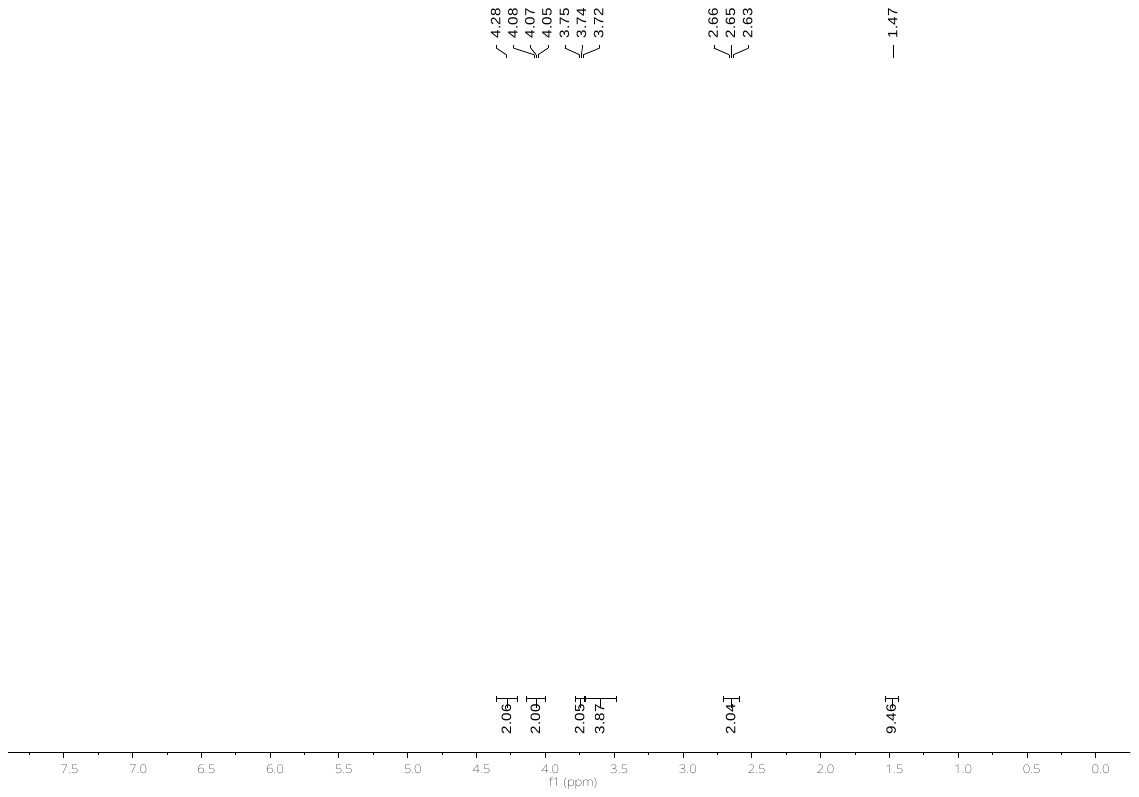

^13^C NMR spectrum of compound **20** recorded on a Bruker AV-HD400 spectrometer (101 MHz, CDCl_3_):

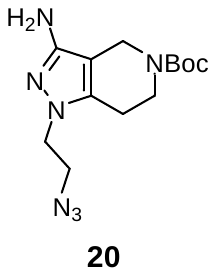

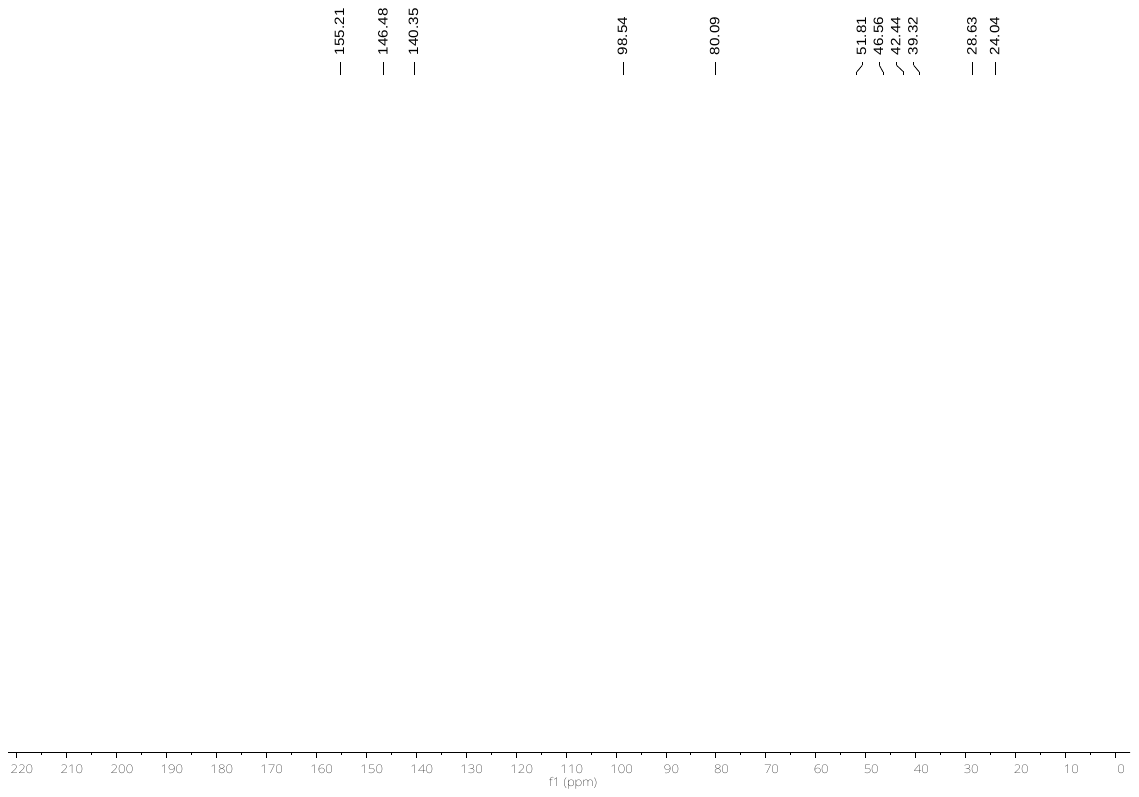

^1^H NMR spectrum of compound **21** recorded on a Bruker AV-HD400 spectrometer (400 MHz, CDCl_3_):

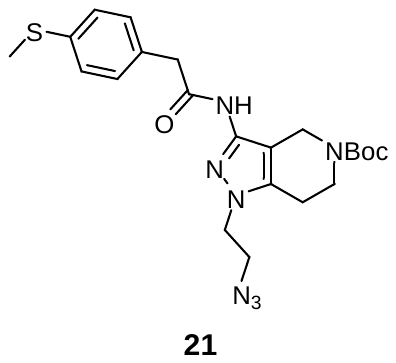

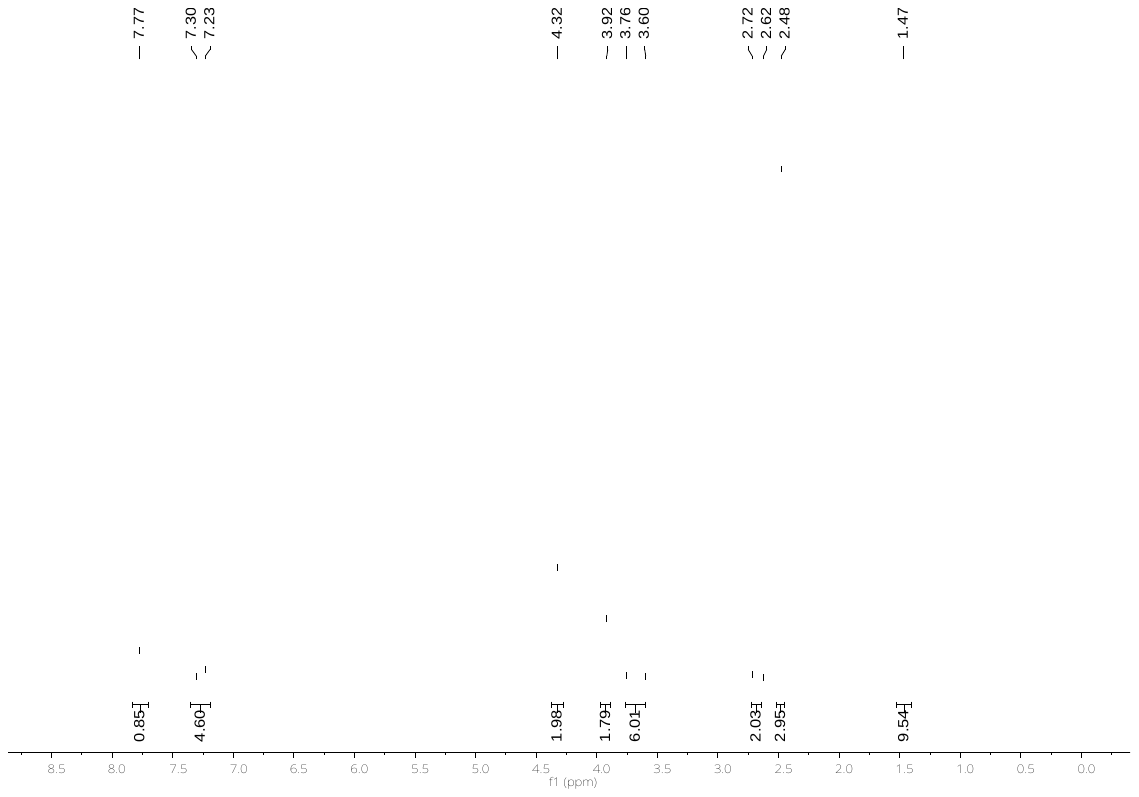

^13^C NMR spectrum of compound **21** recorded on a Bruker AV-HD400 spectrometer (101 MHz, CDCl_3_):

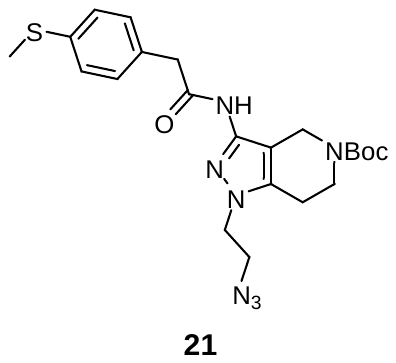

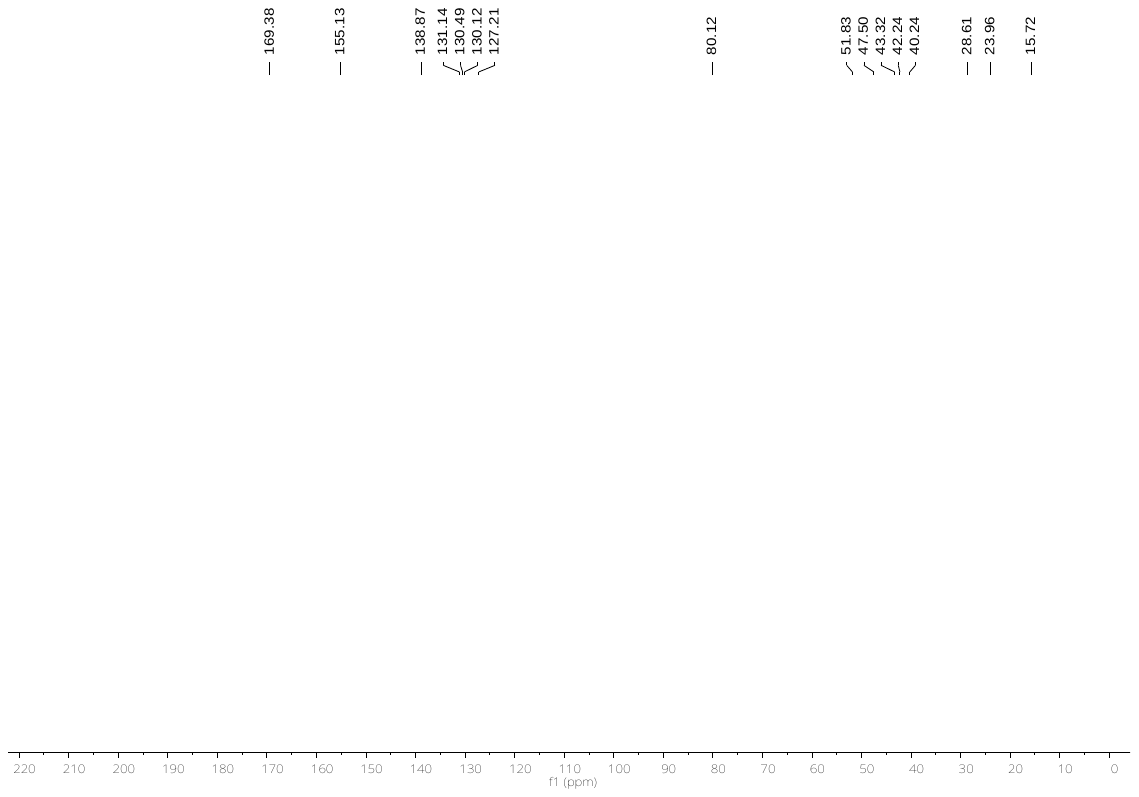

^1^H NMR spectrum of compound **1f** recorded on a Bruker AV-HD400 spectrometer (400 MHz, CD_3_OD):

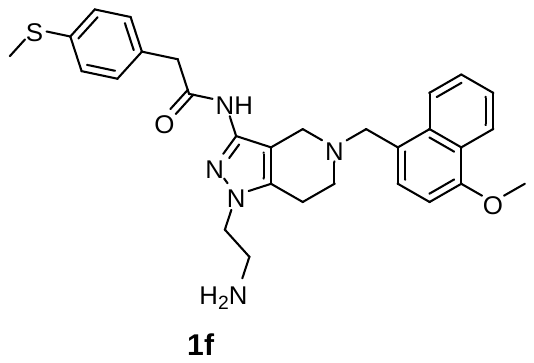

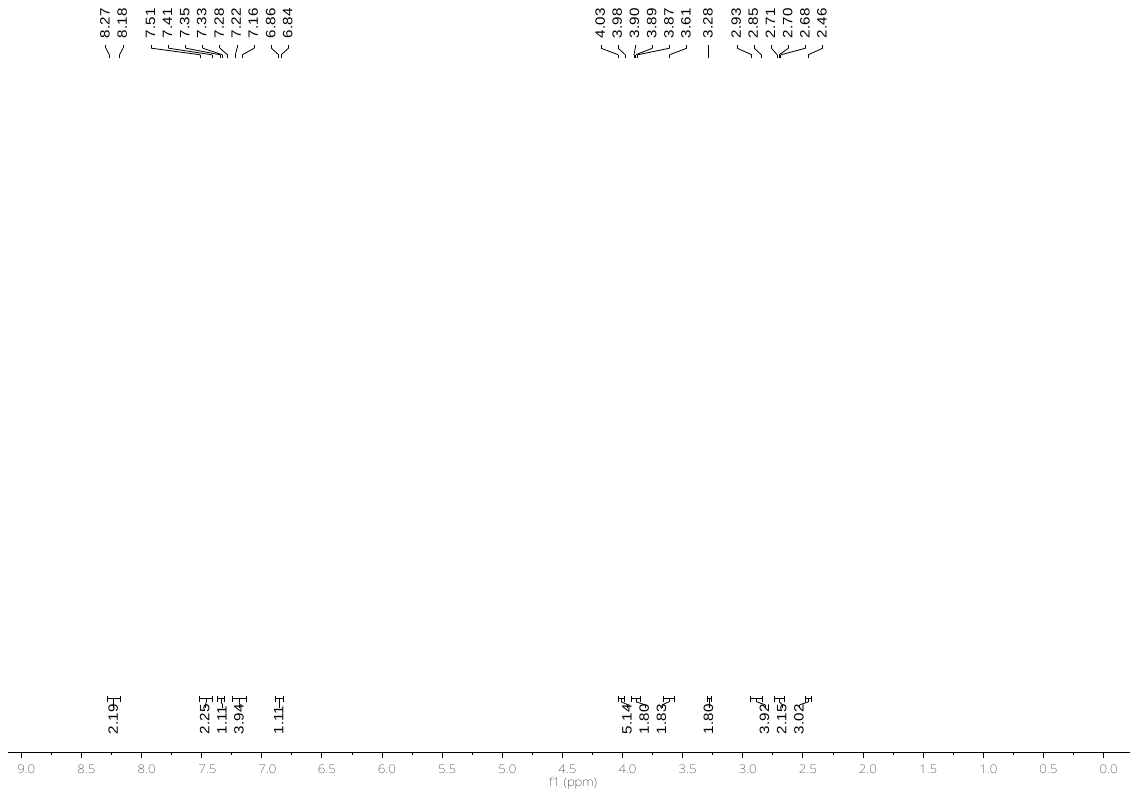

^13^C NMR spectrum of compound **1f** recorded on a Bruker AV-HD400 spectrometer (101 MHz, CD_3_OD):

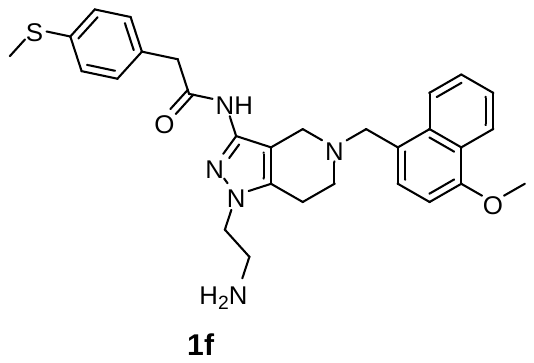

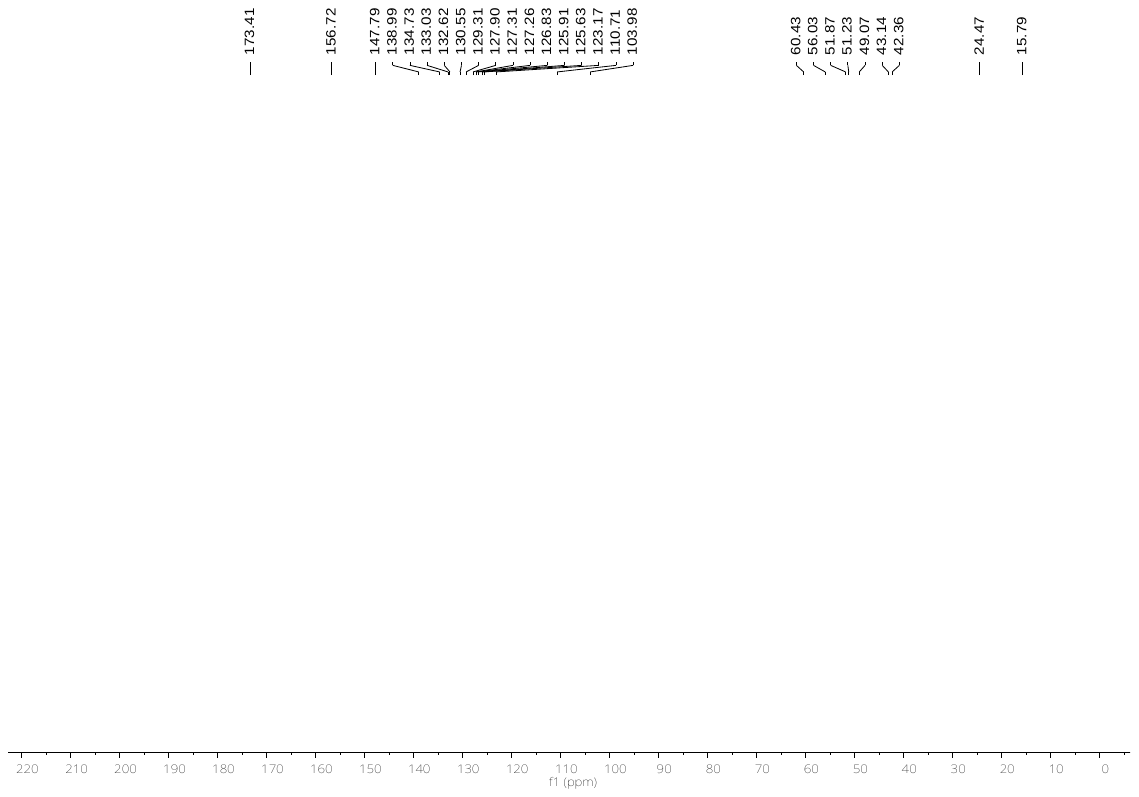
